# Scaling deep identifiable models enables zero-shot characterization of single-cell biological states

**DOI:** 10.1101/2023.11.11.566161

**Authors:** Mingze Dong, Kriti Agrawal, Rong Fan, Esen Sefik, Richard A. Flavell, Yuval Kluger

**Affiliations:** Interdepartmental Program in Computational Biology & Bioinformatics, Yale University, New Haven, CT, USA; Department of Pathology, Yale School of Medicine, New Haven, CT, USA; Department of Biomedical Engineering, Yale University, New Haven, CT, USA; Department of Immunobiology, Yale University, New Haven, CT, USA; Yale Stem Cell Center and Yale Cancer Center, Yale University, New Haven, CT, USA; Human and Translational Immunology, Yale University, New Haven, CT, USA; Howard Hughes Medical Institute, Yale University, New Haven, CT, USA; Applied Mathematics Program, Yale University, New Haven, CT, USA

## Abstract

How to identify true biological differences across samples while overcoming batch effects has been a persistent challenge in single-cell RNA-seq data analysis, hindering analyses across datasets for transferable biological findings. In this work, we show that scaling up deep identifiable models leads to a surprisingly effective solution for this challenging task. We developed scShift, a deep variational inference framework with theoretical support in disentangling batch-dependent and independent variations. By training the model with compendiums of scRNA-seq atlases, scShift shows remarkable **zero-shot** capabilities in revealing representations of cell types and biological states in single-cell data while overcoming batch effects. We employed scShift to systematically compare lung fibrosis states across different datasets, tissues and experimental systems. scShift uniquely extrapolates lung fibrosis states to previously unseen post-COVID-19 fibrosis, characterizing universal myeloid-fibrosis signatures, potential repurposing drug targets and fibrosis-associated cell interactions. Evaluations of over 200 trained scShift models demonstrate emergent zero-shot capabilities and a scaling law beyond a transition threshold, with respect to dataset diversity. With its scaling performance on massive single-cell compendiums and exceptional zero-shot capabilities, scShift represents an important advance toward next-generation computational models for single-cell analysis.

## Main

With advancements in single-cell technologies, emerging studies have generated millions of single-cell RNA measurements across different cell types and biological states. The incorporation of large-scale scRNA-seq datasets into computational models has been accelerated by consortia such as Tabula Sapiens [1], Human Cell Atlas (HCA) [2], and Human Biomolecular Atlas Program (HuBMAP) [3] that collect massive sequencing data, along with data standardization and curation efforts such as Chan-Zuckerberg CellxGene census [4], Azimuth [5], and gget [6]. With the rapid accumulation of large-scale scRNA atlas datasets, numerous computational methods have been proposed to integrate these datasets [7–22]. These works mostly aim to learn representations that reveal cell type organization while mitigating batch effects (i.e., measurement differences resulting from different technologies, labs, etc.).

Beyond identifying shared cell types, a critical next step in revealing novel biology is to compare different biological states (such as disease conditions, perturbations, and treatments) across datasets. However, a fundamental challenge in the field—distinguishing batch effects from true biological differences—has made this task particularly challenging. This challenge represents a classic case of non-identifiability, a well-documented problem in statistics [23]. Establishing identifiability in nonlinear models, particularly neural networks, generally requires comprehensive annotations [24], which are almost impossible to obtain in real datasets. The non-identifiable nature of batch effects versus biological variation presents a conceptual, rather than technical, barrier—one that cannot be overcome simply by enhancing model architectures, expanding training datasets, or optimizing parameter tuning.

Our key motivation for this work is that, while comprehensive annotations of biological states are mostly infeasible, we can leverage dataset labels as supervision to identify batch-dependent variations (Supplementary Note). The batch-dependent variations comprise both biological states and batch effects. Within each individual dataset (batch), these variations effectively represent the biological differences we wish to study. Under appropriate assumptions, we can directly compare how these biological differences change (for example, from normal to disease states) across multiple independent studies, overcoming batch effects between them. This framework enables leveraging existing compendiums of scRNA-seq atlases to build unbiased feature extractors of biological states **without the need of external annotation**. The model’s theoretical capacity to identify distinct biological states scales with the number of biologically diverse datasets used for training.

Building on this insight, we developed scShift, a novel variational inference framework that deconvolves batch-dependent and independent variations in single-cell data. We established theoretical support for model identifiability (Supplementary Note) and incorporated essential identifiability designs into the scShift architecture. When pretrained on comprehensive scRNA-seq datasets from blood and lung tissues, scShift demonstrated remarkable zero-shot capability in separating cell type signatures (batch-independent variation) and biological state signals (batch-dependent variation). scShift also exhibited substantially improved power for classification of disease states in out-of-distribution (OOD) settings. Specifically, scShift uniquely characterized unseen post-COVID-fibrosis states using lung fibrosis data and identified universal myeloid-fibrosis signatures, potential drug repurposing targets and fibrosis-associated cell interaction network. The fibrosis state characterized by scShift was further validated by scRNA-seq measurements from chronic COVID-19 humanized mouse models. Systematic evaluation of over 200 scShift models under varying training conditions unveiled emergent zero-shot capabilities with respect to donor numbers in the training set. Moreover, a scaling law of zero-shot capabilities with respect to donor and cell numbers was observed beyond a transition threshold.

## Results

### scShift disentangles batch-dependent and independent variations to reveal biological states

In this work, we model the high-dimensional gene expression distribution *p*(*x*_*i*_) for each cell *i* using two sets of low-dimensional latent variables *z*_*i*_, *s*_*i*_. *z*_*i*_ represents intrinsic cellular properties (such as cell types) shared across datasets, whereas *s*_*i*_ encodes both biological states and batch effects, thus should be different across datasets (Fig. 1a). Our approach is notably different from most previous representation learning methods [7, 9, 20, 25–27] for single-cell data, which model gene expression by concatenations (instead of summations) of low-dimensional representations and batch label encodings.

**Fig 1.**
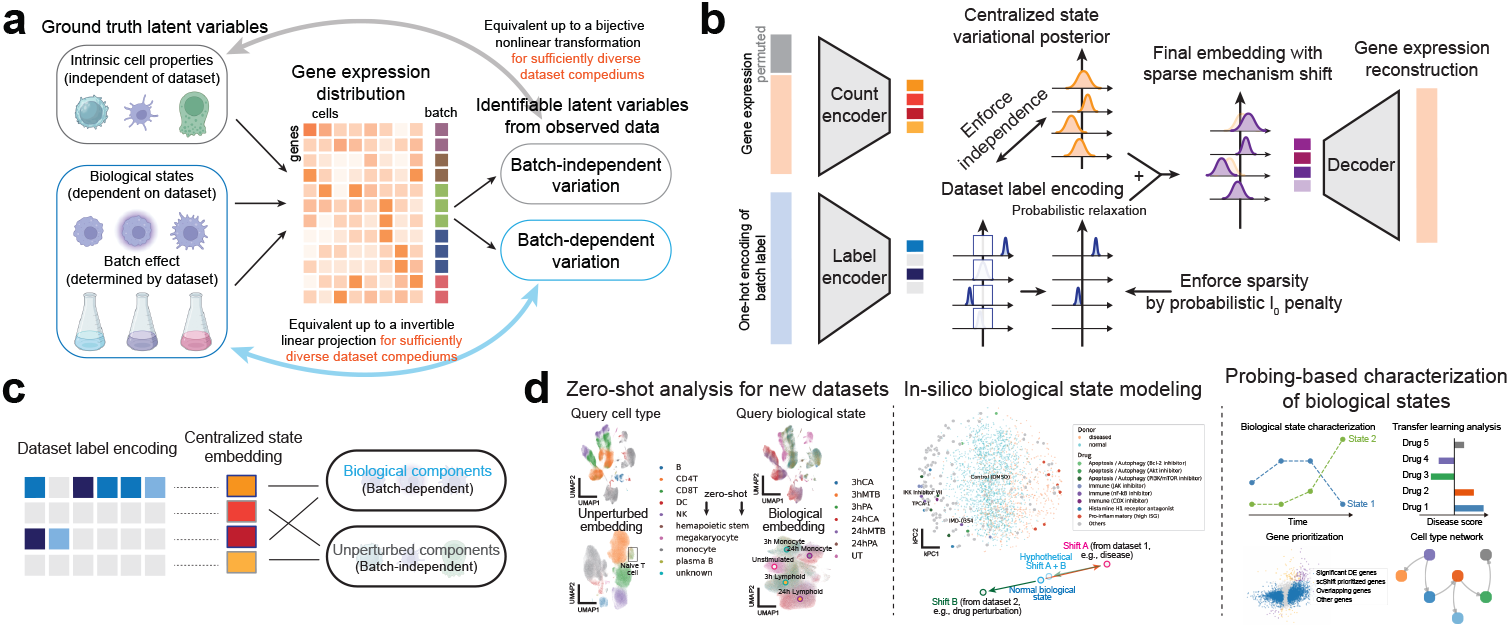
Overview of the scShift framework. **a**. The conceptual foundations of scShift, including the generative model and the theoretical identifiability guarantee. **b**. The workflow of scShift. **c**. Variation decomposition strategy in scShift. The centralized state is separated into biological and unperturbed components using dataset label encoding. **d**. Key applications and use cases of scShift.

The key advantage of our model lies in identifiability. In approaches that model cell representations and batch encodings separately, biological differences across samples or datasets are arbitrarily entangled with cell type or subtype information, due to the well-known non-identifiability result for nonlinear ICA [23]. This not only prevents the extraction of biological states but also precludes meaningful comparisons of biological states across datasets with different batch or baseline conditions (Supplementary Note). For instance, disease versus normal representations observed *in vivo* may not be comparable to drug perturbation datasets conducted *in vitro* due to differences in culture conditions.

In contrast, our approach approximates the biological state *s* and cell type *z* representations through identifiable batch-dependent and batch-independent variations, respectively. This approximation holds when sufficient biologically distinct datasets are available to span all possible biological variations. In practice, we expect the batch-dependent representation to capture the majority of possible biological states when training on extensive single-cell atlas collections. Through additional supervision from the dataset label, we derived theoretical support for model identifiability in this task under mild assumptions. In particular, our theoretical results demonstrate that batch-dependent variations are linearly identifiable from data, enabling comparisons of biological state shifts across datasets (Fig. 1a, Supplementary Note).

Our theoretical framework identifies two key requirements for successful disentanglement: first, batch-dependent components should be the sparsest among possible realizations, and second, batch labels should be statistically independent from the latent variables after centralization. Here, the centralized latent variables refer to the full (batch-dependent and independent) variation after removing the batch-specific term (see Methods and Supplementary Note). Consequently, we designed the scShift model architecture as illustrated in Fig. 1b. scShift consists of two encoders for the centralized latent variables and dataset label, respectively. The outputs of these encoders are added to yield the full variation, which is then used to reconstruct the gene expression distribution (Fig. 1b, Methods). We enforce sparsity in the dataset label encoding using probabilistic *l*_0_ regularization, building upon the stochastic gate approach [28–30] (See Methods). We use kernel maximum mean discrepancy (MMD) regularization [31] to enforce independence between the centralized latent variables and the dataset label encoding. To enhance model generalization, we introduce noise during training by randomly permuting a small subset of genes (25%) across cells within each mini-batch (Fig. 1b). The parameters of the scShift model are optimized by maximizing the evidence lower bound (ELBO) [25, 32] together with sparsity and independence regularizations.

After training, we are able to decompose the full centralized state into two parts. The batch-dependent part, which we term the *biological embedding*, corresponds to the components associated with any non-zero entries in the dataset label encoding. The batch-independent part, which we term the *unperturbed embedding*, comprises the remaining components with all-zero dataset label encoding (Fig. 1c). Since dataset label encoding is the only model component requiring batch labels, both biological and unperturbed embeddings can be extracted from new datasets in a zero-shot manner without additional training.

To our knowledge, scShift is the first model demonstrating the capability to reveal biological state representations without annotations of these states. As such, it can be effectively trained on existing compendiums of single-cell atlases (such as CellXGene [4]) and enables various novel analyses for scRNA-seq datasets. First, scShift automatically reveals intrinsic representations and biological states for new query datasets in a zero-shot manner, powering downstream analysis. Second, scShift enables unsupervised comparison of biological state shifts across datasets, which we term *in silico* modeling of biological states. For instance, *in vitro* perturbation conditions can be directly compared to specific disease conditions from patient samples collected *in vivo* using scShift (Fig. 1d).

Finally, scShift enables exploration of biological-state-specific or disease-specific representations, when a certain condition is of interest. In this case, we can construct classifiers of scShift embeddings for the biological state based on existing datasets (see Methods), analogous to probing classifiers in vision or language models [33, 34]. Through scShift’s capacity for cross-dataset comparisons, these classifiers provide projections of specific biological states that generalize to hold-out datasets, enabling characterization of disease-specific cellular states, key genes, potential therapeutic targets, and cellular interactions, ultimately facilitating the creation of unified disease-centric atlases integrating millions of cells from all existing datasets (Fig. 1d).

### scShift dissects cellular-level biological states in human blood

As a proof-of-concept, we first trained the scShift model on our assembled human blood scRNA-seq data compendium. It includes 1,000,000 cells from 30 studies and 2,538 donors subsampled from the CellXGene blood compendium [4], plus 240,090 cells from 144 drug perturbations and 3 negative/positive control drug conditions [35] (Fig. 2a, Methods). After training, the scShift unperturbed embedding of the training compendium preserves cell type information while eliminating evident batch effects (Fig. 2b, Supplementary Fig. 1). Combining the biological and unperturbed embeddings reveals additional disease or perturbation-specific clusters (Fig. 2c). As expected, scShift does not outperform alternative methods (Harmony, scVI, scANVI, scPoli [7–9, 20]) in scIB atlas integration benchmarks [18] on the training set, since these tasks do not involve correct specification of biological differences or zero-shot capabilities (Extended Data Fig. 1a, Supplementary Methods). Nevertheless, the benchmarking results indicate successful disentanglement, attributing cell type information to the unperturbed embedding while removing this information from the biological embedding (Extended Data Fig. 1a, Supplementary Fig. 1).

**Fig 2.**
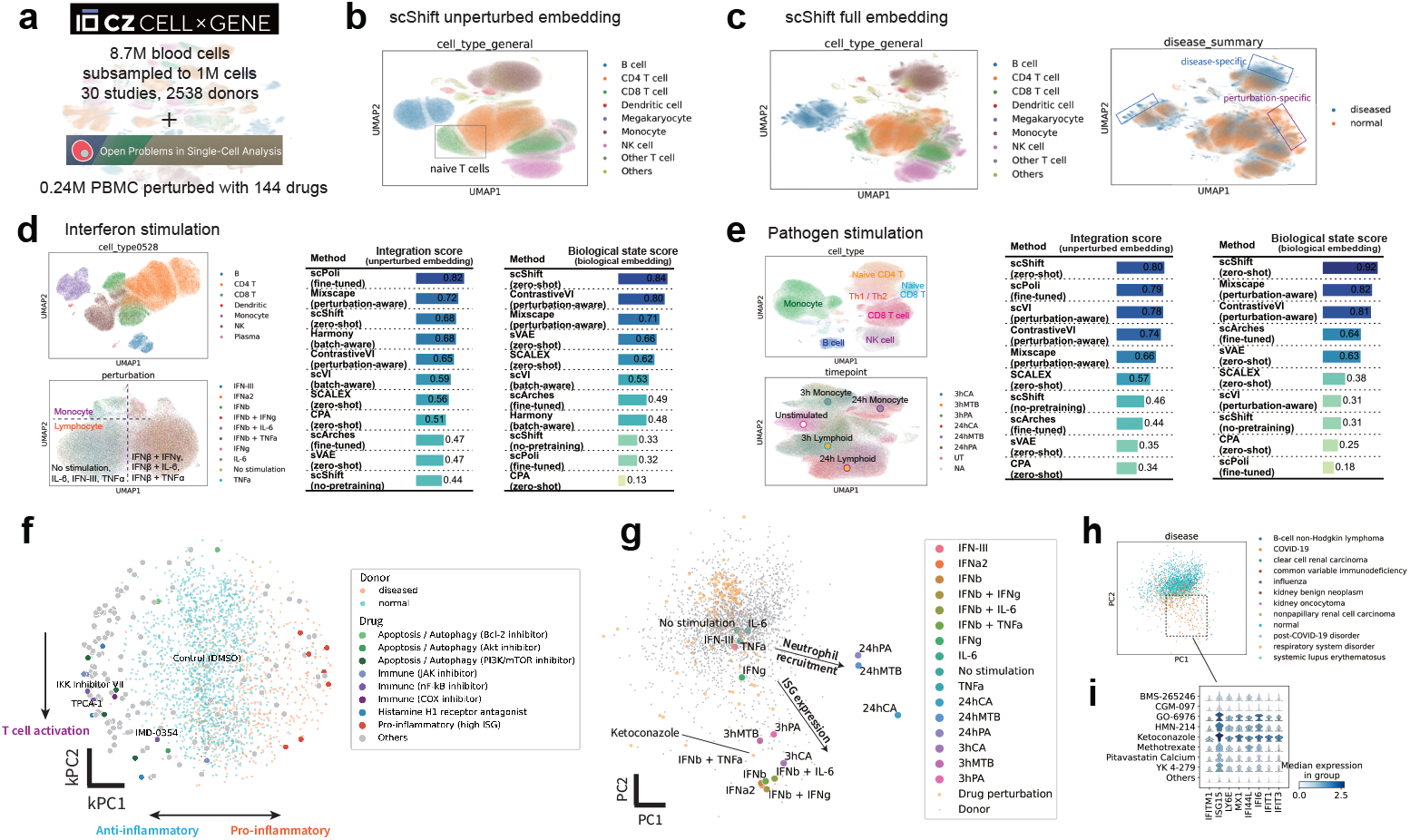
Training and validation of scShift on blood data. **a**. Overview of the training compendium for the scShift blood model. The training set comprises subsampled 1 million blood cells from the CellxGene atlas [4], and the open-source drug perturbation data including 144 drug perturbations in 0.24 million blood cells [35]. **b**. UMAP visualization of the training compendium using scShift unperturbed embedding, colored by cell type (left), sequencing assay (middle) and disease state (right). **c**. UMAP visualization of the training compendium using scShift full embedding after adjustment (Methods), colored by cell type (left) and disease state (right). The UMAP1 axis was flipped for better concordance with Fig. 2b. **d**. UMAP visualizations of the interferon stimulation dataset using scShift unperturbed/biological embedding colored by cell type/condition (left); scIB rescaled overall score [18] for cell type and condition representation evaluations across methods on the interferon stimulation dataset (right). Integration score evaluates cell type preservation and perturbation/batch removal. Biological state score evaluates perturbation preservation and removal of selected cell type information (see Methods). **e**. UMAP visualizations of the pathogen stimulation dataset using scShift unperturbed/biological embedding colored by cell type/condition (left); scIB rescaled overall score [18] for cell type and condition representation evaluations across methods on the pathogen stimulation dataset (right). Integration score evaluates cell type preservation and perturbation removal. Biological state score evaluates perturbation preservation and removal of selected cell type information (see Methods). **f**. Combined visualization of CD4 T cell pseudobulk profiles across donor samples and drug perturbations, colored by the donor disease state or drug function respectively. **g**. Combined visualization of CD4 T cell pseudobulk profiles across datasets, colored by perturbation type or data source (see Text and Supplementary Methods for f-g). **h**. Visualization of donor samples from **g** colored by disease type. **i**. Stacked violin plot for log-normalized interferon stimulated gene (ISG) expressions among drugs in the high ISG wing versus other drugs.

We further assessed various methods’ performance on hold-out datasets unseen during training. This evaluation examined each model’s ability to detect cell types and biological states across zero-shot, fine-tuning, and training-from-scratch settings. For assessment, we used two public datasets of cultured blood cells with well-characterized biological states: one containing 9 distinct interferon stimulation conditions and another with 6 different pathogen exposure conditions, plus control conditions [36, 37] (Extended Data Fig. 1). Our benchmarking included three categories of methods: batch correction methods requiring batch/perturbation label inputs (Harmony, scVI [7, 8]), perturbation modeling approaches requiring control/non-control labels and optimized for the task (Mixscape, contrastiveVI [38, 39]), and transfer learning approaches in either fine-tuned (scArches, scPoli [19, 20]) or zero-shot settings (CPA, sVAE [40, 41]).

Our results suggest that in both datasets, scShift achieved top performance in identifying accurate cell type representations and outperformed all other methods in inferring accurate biological states by a notable margin (Fig. 2d-e, Supplementary Figs. 3-4). scShift demonstrated superior capability in detecting fine-grained cell subtypes within the pathogen stimulation dataset, outperforming all comparative methods in quantitative assessments (Fig. 2e, Supplementary Fig. 4). The distinct clustering of cell subtypes (particularly naive CD8 T cells) observed in the pathogen stimulation dataset may explain the separated CD8 T clusters seen in the interferon stimulation dataset, and scShift’s slightly lower cell type preservation score in this dataset (Fig. 2d, Supplementary Fig. 3). In terms of biological states, scShift demonstrated superior performance as a zero-shot method, outperforming even optimized condition-aware approaches (Mixscape, ContrastiveVI), while substantially exceeding all other methods. In the interferon stimulation dataset, scShift revealed distinct response patterns of monocytes and converging response patterns of lymphocytes (CD4 T cells, CD8 T cells, NK cells), consistent with original findings using a supervised approach [36] (Fig. 2d). Ablation studies demonstrated the importance of all key scShift design components (sparse encoding, denoising training scheme, and MMD regularization) for achieving accurate zero-shot inference (Supplementary Fig. 5 and Methods).

**Fig 3.**
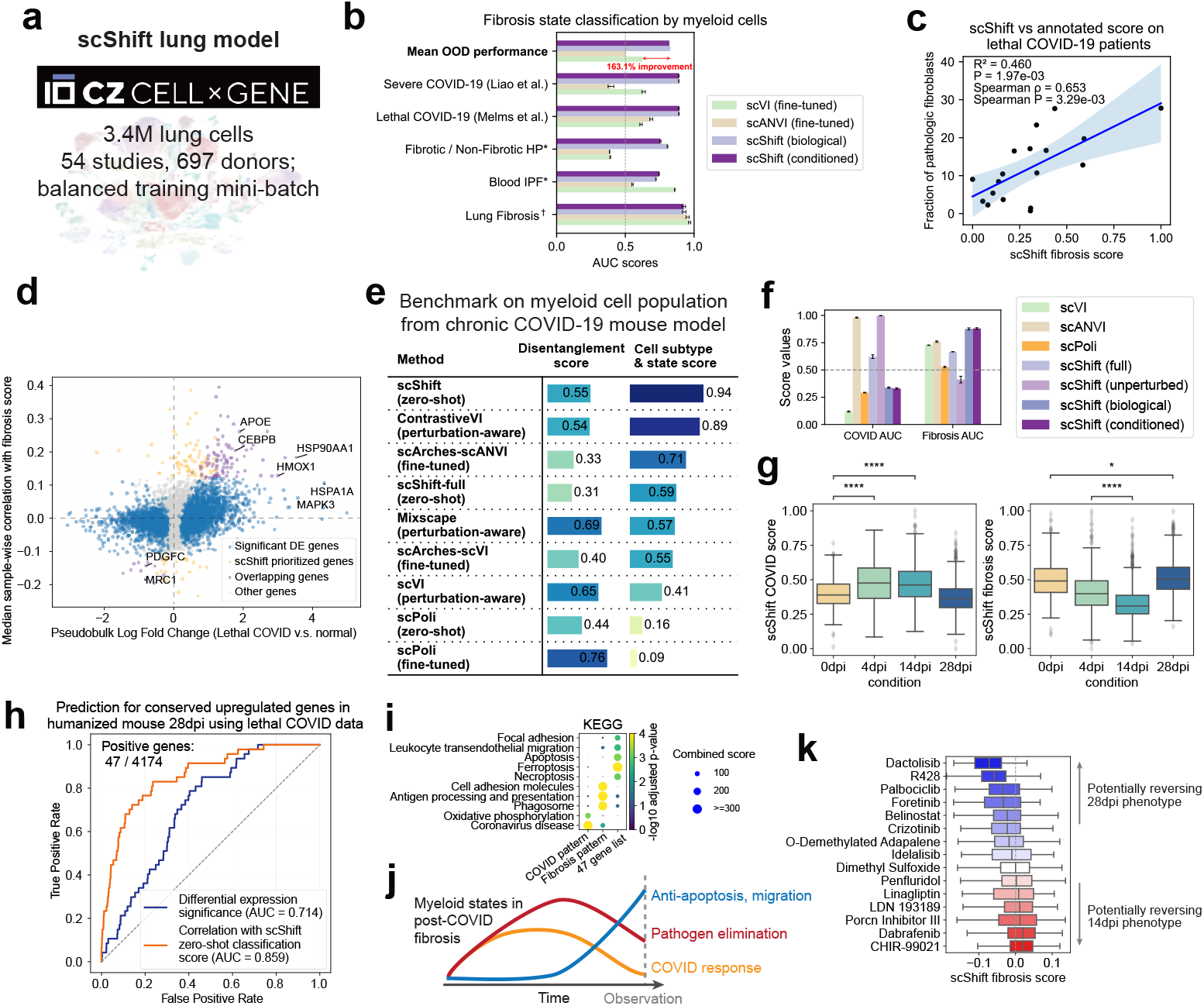
scShift characterizes universal myeloid-fibrosis states in lung. **a**. Overview of the CellxGene training compendium for scShift lung model. **b**. Barplot showing performances of different methods in predicting fibrosis / disease states using myeloid cells across 5 datasets. Bars represent mean Area Under the Curve (AUC) values from 10 independent experiments with different random seeds, with error bars indicating standard error. *†* denotes evaluations performed using hold-out patients within the same dataset, while other evaluations represent out-of-distribution (OOD) performance on hold-out datasets. *\** indicates data collected from blood samples, in contrast to other lung datasets. **c**. Scatterplot showing the relationship between scShift fibrosis scores and pathologic fibroblast proportions in lung tissue from lethal COVID-19 patients [48]. **d**. Scatterplot showing pseudobulk log fold change (lethal COVID-19 vs. healthy donors) and the scShift correlation score for each gene (n=21844). The scShift correlation scores were defined as median sample-wise correlations with scShift fibrosis scores in lethal COVID-19 patients (see Methods). **e**. scIB rescaled average scores evaluating disentanglement and cell subtype/condition preservation performances on the myeloid cells from the humanized mouse dataset [53]. Disentanglement score was defined as the average of condition and cell subtype removal scores. Cell subtype & state score was defined as the average of cell subtype and condition preservation scores (see Supplementary Methods). **f**. Barplot showing performances of different methods in predicting fibrosis and COVID-19 states in the humanized mouse dataset [53]. Bars represent mean Area Under the Curve (AUC) values from 10 independent experiments with different random seeds, with error bars indicating standard error. **g**. Boxplot showing single-cell COVID/fibrosis scores for myeloid cells in the humanized mouse dataset. n=381,275,752,1209. Statistical significance were determined using one-sided Mann- Whitney-Wilcoxon test. ns: 5.00e-02*<*p≤1.00e+00; *: 1.00e-02*<*p≤5.00e-02; **: 1.00e-03*<*p ≤1.00e-02; ***: 1.00e-04*<*p ≤1.00e-03; ****: p≤1.00e-04. **h**. ROC curves and AUC values for pseudobulk differential expression significance (lethal COVID-19 versus control) and scShift correlation scores (derived from lethal COVID-19 patients, see Methods) in distinguishing consistently upregulated genes (n=47) across lethal COVID-19 patients and humanized mouse 28dpi. **i**. Dotplot showing KEGG pathway enrichment analysis for genes of COVID pattern, fibrosis pattern, and the identified 47 gene list. **j**. Illustration of myeloid gene expression temporal dynamics in post- COVID fibrosis. **k**. Boxplot showing fibrosis scores of drug candidates from [35].

**Fig 4.**
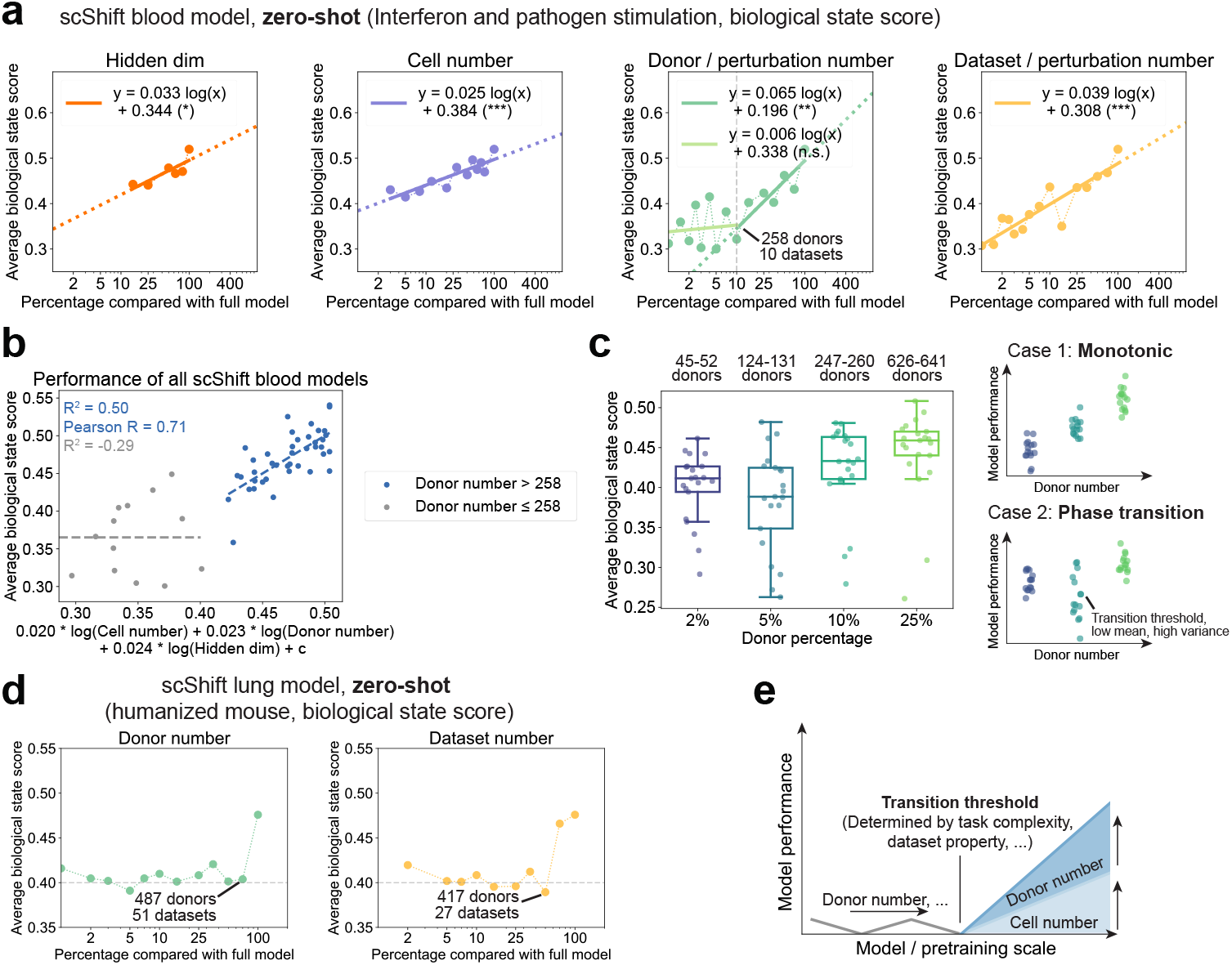
scShift exhibits emergent and scaling behaviors in zero-shot tasks with respect to dataset diversity. **a**. Scatterplots of average biological state scores (average scIB scores of condition preservation and cell type removal on interferon and pathogen stimulation datasets, see Methods) across biological embeddings of scShift blood models trained under different settings. **b**. Scatterplot showing the empirically fitted scaling function and the average biological state score in scShift blood models. The linear regression *R*^2^ and Pearson R for models with donor numbers *>* 258 are shown in blue. The linear regression *R*^2^ between the empirically fitted scaling function and the average biological scores for models with donor numbers ≤258 is shown in gray. **c**. Boxplots showing average biological scores of scShift models when donors/perturbations are subsampled to 2%, 5%, 10%, and 25% of the full dataset (n=21 for each setting, left); Illustrations of model behavior under two hypotheses (right). **d**. Scatterplots of average biological state scores (average scIB scores of condition preservation and cell type removal for myeloid cells from the COVID-19 humanized mouse dataset [53], see Methods) across biological embeddings of scShift lung models trained under different settings. **e**. A schematic overview illustrating key scaling properties of scShift with a fixed model architecture.

### scShift enables cross-dataset comparisons and *in silico* biological state modeling

We next demonstrated scShift’s capabilities in cross-dataset analysis and *in silico* cell state modeling, focusing on donor-level CD4 T cell biological states. To compare donor and drug effects, we first constructed a kernel principal component space using drug perturbation conditions. We then projected the donor-wise median biological embeddings of CD4 T cells onto this space (Fig. 2f, Supplementary Methods). The first two kernel principal components of drug perturbation states corresponded to well-established T cell activity axes: inflammation states (measured by Interferon Stimulating Gene (ISG) expression) and T cell activation states (measured by expression of markers including *CXCR4, CD69*, Extended Data Fig. 2a). Comparison of the NF- kB inhibitor IMD-0354 with two other NF-kB inhibitors (IKK inhibitor VII and TPCA-1) revealed a distinct shift in T cell activation markers for IMD-0354 (Extended Data Fig. 2b), consistent with their positions on the kernel principal components.

We next analyzed how donor and drug responses compared to interferon and pathogen stimulation states from the two hold-out studies. First, we calculated the median biological embeddings for each interferon and pathogen condition and centralized them relative to control conditions. We then constructed a principal component space from these hold-out conditions and projected our donor and drug representations onto it (Supplementary Methods). This analysis revealed three distinct regions: 1. a “control region”; 2. an “ISG expression” wing, containing interferon and early pathogen stimulation conditions; and 3. a “neutrophil recruitment” region, characterized by 24-hour pathogen stimulation (Fig. 2g). Further analysis of donors and drugs within the high ISG wing revealed their associations with disease states and elevated ISG expression levels (Fig. 2h-i, Extended Data Fig. 2c, Supplementary Fig. 6). In contrast, fine-tuned scPoli and scVI models failed to merge interferon and short-term pathogen stimulation conditions, and did not properly align hold-out conditions with donors or drugs showing high ISG expression (Supplementary Fig. 7). This limitation likely stems from the improper handling of batch effects and lack of identifiability (Supplementary Note).

### scShift characterizes universal lung fibrosis states

Lung fibrosis has been a major health concern for decades, marked by scarring and thickening of lung tissue that impairs respiratory function. While various factors can trigger this condition - including environmental toxins, certain medications, and autoimmune disorders - many cases remain idiopathic (IPF). The recent discovery of lung fibrosis in COVID-19 patients, termed post-COVID-fibrosis [42, 43], adds urgency to understanding its mechanisms and developing effective treatments. scShift offers a unique opportunity to align diverse patients, datasets, and experimental systems, enabling a universal characterization of lung fibrosis.

We trained the scShift model on the CellXGene lung compendium, comprising 3,433,014 cells from 697 donors across 54 datasets (Fig. 3a). The scShift unperturbed embedding achieved batch integration performance comparable to alternative methods as quantified by scIB scores (Extended Data Fig. 3), confirming our findings from the blood model. Given myeloid cells’ critical role in lung fibrosis [44], we focused our initial investigation on analyzing fibrotic states within these cells. We assembled a comprehensive collection of lung fibrosis datasets, including pulmonary fibrosis samples from the Human Cell Lung Atlas (HLCA) [45], blood samples from IPF patients [46], and patients with fibrotic Hypersensitivity Pneumonitis (fibrotic HP) [47]. We also included two datasets showing post-COVID-fibrosis phenotypes: one from lethal COVID-19 cases [48] and another examining bronchoalveolar lavage fluid immune cells from severe COVID-19 patients [49].

We trained donor-level classifiers for fibrosis phenotypes using HLCA pulmonary fibrosis datasets as the training set, comparing embeddings generated by different methods (see Methods). We evaluated these classifiers using both hold-out patients from the training set and other datasets from our collection. While all methods achieved strong performance (AUC near 1) in the training set, with scShift performing marginally lower, scShift demonstrated superior performance in predicting fibrosis and COVID-19 phenotypes across hold-out datasets compared to alternative models fine-tuned on these datasets (Fig. 3b). In blood datasets (Blood IPF, Fibrotic/Non-Fibrotic HP), scShift classifiers showed reduced but still above-average performance (AUC ∼0.7) compared to lung datasets, indicating attenuated yet persistent pathologic signatures in blood myeloid cells. To validate our classifiers’ accuracy in capturing fibrosis phenotypes in COVID-19 patients, we conducted additional analyses on both COVID-19 datasets. In the first dataset, we found a significant positive correlation between pathologic fibroblast fraction (annotated in the original study [48]) and classifier scores within COVID-19 patients (Fig. 3c). The second dataset, while lacking external annotations, included moderate COVID-19 patients previously reported to be free of fibrosis-associated signatures [49]. Our scShift classifiers showed worse-than-random performance on these moderate cases (Extended Data Fig. 4a), supporting their specificity for fibrosis phenotype classification. Comparison of genes prioritized by scShift-based correlation scores (see Methods) with those identified using a current gold-standard pseudobulk pipeline (DESeq2 [50– 52]) revealed systematic differences in prioritized genes (Fig. 3d). This is anticipated as differential expression testing methods unbiasedly prioritize COVID-19 and fibrosis signatures among patients.

To validate our findings from human COVID-19 cohorts, we analyzed a humanized mouse model dataset profiling lung immune cells following COVID-19 infection [53]. The dataset provided ground truth biological state annotations (as days post infection), enabling evaluations across different approaches similar to our benchmarks of interferon and pathogen stimulations, particularly for myeloid cells where cell subtype annotations were available. scShift demonstrated top performance in both disentanglement and label preservation metrics (Fig. 3e, Extended Data Fig. 5a-b, Supplementary Fig. 8). We then evaluated how COVID-19 and fibrosis classifiers performed in categorizing different post-infection timepoints in the dataset (see Methods). Classifiers accounting for cell type composition information (scShift unperturbed, scANVI) excelled at predicting COVID-19 infection phenotype at 7 and 14 days post-infection (dpi) with AUCs approximately 1 (Fig. 3f). The single-cell level scores from the optimal scShift classifier showed clear prioritization of CD14 monocytes and macrophages (Extended Data Fig. 5c). The classifiers based on zero-shot scShift biological embedding showed superior performance in identifying the fibrosis phenotype at 28 dpi compared to other approaches (Fig. 3f). Comparison of scShift classifier scores revealed that while the COVID-19 phenotype in myeloid cells disappeared by 28 dpi, fibrosis scores increased globally in myeloid cells at 28 dpi compared to 0 dpi, indicating a post-COVID-fibrosis phenotype consistent with the original study [53] (Fig. 3g). Furthermore, while COVID scores remained stable between 7 dpi and 14 dpi, fibrosis scores showed a continuous decrease from 0 through 14 dpi (Fig. 3g). Based on these observations, we identified gene lists aligning with COVID and fibrosis score patterns from 0 to 14 dpi respectively (Supplementary Methods).

We combined the humanized mouse model and lethal COVID-19 datasets to identify genes consistently upregulated in lethal-COVID-19/28 dpi conditions versus control (Supplementary Methods). These consistently upregulated genes were expected to contribute to universal post-COVID-fibrosis phenotypes across datasets.

In identifying a stringent list of 47 genes, the scShift-based approach outperformed pseudobulk differential expression significance (Supplementary Methods), achieving a high AUC (0.859) with balanced sensitivity and specificity (both approximately 0.8, Fig. 3h). This advantage persisted but diminished for a broader list of 229 genes (Extended Fig. 4b). Pathway analysis revealed distinct functionalities among genes associated with the COVID pattern, fibrosis pattern, and the 47-gene list (Fig. 3i). Specifically, COVID pattern genes showed COVID signatures and enrichment in oxidative phosphorylation, while fibrosis pattern genes displayed enrichment in cell adhesion, antigen presentation, and phagosome pathways. The 47-gene list was enriched in pathways associated with cell migration and resistance to multiple cell death mechanisms (Fig. 3j, Supplementary Table 3). We next used scShift to refine the 47/229 gene lists by filtering out false negative predictions based on scShift correlation scores (Supplementary Methods). This refinement resulted in a stronger enrichment of apoptosis and migration KEGG pathways in both gene lists (Extended Data Fig. 5d). Notably, this procedure filters out *MAF* but retains *MAFB*, two genes displayed similar log-fold changes under the lethal COVID-19 condition (Extended Data Fig. 5e). This aligns with recent studies proposing the *MAFB/MAF* ratio as a marker of severe COVID-19 and associated fibrosis phenotypes [54]. Together, our findings suggest that the scShift classifier, while solely trained on lung fibrosis datasets, is able to generalize to post-COVID-fibrosis phenotype in donor cohorts and animal models, providing a universal characterization of myeloid-fibrosis signatures.

We hypothesized that drugs affecting the fibrosis phenotype in myeloid cells could serve as potential therapeutic agents. We applied our scShift-based classifier to derive fibrosis scores in myeloid cells treated with various drug perturbations (previously used in blood models [35]). While none of these drugs was originally developed for treating fibrosis (most being experimental anti-cancer agents), several showed promising potential in reversing observed myeloid cell states (Fig. 3k). The predicted roles of these drugs are consistent with their effects on chemokine and interferon-stimulated gene (ISG) expression (Extended Data Fig. 5f). When comparing gene expression changes induced by top drug candidates with earlier derived gene lists, we observed relatively low AUC values, indicating limited gene expression overlap (Extended Data Fig. 6a). This modest correlation likely reflects multiple biological differences between myeloid cells in the humanized mouse model and the drug perturbation dataset, emphasizing the importance of extracting biological state shifts for cross-dataset analysis.

We further extended our probing approach beyond myeloid cells to analyze 10 major lung cell types (macrophages, monocytes, CD4 T cells, CD8 T cells, natural killer (NK) cells, B cells, Alveolar type I (AT1) cells, Alveolar type II (AT2) cells, fibroblasts, and endothelial cells). In the lethal COVID-19 dataset, different cell types required distinct scShift representations to maximize prediction accuracy, achieving AUCs ranging from 0.75 to 0.92 (Extended Data Fig. 6b). Using these optimal classifiers, we calculated 10 cell-type-specific fibrosis scores for all donors in the CellXGene compendium, encompassing 431 donors from 35 datasets (see Methods). Through estimating the network structure of these variables, scShift provides a unique opportunity to systematically analyze how different cell types contribute to human *in vivo* fibrosis. Causal discovery methods capable of handling latent confounders (such as FCI [55]) eliminate the conceptual need for dataset-specific variable standardization in network inference. Cross-validated Lasso and FCI analysis with stability selection yielded identical dependency network skeletons (edges regardless of directionality) across variables (Extended Data Fig. 6c) [55]. The final causal graph derived from FCI across multiple significance levels confirmed previously established links in fibrosis, such as the roles of macrophages and AT1 cells (Extended Data Fig. 6d). As expected, the graph also attributed links between similar cell types to potential latent confounders (CD4 T, CD8 T, and NK cells; monocytes and macrophages) (Extended Data Fig. 6d, Supplementary Fig. 10) [56, 57]. The derived link between T cells and monocytes may be supported by the observed increase of T-myeloid doublets at later timepoints in humanized mouse models and IPF patients [58] (Supplementary Fig. 9d-e).

### Emergent and scaling behaviors of scShift

scShift shows remarkable zero-shot capabilities when pretrained on huge compendiums of single-cell atlases, which according to our theory, only emerges when the training data covers a majority of possible biological states. Previous studies of large language models have defined two regimes of model behavior regarding model and training resources: 1. scaling laws that describe quantitative relationships between test loss and model size/training compute [59, 60]; and 2. emergent abilities that manifest only when training resources exceed certain thresholds. Characterizing these behavioral regimes in scShift is of particular interest, as it may provide a conceptual foundation for large-scale biological models.

We examined four key parameters in the scShift model: the hidden dimensionality size and three parameters of the training set: cell number, donor number, and dataset number. Different settings of latent variable dimensionalities were not included in the study due to their possible interactions with disentanglement attribution and evaluation metrics. In the blood model, drug perturbations were counted as both donors and datasets. We systematically evaluated model performance by individually downsampling each parameter while maintaining fixed training epochs for the scShift blood model. Analysis across different cell number settings showed that fixing training compute (more training epochs for smaller training sets) yielded results similar to the fixed epoch case when training compute exceeded 10% of the full model (Extended Data Fig. 7a). Our analysis revealed clear scaling behaviors for all four parameters in detecting biological states in interferon and pathogen stimulation datasets (see Methods, Fig. 4a). However, model performance showed substantial fluctuation when the donor/perturbation number fell below 10 percent (258 donors). Additionally, the model exhibited instability in settings where cell number was subsampled below 10% (Methods). The performance of unperturbed embedding in detecting cell types did not demonstrate a clear scaling behavior (Extended Data Fig. 7b).

Given the collinearity among cell, donor, perturbation, and dataset numbers, the quantitative relationship between these parameters and model performance remained unclear. To address this, we analyzed models from previous configurations and trained an additional set of models with varying donor/perturbation ratios (Methods). We focused our analysis on scenarios where donor numbers exceeded 10% of the total, a range beyond the fluctuation region and where training compute was not a limiting factor. By fitting model performance in detecting biological states against all variables, we identified three key factors: cell number, donor number, and hidden dimension size that explained model performance when donor numbers exceeded 258 (Fig. 4b, Supplementary Figs. 11-12, Tables. 1-2). The observed scaling relationship (*R*^2^ = 0.50, Pearson r = 0.71) is considered notable for zero-shot performance, where model behavior is typically less predictable than test loss [59, 60]. The scaling relationship with respect to a fixed architecture can be approximately simplified as follows:

Model performance ∝ log(Donor number × Cell number) + const.

This finding supports our theoretical framework that scaling dataset size directly enhances the model’s ability to identify biological states. It additionally suggests that incorporating additional datasets and donors would improve model performance more effectively than increasing cell numbers alone, as the former jointly increases both donor and cell numbers. The impact of hidden dimensionality was not included in the equation due to its undetermined interactions with data size under current experimental settings.

We next investigated model performance in the low-donor-number regime (fewer than 10% (258) of donors from the training set). To determine if the observed performance fluctuations were due to insufficient training, we tested scShift models with donor percentages of 2%, 5%, 10%, and 25%, with additional 20 experiments for each configuration, increasing training epochs by 5x and 2x for the 2% and 5% settings respectively. Unexpectedly, the 5% donor setting showed both lower median performance and higher variance (Fig. 4c). Training convergence was confirmed in the 5% setting by consistent results across different training epochs, which rules out the potential explanation of insufficient training (Extended Data Fig. 7c). After removing outliers in 10% and 25% settings only, the variance at 5% was significantly different from both 10% and 25% settings (Levene’s test, P=0.01, 0.004), while the 2% setting showed no such difference (P=0.33, 0.13). Mean performance increased significantly when moving from 5% to 10% and from 10% to 25% (One-sided Mann-Whitney U test, P=0.007, 0.0004). These findings suggest a “phase transition” phenomenon exhibiting non-monotonic variance (Fig. 4c). This likely relates to the interpolation threshold in double/multiple descent phenomena [61–63]. Low-performing outliers in higher donor percentage settings may suggest additional donor-specific effects that warrant further investigation.

We next examined whether the scShift lung model performance exhibits a similar pattern. The zero-shot performance of the lung model in detecting biological conditions shows first a fluctuation phase then a sharp increase with respect to cell number, donor number, and dataset number (Fig. 4d, Extended Data Fig. 7d), resembling emergent behaviors described in large language models [64]. Within these settings, those altering cell numbers show both a higher baseline and higher-frequency fluctuations compared to the other two types, suggesting a distinct mechanism (Extended Data Fig. 7d). As the transition threshold in the lung model approaches the full dataset size, the scaling behavior is no longer observed. The ability of unperturbed embeddings to detect myeloid subtypes showed less clear relationship with dataset size but scaled consistently with hidden dimensionality (Extended Data Fig. 7e).

In linear probing tasks, non-monotonic patterns emerge in settings altering hidden dimensionality and donor numbers, while performance increases more uniformly (despite several outliers) in settings altering dataset numbers. This distinct pattern in dataset number scaling may reflect the additional factor of fibrosis dataset/- donor number in the training set (Extended Data Fig. 7f). Evaluations of lethal COVID-19 classification AUCs and correlation levels with pathologic fibroblast ratios indicate different transition thresholds with respect to donor numbers (Extended Data Fig. 7g). The higher transition threshold for classification AUCs compared to correlation levels may reflect its additional requirement of deconvolving COVID-19 signatures. All donor number transition thresholds in lung models (487, 417, 243) are comparable to those in blood (between 124 and 260, Fig. 4c). In summary, our evaluations of scShift models suggest two distinct phases regarding training data. The first phase, when the dataset is insufficient, exhibits fluctuation and no clear increase in mean performance. Beyond a threshold related to donor number, task complexity, and other dataset-specific properties, the model exhibits consistent performance gains in terms of donor and cell numbers (Fig. 4e).

## Discussion

We presented scShift, an identifiable variational inference framework that enables zero-shot query of cellular states. To our knowledge, scShift is the first model capable of revealing biological states without prior annotations. By leveraging atlas-level scRNA-seq compendiums during pretraining, scShift acquires remarkable zero-shot capabilities, facilitating seamless adoption and interpretation of new datasets. Through our lung fibrosis case study, we illustrated scShift’s ability to characterize and generalize complex disease states in unseen contexts, enabling various novel analyses. As far as we know, scShift shows the first clear scaling law and emergent behavior observed in models of single-cell RNA sequencing data.

Despite its strong zero-shot capabilities, scShift may not generate meaningful outputs when analyzing datasets from entirely different contexts, such as those containing unseen cell types. Integration of epigenetic data or prioritization of gene regulatory information may enhance the model’s transfer learning abilities in such contexts [20, 65, 66]. Current scShift models are trained on blood and lung scRNA-seq atlases, two tissues with diverse single-cell profiles. Extending the framework across different tissues, eventually to all cell types across the entire human body, may require additional model designs due to violation of the generative process assumptions. While scShift currently uses a variational autoencoder architecture with multiple layer perceptron (MLP) backbone, future studies may evaluate the use of more advanced neural network architectures and particularly examine their scaling behavior.

## Data availability

All datasets analyzed in this paper from previous publications are publicly available. The CellxGene blood and lung atlas dataset were downloaded from CellxGene census (https://chanzuckerberg.github.io/cellxgene-census/) via gget [6] with version 2023-07-15. The drug perturbation dataset was downloaded from Kaggle (https://www.kaggle.com/competitions/open-problems-single-cell-perturbations). The interferon stimulation dataset was downloaded from dryad (https://datadryad.org/stash/dataset/doi:10.5061/dryad.4xgxd25g1). The pathogen stimulation dataset (EGAS00001005376) was downloaded from https://eqtlgen.org/sc/datasets/1m-scbloodnl.html.

## Code availability

We have made scShift available as a public open-source Python package, which can be accessed at https://github.com/MingzeDong/scShift.

## Acknowledgements

Y.K. disclose support for the research of this work from NIH [R01GM131642, UM1PA051410, U54AG076043, U54AG079759, U01DA053628, P50CA121974, and R33DA047037].

## Author contributions

M.D. conceived the study, developed scShift, established its theoretical foundation, and performed the computational analysis in the study. K.A. provided assistance in analyzing humanized mouse model data. E.S., R.A.F. provided feedback and supervision on the fibrosis data analysis. R.F. and Y.K. provided overall supervision of the study. M.D., and Y.K. wrote the manuscript.

## Methods

scShift is a deep identifiable variational inference framework designed to uncover both shared information and dataset-specific biological states in atlas-level scRNA-seq datasets. A rigorous treatment of its assumptions and theoretical identifiability is presented in the Supplementary Note. In the Methods section, we focus on the practical implementation of scShift and discuss its connection with the theoretical guarantee.

## Generative model of scShift

scShift considers the following generative process for modeling the distribution of entries *x*_*ig*_ in the count matrix of the single-cell RNA-sequencing data. For simplicity, we drop the cell index *i* in the formula:

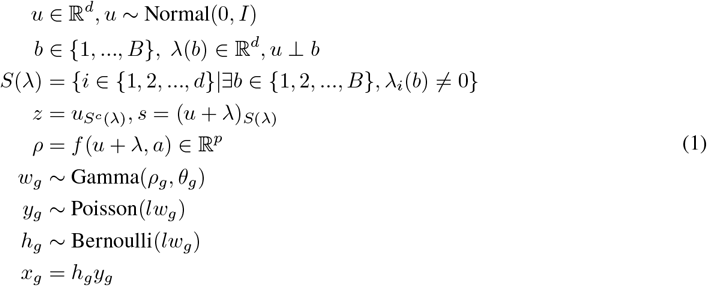

In the generative process, the latent variable *u* represents the centralized full state of each cell. *b* represents the dataset batch, and *λ*(*b*) is a projection of *b* on the latent space indicating the dataset-specific encoding. *b* should be considered as a fixed constant for each cell. *S*(*λ*) is a set containing indices with at least one non-zero entry among *λ*(1), …, *λ*(*B*). *S*^*c*^(*λ*) denotes the complement of *S*(*λ*), which contains the indices where *λ*(1), …, *λ*(*B*) are all zero. Based on *S*(*λ*) and *S*^*c*^(*λ*), *u* + *λ* can be decomposed into the batch-independent latent variable *z* and batch-dependent latent variable *s. a* is an optional label that accounts for other possible experimental covariates such as sequencing assays. *ρ* ∈ ℝ^*p*^ represents the mean of gene expression output by neural network decoder *f* . The function *f* enforces 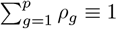 for each cell through a softmax layer. *θ* ∈ ℝ^*p*^ defines the gene-specific shape parameter of the Gamma distribution. *l* is fixed as the observed library size of each cell. *y*_*g*_ follows a negative binomial (NB) distribution by its Gamma-Poisson mixture representation. *h*_*g*_ represents the zero inflation level. Together, *x*_*g*_ is modeled as a sample from a zero-inflated negative binomial (ZINB) distribution. The generative process of scShift from *ρ* to *x* aligns with the scvi-tools framework [9, 25].

### Model theoretical identifiability

Without additional constraints, it is known that a non-linear generative process is generally not identifiable [23, 24]. In our context, the ground truth latent variables *z, s* can be arbitrarily entangled in *u*+*λ*. Our theoretical analysis demonstrates that the desired disentanglement can be achieved in solutions with minimal size of *S*(*λ*) under moderate assumptions (Supplementary Note). Our theoretical result is relevant to, but fundamentally different from, the recently proposed sparse mechanism shift framework [41, 67]. Our work considers a generative process where the label encoding *λ* affects only part of the latent variables. We consider solutions with the sparsest batch-dependent components and prove that both batch-dependent and batch-independent variations are identifiable up to disentanglement equivalence (Supplementary Note). In comparison, the sparse mechanism shift framework relies on the effect of supervision *λ* on all components. The framework further considers the sparsity of interaction graphs in time-series data [67], and achieves identifiability with respect to permutation equivalence. Our theoretical support also relates to but differs from [68], which considers a spatial-aware generative process and proves that with auxiliary spatial information, intrinsic and spatial variations can be identified with respect to a slightly milder form of disentanglement equivalence.

### Approximate posterior inference of scShift

In order to infer the parameters in the scShift model, we approximate the posterior distribution via autoencoding variational Bayes. The posterior distribution *p*(*u*|*x, a*) is approximated by the following variational posterior family:

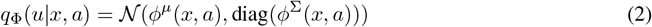

Here *ϕ*^*µ*^, *ϕ*^Σ^ : ℝ^*p*^ → ℝ^*d*^ denotes encoders for mean and variance terms in the variational posterior. Then the evidence lower bound (ELBO) can be derived via straightforward computation.

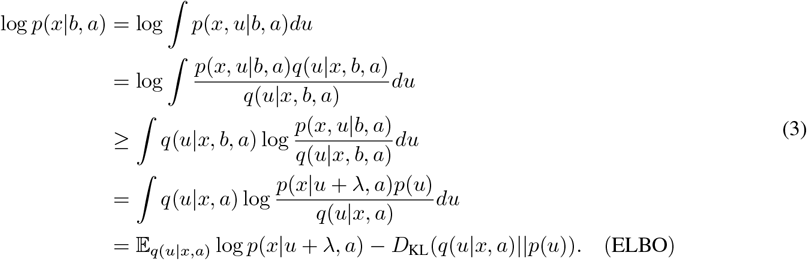

In addition to the ELBO, our theoretical analysis suggests two essential designs for model identifiability. One is that the model should learn the parameters with the sparsest / smallest *S*(*λ*); the other is that the latent variable *u* should be independent of the batch label *b*. We next describe the model designs that address both points.

### Sparsity regularization through probabilistic relaxation

Minimizing the batch-dependent biological component number |*S*(*λ*)| is not a differentiable objective. To effectively incorporate the sparsity regularization in the loss term, we consider the following probabilistic relaxation of |*S*(*λ*)|, where we model 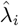 by a variational posterior conditioned on the dataset label *b*, and *λ* is an element-wise truncation of 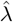, with the threshold *ϵ* = 0.1:

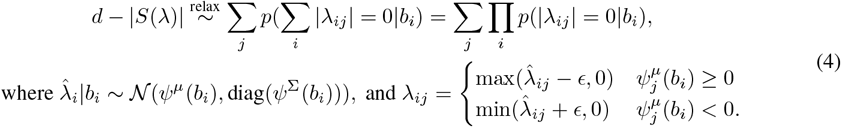

Nevertheless, maximization of the term involves taking product across all cells or batches, making the term difficult to optimize. Here we instead consider the following regularization:

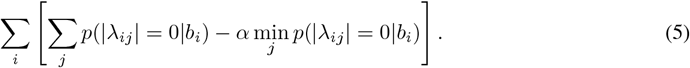

The newly designed regularization encourages overall increase of all probabilities *p*(|*λ*_*ij*_| = 0|*b*_*i*_). Additionally, by the second term where we set *α* = 2, it enforces each *λ*_*i*_ will have at least one non-zero entry with *p*(|*λ*_*ij*_| = 0|*b*_*i*_) = 0. This implies non-zero batch effects and aligns with Assumption 2 in our theoretical justification. Although this regularization term differs from the original by promoting sparse label encoding within batch-dependent components, we empirically achieve effective disentanglement. This success likely stems from the fact that *S*(*λ*) is constructed using the aggregate of all datasets, thus is not sensitive to individual dataset sparsity. Empirically, the penalty led to two essential properties for disentanglement that were not achieved by alternative models, including CPA and sVAE [40, 41]: sparsity in the dataset label encoding and linear independence among components of the latent space (Supplementary Fig. 2).

We next derive the sparsity regularization based on the probabilistic relaxation. Denoting the Gaussian error function as erf(), Then the probability of *λ*_*ij*_|*b*_*i*_ being zero can be expressed as follows.

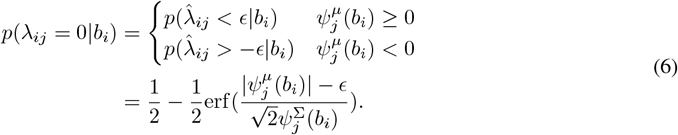

The final sparsity regularization to be maximized is of form

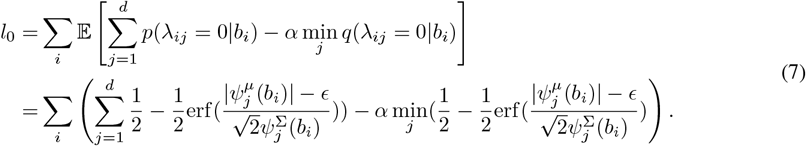

### Independence regularization

Apart from the sparsity regularization, an explicit regularization for the independence between *u* and *λ* is needed for the scShift model. We formulate the regularization term leveraging the following observation: When *u* and *λ* are independent, the distribution of *u* + *λ*|*b* is a shifted Gaussian distribution as follows. With a slight abuse of notation, we denote *λ*(*b*) as the expectation of the previously described probabilistic relaxation.

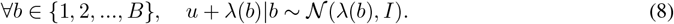

Taking advantage of the above formula, we aim to quantify the discrepancy between *q*(*u* + *λ*(*b*)|*b*) and 𝒩 (*λ*(*b*), *I*) as an indicator for the dependence level between *u* and *λ*. We employ kernel MMD (kernel Maximum Mean Discrepancy) to measure the discrepancy as implemented in the InfoVAE work [31]. Additionally, we calculate the kernel MMD between *q*(*u*) and 𝒩(0, *I*) to enforce the standard normality of *q*(*u*), in which case the previously described term is a meaningful measure of independence. We only update *u* (but not *λ*) based on the MMD regularization terms. Denoting the sum of the two MMD regularization terms as *l*_MMD_, the final optimization problem can be formulated as

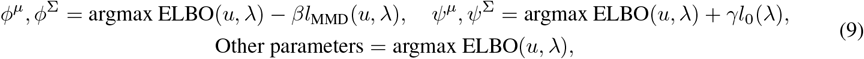

where *β* and *γ* are weights of the regularization terms. After the scShift model is trained, it returns both *u* and *λ* for cells in the training set. We then use the obtained *S*(*λ*) to separate the unperturbed components and the biological components as 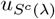 and *u*_*S*(*λ*)_ respectively. For new datasets not in the training set, we only return *u* and separate unperturbed and biological components using *S*(*λ*) defined from the training set.

### Donor-level phenotype classification

To harness scShift’s zero-shot capabilities for transfer learning, we developed a linear probing procedure to extract disease or biological-state-associated features from scShift representations. Since biological states may involve changes in cell type composition, gene expression shifts, or both, we construct probing classifiers at a donor level. This method builds upon linear probing benchmarking frameworks widely used in representation learning [33, 34, 69], and closely resembles recently proposed methods for multi-instance-learning based disease classification in single-cell data [70].

A simple aggregation of all cells for each donor would imply equal weights across cells, resulting in a loss of expressivity. Therefore, we considered a generalized linear classifier with learnable weights for each cell. Denoting the embedding for cell *i* as *x*_*i*_ ∈ℝ^*d*^, and donors {1, 2, …, *k*}, with ground truth one-hot labels *y* ∈ℝ^*m*^ for each donor, we denote the model parameters as *β*_1_ ∈ R^*d*^, *b*_1_ ∈ R, *β*_2_ ∈ R^*d*^*×m, b*_2_ ∈ R^*m*^, the total cell number in donor *j* as *Z*_*j*_, then the classifier model can be formulated as:

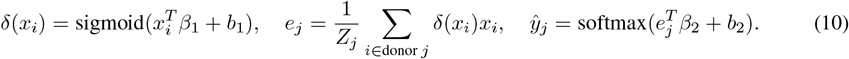

Finally, we minimize the cross entropy loss between *y*ŷ and *y* with mini-batch gradient descent. We implemented batch normalization in the classifier to accelerate convergence. However, we observed that sometimes not using batch normalization may lead to better overall performance across different methods, presumably due to the implicit early stopping mechanism. This mainly affects alternative methods (scVI, scANVI) as their behavior on hold-out datasets is not robust due to the lack of identifiability. In these cases, we report the results without batch normalization.

We also implemented a version of the classifier (“conditioned”) that accounts for both unperturbed and biological embeddings in the model, which fully leverages scShift’s disentanglement power. Denoting the unperturbed and biological embeddings for cell *i* as *z*_*i*_, *s*_*i*_, the model can be formulated as:

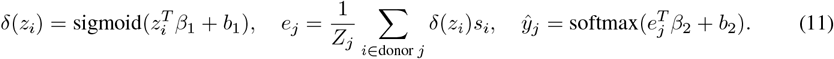

Finally, for donor-level embeddings (such as the output of scPoli) *x*_*j*_, we constructed an ordinary linear classifier:

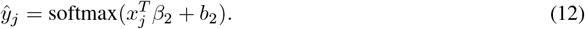

After the model is trained, it can output disease scores at either single-cell or donor level. Following Eq. (10), for donor *j*, the donor-level score is expressed as *y*ŷ_*j*_, while for cell *i*, the single-cell level score is defined as softmax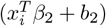.

## Model training

We trained scShift models for blood and lung tissues in this study. For the blood model, we combined the subsampled CellxGene blood compendium (containing 1 million cells from 30 studies and 2,538 donors) and a drug perturbation dataset comprising 240,090 PBMCs from three donors with 144 drug perturbations [35] and three other negative/positive control conditions. Each drug condition was treated as one dataset. Using the Scanpy pipeline [71] with default settings, we performed library size normalization, log1p transformation, and selected 5,036 highly variable genes from the drug perturbation dataset. We used raw expression counts of the highly variable genes for model training. We further annotated coarse-grained cell type labels for these cells, which were used in benchmarking (Supplementary Methods).

The lung model was trained on the CellxGenes lung compendium, which contained 3.4 million cells from 697 donors across 54 studies, excluding the lethal COVID-19 dataset [48]. Following the same preprocessing pipeline used for the blood model, we selected 5,000 highly variable genes from the lethal COVID-19 dataset [48]. These genes were then intersected with the Human Lung Cell Atlas (HLCA) [45] gene set, resulting in 4,634 genes. We used raw expression counts of these genes for model training. We further annotated coarse-grained cell type labels for these cells, which were used in constructing balanced mini-batches for training and later benchmarking (Supplementary Methods).

Both models shared a common architecture of 100 latent components and 1,000 hidden components. For balanced training, we employed a semisupervised data splitter: the blood model balanced the dataset label with a batch size of 400, while the lung model balanced donor and coarse-grained cell type labels with a batch size of 500. Training was conducted for 50 epochs for the blood model and 20 epochs for the lung model.

### Benchmarking on hold-out datasets for the blood model

We used two hold-out datasets with ground truth biological conditions for benchmarking of the blood model: the interferon stimulation dataset [36] and the pathogen stimulation dataset [37]. Full details regarding the implementation and parameter settings of all comparison methods are provided in Supplementary Methods. For the interferon stimulation dataset, we used the PCA embedding from the original work as the baseline pre-integrated embedding [36]. For the pathogen stimulation dataset, we employed the v3 dataset in the data repository containing 302,598 cells. The baseline pre-integrated embedding was set as the PCA embedding computed from the top 2,000 highly variable genes with library normalization and log1p transformation implemented with Scanpy default settings [71]. For the interferon stimulation dataset, we ran two evaluations on the unperturbed (cell type) representation across different methods. For both evaluations, we used the cell type annotation as the label key, while using batch/perturbation as the batch key respectively. The computed scIB metrics from the two evaluations were averaged to generate the final result table. For the pathogen stimulation dataset, we used the cell type annotation as the label key and the perturbation label as the batch key.

As different cell types may have different responses to certain stimulations, benchmarking removal of cell type signatures in the biological embedding may no longer be meaningful. We took advantage of the observation in the original analysis [36] that responses of CD4 T, CD8 T and natural killer (NK) cells to interferon stimulations group together. Analysis in [37] also showed that CD4 T and CD8 T cells have shared responses to pathogens with minimal distinct differentially expressed genes. Thus, we quantified the scIB metrics in these cell populations to cover both disentanglement and biological state metrics, using perturbation as the biological label and cell type as the batch key. We found that for the interferon dataset, most approaches generated negative PCR scores; therefore, this metric was excluded from the comparison. The PCR score was also noted to be excluded from a recent study [72]. Each individual metric was rescaled to have a minimum of 0 and maximum of 1. The rescaled overall score was defined as the 60/40 weighted mean of rescaled biological conservation metrics and batch correction metrics, consistent with scIB default [18].

### Unsupervised characterization of donor-level CD4 T cell biological states

As previously described, the biological embedding may differ from ground truth biological variation by a dataset-specific constant. Therefore, we implemented two heuristic procedures for treating biological components for the donor cohort in the training set. The first is to globally subtract the median biological components of all cells. The second is to perform a dataset-specific centralization with respect to the normal state in each dataset. In our study, the normal state is selected to be all cells from healthy donors or negative control drugs. The centralization constant of these datasets is thus calculated as the mean of median biological embeddings of the healthy donors (or the median biological embedding of the control drug). For datasets that do not have such control state, we trained a Ridge regression model with cross-validation (RidgeCV) on the known datasets for predicting the shift using the dataset label encoding *λ*. We then used the predictor to impute the shift vector for datasets without healthy donors. If the dataset encoding *λ* does not contain information of the healthy state, then the standardization reduces to no standardization on these datasets. Empirically, we observed that the two schemes lead to negligible differences in unsupervised analysis ranging from UMAP to principal component projections of the CD4 T cell subset (No adjustment: Supplementary Fig. 1c-d; Adjustment 1: Fig. 2g-h, Extended Data Fig. 2c; Adjustment 2: Fig. 2c,f, Extended Data Fig. 2a-b, Supplementary Fig. 1e-f). Additional details on Fig. 2f-g and comparative analysis are presented in Supplementary Methods.

### Fibrosis probing-based analysis

We utilized the lung model to develop classifiers for predicting pulmonary fibrosis phenotypes. Drawing from the Human Lung Cell Atlas [45], we analyzed datasets containing both normal and pulmonary fibrosis patients, selecting donors with these respective phenotypes. This selection yielded 5 datasets encompassing 143 individuals and 526,257 cells. We then focused on the myeloid population, specifically cells identified as macrophages or monocytes, resulting in a subset of 142 individuals and 161,069 cells. Given the moderate number of donors in the training set, we employed different evaluation strategies: for hold-out datasets, we used the complete set as the training data, while for training set evaluation, we implemented a 60/40 split ratio between training and test sets. We applied batch normalization in the classifiers to evaluate all datasets except those involving post-COVID fibrosis (including the humanized mouse data analyzed later).

We evaluated the scShift donor-level fibrosis score using the lethal COVID-19 dataset [48], comparing it with each donor’s ratio of pathological fibroblasts to total fibroblasts as labeled in the study. We conducted pseudobulk level differential gene expression analysis using DESeq2py [50–52]. We generated single-cell level scShift scores for the dataset using the optimal scShift classifier in terms of disease classification AUC. We further computed gene-level scShift correlation scores for the dataset through a two-step process. First, we calculated the Spearman correlation between log-normalized gene expression and single-cell level scShift scores for each lethal COVID-19 patient. We then determined the median correlation value for each gene across all patients. For visualization in Fig. 4d, genes with pseudobulk p-values *<* 0.05 were labeled as “significant DE genes”, and genes with absolute correlation scores *>* 0.12 were labeled as “scShift prioritized genes”.

We further tested different methods’ performance on this humanized mouse dataset using COVID-19 and fibrosis linear probing classifiers. For the COVID-19 classifier, we selected datasets from HLCA [45] that contained either more than 100 cells from healthy donors or more than 100 cells from COVID-19 patients. This yielded a total of 292,283 myeloid cells from 213 donors across 27 datasets. The fibrosis classifiers were the same as those used for fibrosis classification in hold-out datasets. Using a fixed random seed, we divided the humanized mouse data from each timepoint (0, 4, 14, and 28 days post-infection) into five segments. For classification purposes, we labeled the 4 and 14 dpi segments as COVID-19 and the 28 dpi segments as pulmonary fibrosis. We additionally included scPoli donor-level embeddings fine-tuned on the humanized mouse dataset, using the 20 segments as donor identity. The fine-tuning setting was the same as that for scPoli blood model fine-tuning. We generated single-cell level scShift COVID/fibrosis scores for the dataset using the optimal scShift classifier in terms of COVID/fibrosis classification AUC.

We constructed fibrosis classifiers for 10 major lung cell types: macrophages, monocytes, CD4 T cells, CD8 T cells, natural killer (NK) cells, B cells, Alveolar type I (AT1) cells, Alveolar type II (AT2) cells, fibroblasts, and endothelial cells. The classifiers from optimal settings (among ‘unperturbed’, ‘biological’, and ‘all’) for each cell type were employed to compute donor-level cell-type-specific fibrosis scores across all training set donors. For donors missing three or fewer cell types, we imputed missing values using the median fibrosis score of the corresponding cell type. We excluded donors missing more than three cell types, resulting in a final set of 431 donors from 35 datasets. We then applied Lasso with 5-fold cross-validation, and causal discovery methods PC and FCI [55] using the causal-learn implementation [73]. To ensure robust results, we employed stability selection [74], by executing each causal discovery algorithm 100 times, each using a random 70% subset of the data. Edges appearing in over 50% of runs were retained, with their types determined by majority voting.

### Scaling analysis

For the scaling analysis, we systematically adjusted the hidden dimensionality and tested various subsampling ratios based on the number of cells, donors, or datasets in the training set. In addition to the full model, we evaluated the following configurations of the scShift blood models, as presented in Fig. 4a (taking the integer portion for settings that involve percentage):

- Hidden dimensionality: [100,250,500,630,800];
- Cell number: [37202, 62004, 100000, 150000, 250000, 350000, 500000, 700000, 800000, 900000]. Settings with 12400, 24801, and 86806 cells were excluded due to training instability.
- Donor/perturbation number: [1.5%, 2%, 2.5%, 3%, 4%, 5%, 7%, 10%, 15%, 25%, 35%, 50%, 70%] of the total donor/perturbation number (2685);
- Dataset/perturbation number: [1.5%, 2%, 2.5%, 3%, 4%, 5%, 7%, 10%, 15%, 25%, 35%, 50%, 70%] of the total dataset/perturbation number (177).

For each trained model, we computed two performance metrics: cell type scores and biological state scores. Both scores were calculated as the average of scIB metrics (excluding the isolated F1 score for computational efficiency) evaluating unperturbed / biological embeddings. We evaluated these metrics on the complete interferon stimulation dataset and a subsampled pathogen stimulation dataset containing 50,000 cells from control and three-hour stimulation conditions. All scIB metric calculations followed the same benchmarking settings described earlier.

In fitting the scaling function, to address collinearity in the regression, we additionally trained models with decoupled donor and perturbation numbers. Specifically, the following settings were used: [(10%, 5%), (10%, 20%), (10%, 40%), (10%, 80%), (25%, 5%), (25%, 20%), (25%, 40%), (25%, 80%), (50%, 5%), (50%, 20%), (50%, 40%), (50%, 80%)] as percentages of the total donor and perturbation numbers (2,538 and 147, respectively). To evaluate the model’s behavior in the low-donor-number regime, we subsampled donor/perturbation numbers to 2%, 5%, 10%, and 25% each 20 times and trained the scShift model based on the resulting datasets.

For the scaling analysis of the lung model, following a similar approach as with the blood model, we individually adjusted the hidden dimensionality, cell, donor, and dataset numbers. The specific settings were (taking the integer portion):

- Hidden dimensionality: [150, 250, 350, 500, 700];
- Cell number: [1%, 2%, 3%, 5%, 7%, 10%, 15%, 25%, 35%, 50%, 70%] of the total cell number (3,433,014);
- Donor number: [1%, 2%, 3%, 5%, 7%, 10%, 15%, 25%, 35%, 50%, 70%] of the total donor number (697);
- Dataset number: [2%, 5%, 7%, 10%, 15%, 25%, 35%, 50%, 70%] of the total dataset number (54).

We generated both unperturbed and biological embeddings for the humanized mouse and lethal COVID-19 datasets using each model. On the humanized mouse dataset, we calculated cell type and biological state scores, each defined as the average of scIB metrics, using the same settings as in our fibrosis analysis. For the lethal COVID-19 dataset, we evaluated model performance by training five linear probing classifiers with different random seeds, following the same setup as our fibrosis analysis. We assessed performance using three metrics: mean AUC for lethal COVID-19 prediction (truncated to 0.5 for values below 0.5), and mean Pearson/Spearman correlations between donor-level scores and pathological fibroblast ratios within lethal COVID-19 patients (truncated to 0 for negative values).

Please refer to Supplementary Methods for additional details including coarse-grained cell type annotations, benchmarking on training sets, implementations of alternative methods for benchmarking on hold-out datasets, ablation study, unsupervised analysis of donor-level CD4 T cell biological states, benchmarking on the humanized model dataset, definitions of gene sets in fibrosis analysis, and probing of fibrosis state on drug perturbations.

## Extended data figures

**Extended Data Fig 1.**
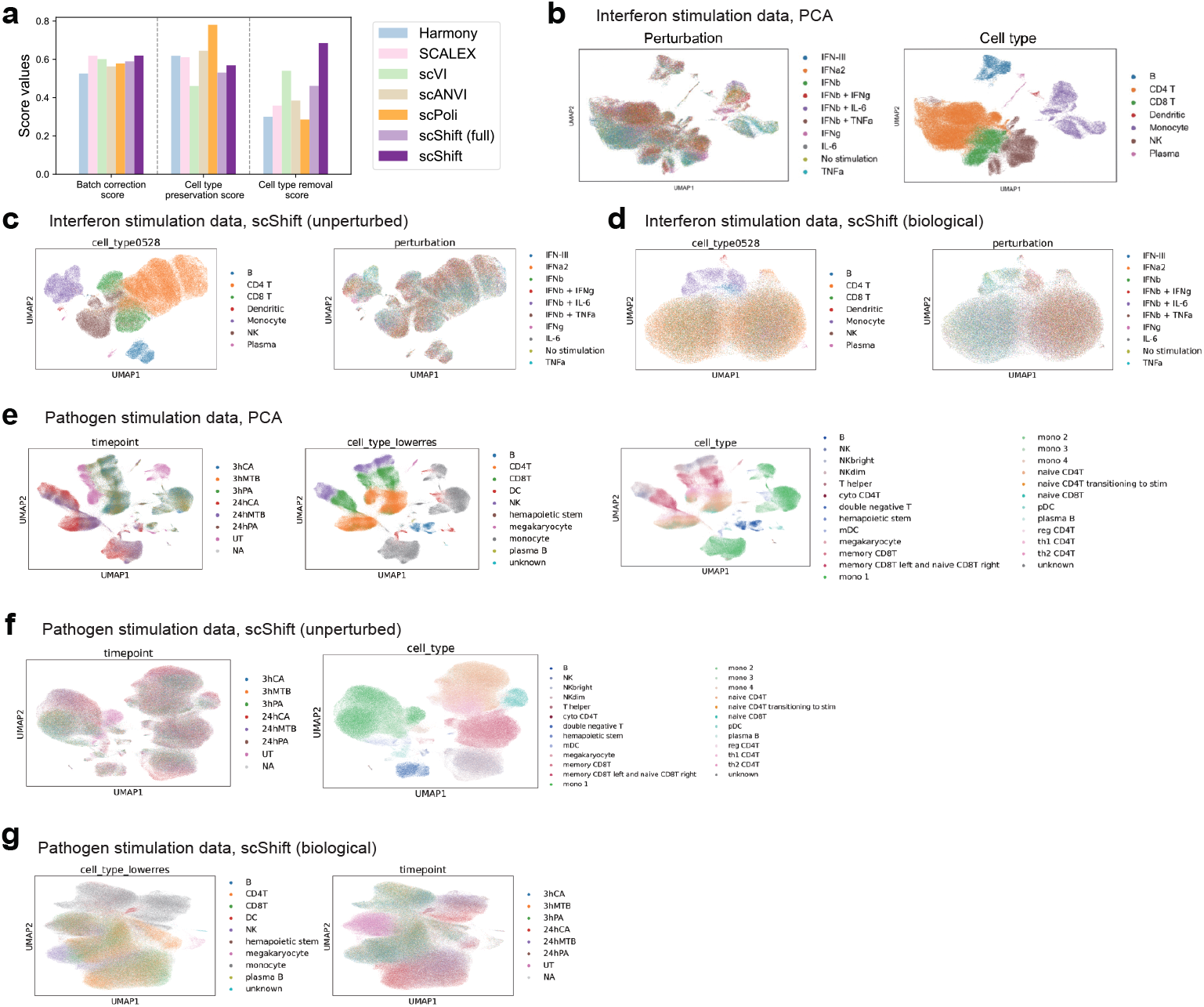
Additional benchmarking and visualization of the scShift blood model. **a**. Comparison of integration methods on a subsampled blood compendium training set (n=100,000) using scIB metrics for batch correction, cell type preservation, and cell type removal. ‘scShift (full)’ represents concatenated unperturbed and biological embeddings, while ‘scShift’ uses unperturbed embedding for the first two metrics and biological embedding for cell type removal. **b-d**. UMAP visualizations of interferon stimulation data using **b**. principal components from the original study, **c**. scShift unperturbed embedding, and **d**. scShift biological embedding, all colored by cell type and stimulation condition. **e-g**. UMAP visualizations of pathogen stimulation data using **e**. principal components, **f**. scShift unperturbed embedding, and **g**. scShift biological embedding colored by broad/fine-grained cell type and stimulation condition.

**Extended Data Fig 2.**
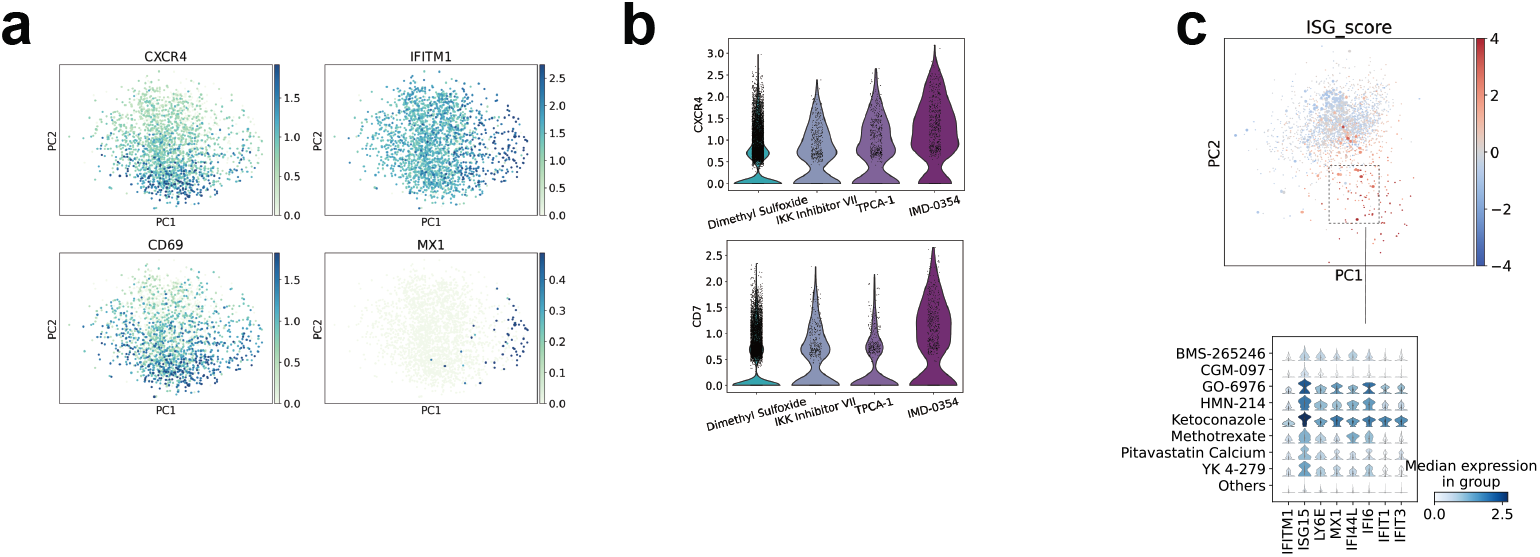
Additional results from integrative analyses across training cohorts, drug perturbations, and hold-out interferon/ pathogen stimulations. **a**. PCA visualization of CD4 T cell pseudobulk-level scShift biological embeddings from donors in the training compendium, colored by median log-normalized expression of representative genes. These coordinates are the same as those shown in Fig. 2f. **b**. Violin plots showing the log-normalized expression levels of *CXCR4* and *CD7* in cells treated with NF-kB inhibitors. **c**. Visualization of CD4 T cell median biological embedding per donor (smaller points) / perturbation (larger points) in the training compendium, colored by z-scores of combined interferon-stimulated gene (ISG) expressions (Upper); Stacked violin plots of log-normalized ISG expressions among representative drugs in the high ISG wing (Lower) versus other drugs. These coordinates are the same as those shown in Fig. 2g.

**Extended Data Fig 3.**
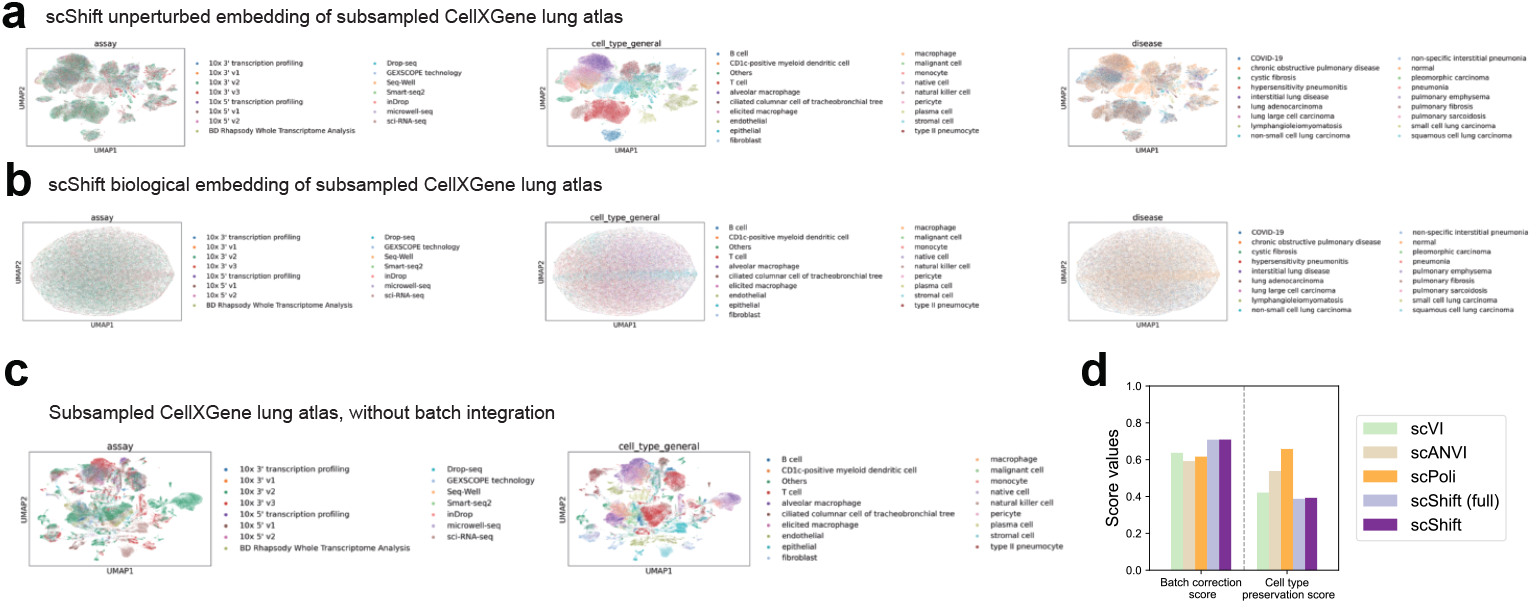
Visualizations and benchmarking results of scShift lung model. **a**. UMAP visualizations of the subsampled training compendium (n=100,000) using scShift unperturbed embedding, colored by sequencing assay, coarse-grained cell type, and disease condition. **b**. UMAP visualizations of the subsampled training compendium using scShift biological embedding, colored by sequencing assay, coarse-grained cell type, and disease condition. **c**. UMAP visualizations of the subsampled compendium before integration, colored by sequencing assay and coarse-grained cell type. **d**. scIB batch correction and cell type preservation scores of different atlas integration methods on the subsampled training compendium.

**Extended Data Fig 4.**
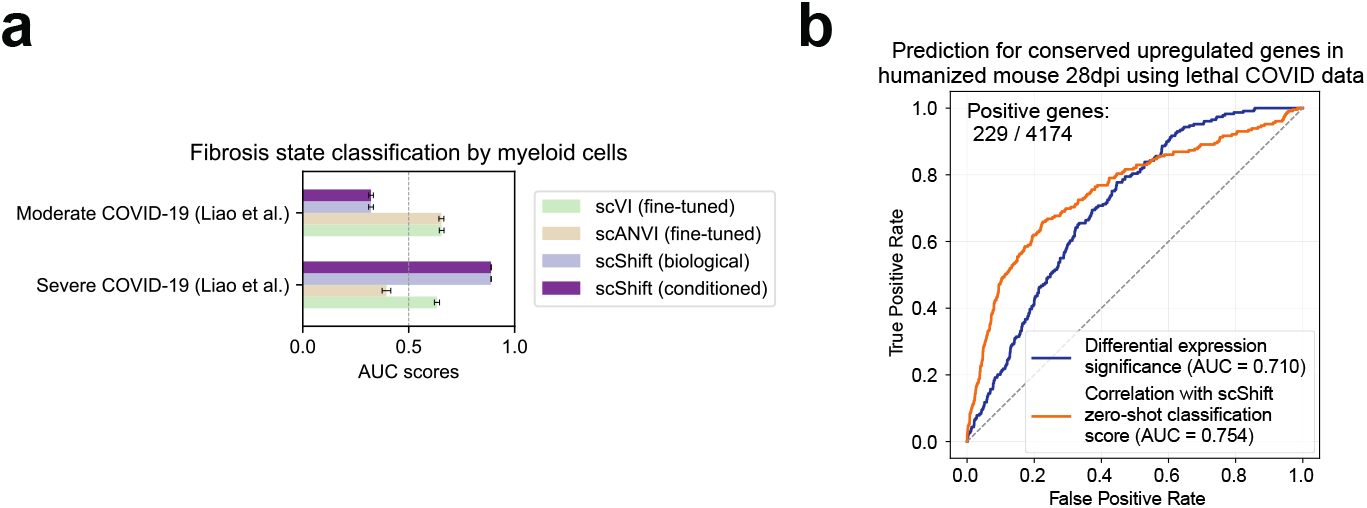
Additional results of post-COVID-fibrosis analysis. **a**. Barplot showing performances of different methods in predicting moderate/severe COVID-19 versus normal using myeloid cells from Liao et al. [49]. Bars represent mean prediction AUC values from 10 independent experiments with different random seeds, with error bars indicating standard error. **b**. ROC curves and AUC values for pseudobulk differential expression significance and scShift correlation scores in distinguishing consistently upregulated genes across both datasets using a loose threshold (n=229).

**Extended Data Fig 5.**
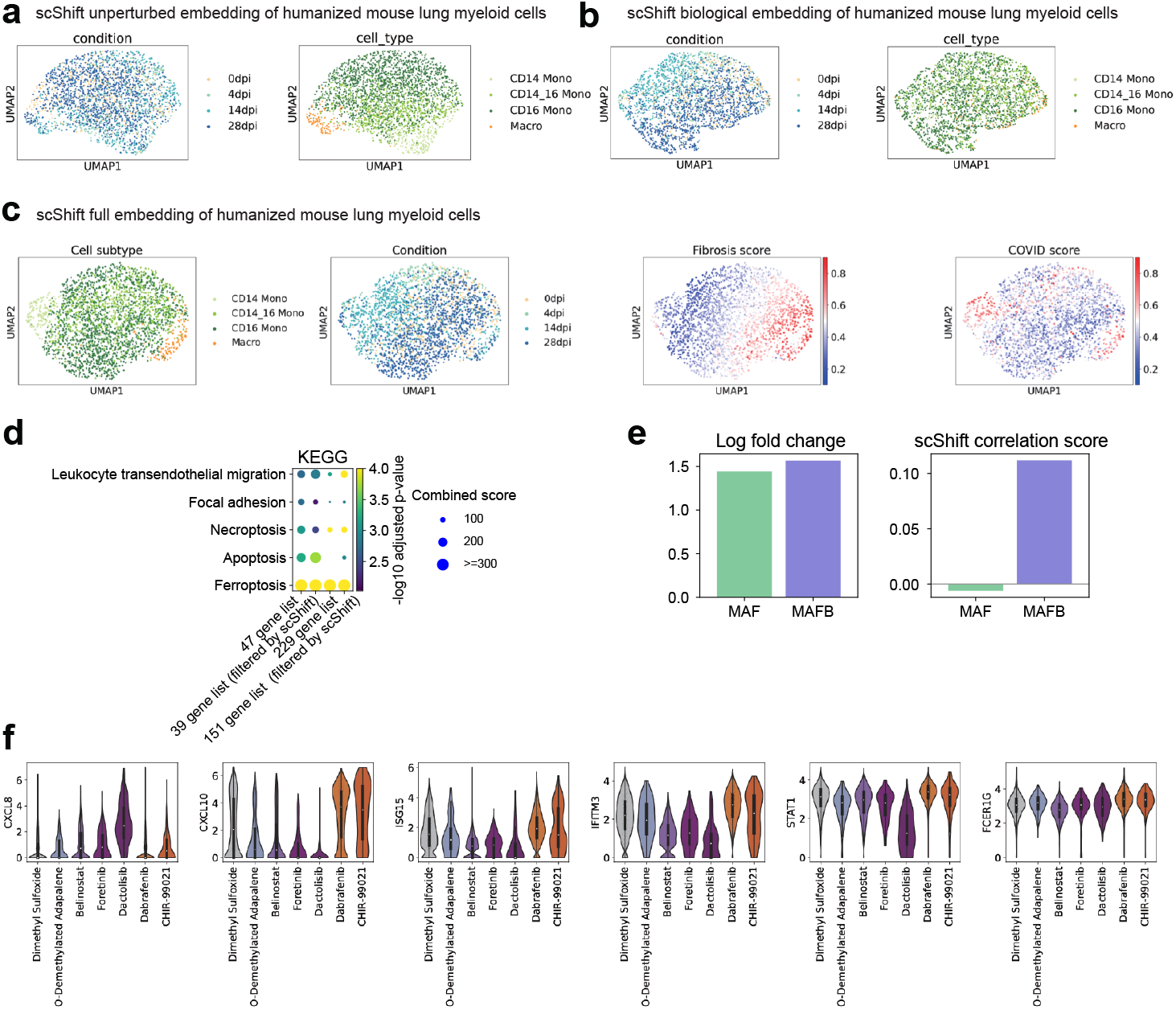
Additional results of fibrosis classification analysis in humanized mouse data. **a**. UMAP visualization of lung myeloid cells using scShift unperturbed embedding, colored by experimental condition and cell type. **b**. UMAP visualization of lung myeloid cells using scShift biological embedding, colored by experimental condition and cell type. **c**. UMAP visualization of lung myeloid cells using scShift full embedding, colored by experimental condition, cell type, fibrosis score, and COVID score. **d**. Dotplot showing KEGG pathway enrichment analysis of the 47-gene list, the 39-gene list (filtered by scShift), the 229-gene list, and the 151-gene list (filtered by scShift). **e**. Barplots showing the pseudobulk log fold changes of *MAF* and *MAFB* in lethal COVID-19 patients versus control, and the scShift correlation scores of these two genes. **f**. Violin plots showing log-normalized expression of example genes in myeloid cells across drug perturbations [35].

**Extended Data Fig 6.**
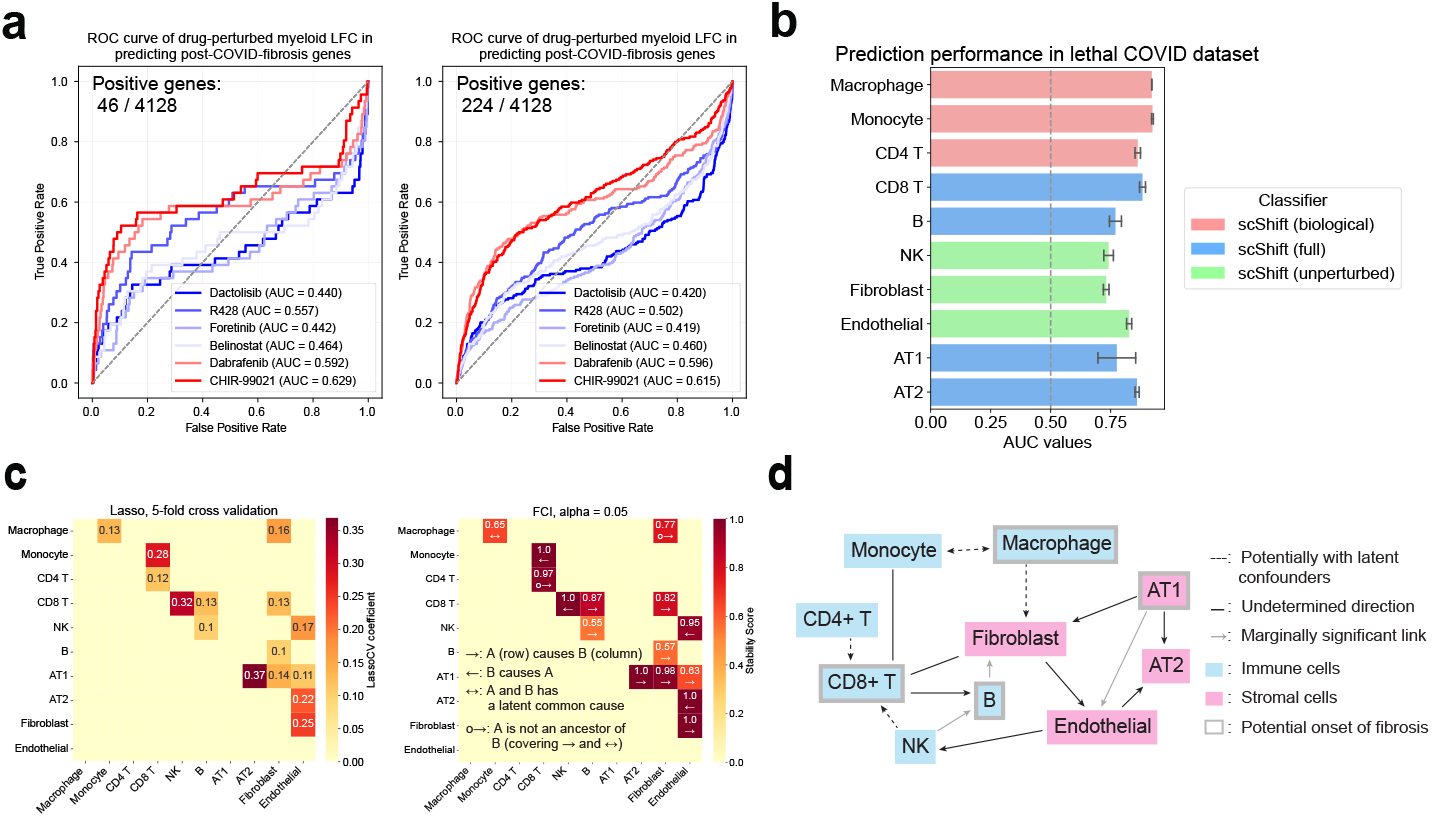
Analyses of scShift-identified therapeutic drug targets and causal modeling of fibrosis across cell types. **a**. ROC curves and AUC values showing drug perturbation log fold changes in predicting post-COVID-19 fibrosis genes in myeloid cells. Drugs were selected by their strongest effects in terms of fibrosis scores. **b**. Predictive performance of the lethal COVID-19 phenotype using fibrosis classifiers across 10 cell types. **c**. Heatmaps showing: LassoCV coefficients for predicting each cell type’s fibrosis score using scores from other cell types (left); stability-selected FCI edges, with colors indicating edge existence ratios and labels denoting edge types (right). Upper triangular matrices are displayed for both heatmaps. **d**. Final causal graph derived from combined FCI results in Supplementary Fig. 10.

**Extended Data Fig 7.**
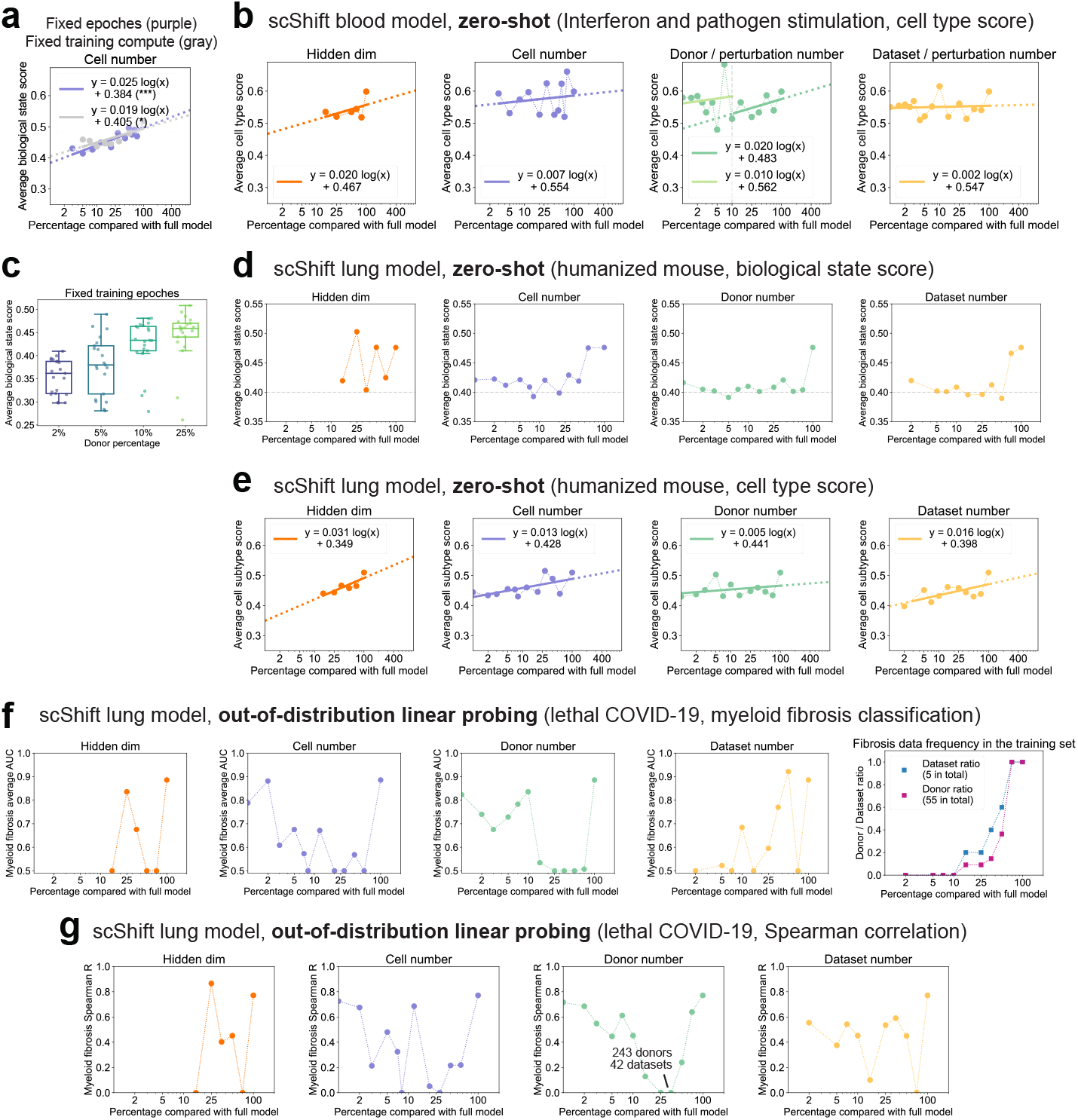
Further results on scaling analysis of scShift. **a-c**. Performance metrics for scShift blood models: **a**. average biological state scores comparing fixed epochs (50) versus fixed training compute (cell number times epochs), showing discrepancy only below 10% of full model compute; **b**. average cell type scores across unperturbed embeddings under different training settings; **c**. average biological state scores when donors/perturbations are subsampled to 2%, 5%, 10%, and 25% of the full dataset with fixed epochs (50) (n=21 per setting). **d-g**. Performance metrics for scShift lung models: **d**. average biological state scores across biological embeddings in humanized mouse myeloid cells; **e**. average cell subtype scores across unperturbed embeddings in humanized mouse myeloid cells; **f**. myeloid fibrosis classification AUCs across biological embeddings, and fibrosis dataset / donor frequency in different dataset number settings; **g**. Spearman correlation between myeloid fibrosis score and pathologic fibroblast ratios in lethal COVID-19 patients.

