## Supplementary Information for "Scaling deep identifiable models enables zero-shot characterization of single-cell biological states"

#### 1 Supplementary Note

In this part, we show the theoretical identifiability of disentangling batch-dependent and independent variations from observational distribution of single-cell gene expression under suitable assumptions. Our theoretical identifiability result serves as a foundation of the scShift model in disentangling variations and enabling comparisons across datasets.

#### 1.1 Problem setting

We consider the following simplified generative process of gene expression distributions in single cells. We define $\boldsymbol{x} \in \mathbb{R}^p$ as the observed random vector (gene expression) for each cell. Each cell belongs to a dataset (batch) that we denote as $b \in \{1, 2, ..., B\}$, which we assume to be known for each cell. $\boldsymbol{x}$ is generated with two sets of latent variables. The first set of latent variables $\boldsymbol{z} \in \mathbb{R}^{d_1}$, indicates the batch-independent latent variables, that comprises information shared across datasets such as cell types. The second set of latent variables $\boldsymbol{s} \in \mathbb{R}^{d_2}$ denotes the batch-dependent latent variables, comprising biological and batch information heterogeneous across samples and datasets. The specific generative process of $\boldsymbol{x}$ given $b$ is defined as follows:

- Generation of batch-centralized full latent variable $\boldsymbol{u}$: $\boldsymbol{u} \sim N(0, I_d)$.
- Definition of $\boldsymbol{\lambda}, \boldsymbol{z}, \boldsymbol{s}$. We define $\boldsymbol{\lambda} : \{1, ..., B\} \rightarrow \mathbb{R}^d$ as a function indicating the effect of each batch $b$ on $\boldsymbol{u}$. $\boldsymbol{\lambda}$ comprises both batch effects and global biological condition differences across datasets. We further define the index set $S(\boldsymbol{\lambda}) = \{i | \exists b \in \{1, 2, ..., B\}, \boldsymbol{\lambda}_i(b) \neq 0\} \subseteq [d]$, and $S^c(\boldsymbol{\lambda})$ as the complement set of $S(\boldsymbol{\lambda})$. We define the latent variables $\boldsymbol{z} \in \mathbb{R}^{d_1}$ and $\boldsymbol{s} \in \mathbb{R}^{d_2}$ as the components in $\boldsymbol{u} + \boldsymbol{\lambda}$ that are unaffected and affected by $\boldsymbol{\lambda}$, respectively. Specifically, we have:

$$\boldsymbol{z} := \boldsymbol{u}_{S^c(\boldsymbol{\lambda})},\ \boldsymbol{s}_b := (\boldsymbol{u} + \boldsymbol{\lambda}(b))_{S(\boldsymbol{\lambda})};\quad d_1 := |S^c(\boldsymbol{\lambda})|,\ d_2 := |S(\boldsymbol{\lambda})|. \tag{1}$$

- Generation of $\boldsymbol{x}$. Finally, we assume the gene expression of cells in batch $b$ ($\boldsymbol{x}_b$) to be sampled from the following negative binomial distribution:

$$\boldsymbol{x}_b \sim NB(f(\boldsymbol{u} + \boldsymbol{\lambda}(b)), g). \tag{2}$$

With a slight abuse of notation, the above equation may be rewritten as:

$$\boldsymbol{x}_b \sim NB(f(\boldsymbol{z}, \boldsymbol{s}), g). \tag{3}$$

Here $f : \mathbb{R}^d \rightarrow \mathbb{R}^p$ indicates the mean of the negative binomial distribution, and $g \in \mathbb{R}^p$ is a constant vector indicating the dispersion of the distribution. We assume that $f$ is injective and smooth with a smooth injective (thus $f$ is a diffeomorphism). To simplify the problem setting, we assume the constant vector $g$ is fixed and known. Our assumptions on the gene expression count distribution mostly align with those established in [1].

We denote the image of $f$ as $\mathcal{P} = \{f(u) | u \in \mathbb{R}^d\}$. Let $\theta = (\boldsymbol{\lambda}, f)$ represent the parameters of the underlying generative model. For notational convenience, we write $f_{\boldsymbol{\lambda}(b)}(\boldsymbol{u}) := f(\boldsymbol{u} + \boldsymbol{\lambda}(b))$. Let $p_{\theta,b}(x)$ represent the probability density function of gene expression vector $\boldsymbol{x} \in \mathbb{R}^p$ for a cell from batch $b$. We then define the set of all possible parameters that yield the same distribution as $p_{\theta,b}(x)$:

**Definition 1.** (Parameter family) Let $\Theta$ be the set of all possible parameters yielding the same distribution for the gene expression of a single cell from any batch $b$:

$$\Theta = \{\tilde{\theta} := (\tilde{\boldsymbol{\lambda}}, \tilde{f}) | p_{\tilde{\theta},b}(x) = p_{\theta,b}(x),\ \forall b \in \{1, 2, ..., B\};\ \tilde{f} : \mathbb{R}^d \rightarrow \mathcal{P} \text{ is smooth and invertible with a smooth inverse}\} \tag{4}$$

Our goal here is to prove that the parameters of the generative process can be correctly identified from observation distribution $p(x)$. More specifically, we aim to prove that, the identified parameter set $(\tilde{\boldsymbol{\lambda}}, \tilde{f}) \in \Theta$ and the ground truth $(\boldsymbol{\lambda}, f)$ are "equivalent" potentially upon certain transformations. We formally define the equivalence class of the parameter set as follows, which we term "disentanglement equivalence". The conditions here are very similar to the definition in [1]. The only difference is that the condition (ii) here is slightly stronger than that in [1], as we specify the function $F$ to be bijective:

**Definition 2.** (Disentanglement equivalence) Let $\tilde{\theta} = (\tilde{\boldsymbol{\lambda}}, \tilde{f}) \in \Theta$ be a set of parameters yielding the same distribution $p_{\tilde{\theta},b}(x)$ as the ground truth $\theta = (\boldsymbol{\lambda}, f)$. We say that $\tilde{\theta}$ satisfies the disentanglement equivalence relationship w.r.t. $\theta$, denoted as $\tilde{\theta} \sim \theta$, if the following three conditions hold:

$$\text{(i):} \quad \tilde{d}_1 = d_1, \tilde{d}_2 = d_2; \tag{5}$$

$$\text{(ii):} \quad \exists \text{ bijective function } F : \mathbb{R}^{d_1} \to \mathbb{R}^{d_1},\ s.t.\ \forall \rho \in \mathcal{P},\ f^{-1}(\rho)_{S^c(\boldsymbol{\lambda})} = F(\tilde{f}^{-1}(\rho)_{S^c(\tilde{\boldsymbol{\lambda}})}); \tag{6}$$

$$\text{(iii):} \quad \exists \text{ invertible } L \in \mathbb{R}^{d_2 \times d_2}, c \in \mathbb{R}^{d_2},\ s.t.\ \forall \rho \in \mathcal{P},\ f^{-1}(\rho)_{S(\boldsymbol{\lambda})} = L\tilde{f}^{-1}(\rho)_{S(\tilde{\boldsymbol{\lambda}})} + c. \tag{7}$$

*Remark* 1. Our problem setting differs from [2] in several key aspects. In our framework, the annotation label has only a "partial" effect on the latent variables—specifically affecting $\boldsymbol{s}$ but not $\boldsymbol{z}$, unlike the settings in [3, 2]. Consequently, the key results from previous works (theorem 1 in [3], theorem 23 in [2]) cannot be directly adopted since linear identifiability for $\boldsymbol{z}$ does not hold, necessitating novel theoretical approaches. Additionally, we do not assume the dimension of $\boldsymbol{z}, \boldsymbol{s}$ are correctly specified. Instead, we show that the model identifiability under our assumptions indicate correct dimensionalities of disentangled representations $\boldsymbol{z}$ and $\boldsymbol{s}$.

We next state our assumptions. In our following text, $(\boldsymbol{\lambda}, f)$ refers to the ground truth solution, a specific instance in the parameter set $\Theta$. $(\tilde{\boldsymbol{\lambda}}, \tilde{f})$ refers to **any parameters** within the parameter set $\Theta$.

#### 1.2 Assumptions

**Assumption 1.** *(MLP decoder) We define the multilayer perceptron (MLP) function family as follows:*

$$MLP = \{f | f = f_n \circ f_{n-1} ... \circ f_1, f_i(x) = \sigma(W_i x + b_i), n \in \mathbb{N}\} \tag{8}$$

*Here for simplicity, we suppose $\sigma$ to be tanh, and $W_i, b_i$ are matrices and vectors that represent weights and biases for each layer. Then we assume for the ground truth $f$ and any $(\tilde{\boldsymbol{\lambda}}, \tilde{f}) \in \Theta$, we have:*

$$\tilde{f}, f \in MLP. \tag{9}$$

**Assumption 2.** *(Batch difference) We assume that the total number of batches $B > d$. Moreover, for the ground truth $\boldsymbol{\lambda}$, we assume that $\forall i \neq j \in \{1, ..., B\}, \boldsymbol{\lambda}(b_i) \neq \boldsymbol{\lambda}(b_j)$.*

**Assumption 3.** *(Sufficient variability) There exist batches $(b_i)_0^{d_2}$ such that the $d_2$ vectors $(\boldsymbol{a}_i)_{i=1}^{d_2} = (\boldsymbol{\lambda}(b_i) - \boldsymbol{\lambda}(b_0))_{i=1}^{d_2}$ are linearly independent.*

**Assumption 4.** *(Independence between batch and latent variables) We consider any value of $u \in \mathbb{R}^d$ and $\boldsymbol{\lambda}(1), ..., \boldsymbol{\lambda}(B)$. By these we denote $\rho_b = f(u + \boldsymbol{\lambda}(b)), b \in \{1, 2, ..., B\}$. We assume that $\forall(\tilde{\boldsymbol{\lambda}}, \tilde{f}) \in \Theta, \tilde{f}^{-1}(\rho_b) - \tilde{\boldsymbol{\lambda}}(b)$ is a constant with respect to $b$. In particular, $\tilde{f}^{-1}(\rho_b)_{S^c(\hat{\boldsymbol{\lambda}})}$ is a constant with respect to $b$.*

*Remark* 2. This assumption is crucial for preventing batch-specific information from being embedded in the centralized representation $\boldsymbol{u}$, which would otherwise compromise model identifiability. It is automatically satisfied for ground truth $(\boldsymbol{\lambda}, f)$. In practice, the assumption is encouraged by enforcing independence between $\tilde{f}^{-1}(\rho_b) - \tilde{\boldsymbol{\lambda}}$ and $\tilde{\boldsymbol{\lambda}}$, where $\tilde{f}^{-1}(\rho_b) - \tilde{\boldsymbol{\lambda}}$ is approximated by the variational posterior.

**Assumption 5.** *(Non-exploding moments) We assume that the ground truth function $f$ satisfies the element-wise property: $\forall j \in \{1, 2, ..., p\}$, $\forall b \in \{1, 2, ..., B\}$, $\mathbb{E} f_{\boldsymbol{\lambda}(b)}^{2m}(\boldsymbol{u})_j = \mathcal{O}((2m)^{2m})$ for $m \in \mathbb{N}$.*

*Remark* 3. The moment assumption serves a technical purpose: it establishes conditions under which probability distributions can be uniquely determined by their moment sequences. Additional discussions can also be seen in [1].

#### 1.3 Main theoretical results

Before presenting the statement and proof of the main theorems, we introduce a lemma that will be used in the proof. The lemma aligns with Lemma 1 in the theoretical foundation of SIMVI [1]. For completeness, we restate this lemma within the context of scShift.

**Lemma 1.** *Suppose $\theta, \tilde{\theta} \in \Theta$ are two parameters that result in the same distribution of $\boldsymbol{x}_b$ for any batch b. We denote the latent variables generated with $\theta, \tilde{\theta}$ as $\boldsymbol{u}$ and $\tilde{\boldsymbol{u}}$ respectively. Then under Assumption 5, for any batch b, $f(\boldsymbol{u} + \boldsymbol{\lambda}(b)), \tilde{f}(\tilde{\boldsymbol{u}} + \tilde{\boldsymbol{\lambda}}(b)))$ have the same distribution:*

$$p_\theta(x_b) = p_{\tilde{\theta}}(x_b) \Rightarrow f_{\boldsymbol{\lambda}(b)}(\boldsymbol{u}) \overset{d}{=} \tilde{f}_{\tilde{\boldsymbol{\lambda}}(b)}(\tilde{\boldsymbol{u}}). \tag{10}$$

The proof of Lemma 1 coincides with the proof of Lemma 1 in [1]. Using the lemma, our first theorem demonstrates that the ground truth batch-dependent variable $\boldsymbol{s}$ can be represented as a linear projection of the estimated $\tilde{\boldsymbol{s}}$. This result mirrors (iii) in Definition 2, differing only in dimensionality and the invertibility of the linear projection (because (i) in Definition 2 is not yet verified).

**Theorem 1.** *(Identifiability of biological components) For any $\tilde{\theta} = (\tilde{\boldsymbol{\lambda}}, \tilde{f}) \in \Theta$, under Assumptions 1, 3, 4, 5, there exists a fixed matrix $L \in \mathbb{R}^{d_2 \times \tilde{d}_2}$ and a constant vector $c \in \mathbb{R}^{d_2}$, such that $\forall \rho \in \mathcal{P}$, $f^{-1}(\rho)_{S(\boldsymbol{\lambda})} = L\tilde{f}^{-1}(\rho)_{S(\tilde{\boldsymbol{\lambda}})} + c$.*

*Proof.* Our proof follows a similar approach to that of [3]. Applying Lemma 1, we have $f_{\boldsymbol{\lambda}(b)}(\boldsymbol{u}) \overset{d}{=} \tilde{f}_{\tilde{\boldsymbol{\lambda}}(b)}(\tilde{\boldsymbol{u}})$. Inspecting the densities at a point $\rho \in \mathcal{P}$ on both sides thus yields

$$\forall \rho \in \mathcal{P},\ \forall b \in \{1, 2, ..., B\}, \quad p(f_{\boldsymbol{\lambda}(b)}(\boldsymbol{u}) = \rho) = p(\tilde{f}_{\tilde{\boldsymbol{\lambda}}(b)}(\tilde{\boldsymbol{u}}) = \rho). \tag{11}$$

Next, we apply the change of variables formula to both sides, resulting in

$$p(f_{\boldsymbol{\lambda}(b)}(\boldsymbol{u}) = \rho) = \mathrm{vol} J_{f^{-1}}(\rho) p(\boldsymbol{u} + \boldsymbol{\lambda}(b) = f^{-1}(\rho))$$

and

$$p(\tilde{f}_{\tilde{\boldsymbol{\lambda}}(b)}(\tilde{\boldsymbol{u}}) = \rho) = \mathrm{vol} J_{\tilde{f}^{-1}}(\rho) p(\tilde{\boldsymbol{u}} + \tilde{\boldsymbol{\lambda}}(b) = \tilde{f}^{-1}(\rho)).$$

Here vol$A$ denotes the product of singular values of a matrix $A$ [3, 4]. Now because $\boldsymbol{u}, \tilde{\boldsymbol{u}} \in \mathcal{N}(0, I_d)$, we can explicitly write the log probability densities in Eq. (11). That is, $\forall \rho \in \mathcal{P}$, $\forall b \in \{1, 2, ..., B\}$,

$$\begin{aligned} &\log \mathrm{vol} J_{f^{-1}}(\rho) + \log p(\boldsymbol{u} = f^{-1}(\rho) - \boldsymbol{\lambda}(b)) = \log \mathrm{vol} J_{\tilde{f}^{-1}}(\rho) + \log p(\tilde{\boldsymbol{u}} = \tilde{f}^{-1}(\rho) - \tilde{\boldsymbol{\lambda}}(b)) \\ \Leftrightarrow &\log \mathrm{vol} J_{f^{-1}}(\rho) - ||f^{-1}(\rho) - \boldsymbol{\lambda}(b)||^2/2 = \log \mathrm{vol} J_{\tilde{f}^{-1}}(\rho) - ||\tilde{f}^{-1}(\rho) - \tilde{\boldsymbol{\lambda}}(b)||^2/2. \end{aligned} \tag{12}$$

Now we fix $\rho$ and plug each value $(b_i)_0^{d_2}$ (described in Assumption 3) in Eq. (12) leading to $d_2 + 1$ equations. We take the difference of $2, ..., d_2 + 1$th equation with the first equation, which leads to $d_2$ equations of form:

$$\begin{aligned} &\forall i \in \{1, ..., d_2\}, \\ &-||f^{-1}(\rho) - \boldsymbol{\lambda}(b_i)||^2 + ||f^{-1}(\rho) - \boldsymbol{\lambda}(b_0)||^2 = -||\tilde{f}^{-1}(\rho) - \tilde{\boldsymbol{\lambda}}(b_i)||^2 + ||\tilde{f}^{-1}(\rho) - \tilde{\boldsymbol{\lambda}}(b_0)||^2 \\ \Leftrightarrow &\langle f^{-1}(\rho), \boldsymbol{a}_i \rangle - \frac{1}{2}(||\boldsymbol{\lambda}(b_i)||^2 - ||\boldsymbol{\lambda}(b_0)||^2) = \langle \tilde{f}^{-1}(\rho), \tilde{\boldsymbol{a}}_i \rangle - \frac{1}{2}(||\tilde{\boldsymbol{\lambda}}(b_i)||^2 - ||\tilde{\boldsymbol{\lambda}}(b_0)||^2). \end{aligned} \tag{13}$$

Here $(\boldsymbol{a}_i)_{i=1}^{d_2} = (\boldsymbol{\lambda}(b_i) - \boldsymbol{\lambda}(b_0))_{i=1}^{d_2}$ as defined in Assumption 3. Correspondingly, we denoted $(\tilde{\boldsymbol{a}}_i)_{i=1}^{d_2} = (\tilde{\boldsymbol{\lambda}}(b_i) - \tilde{\boldsymbol{\lambda}}(b_0))_{i=1}^{d_2}$. Further denote $\boldsymbol{c} \in \mathbb{R}^{d_2}$ with $\boldsymbol{c}_i = \frac{1}{2}(||\boldsymbol{\lambda}(b_i)||^2 - ||\boldsymbol{\lambda}(b_0)||^2) - \frac{1}{2}(||\tilde{\boldsymbol{\lambda}}(b_i)||^2 - ||\tilde{\boldsymbol{\lambda}}(b_0)||^2)$, and matrix $A \in \mathbb{R}^{d_2 \times d} = [\boldsymbol{a}_1, ..., \boldsymbol{a}_{d_2}]^T$, $\tilde{A} \in \mathbb{R}^{d_2 \times d} = [\tilde{\boldsymbol{a}}_1, ..., \tilde{\boldsymbol{a}}_{d_2}]^T$, we have

$$Af^{-1}(\rho) = \tilde{A}\tilde{f}^{-1}(\rho) + \boldsymbol{c}. \tag{14}$$

By definition, $A$ have $d_1$ all-zero columns and the remaining $d_2$ columns have non-zero entries. That is, the submatrix $A_{S^c(\boldsymbol{\lambda})} \in \mathbb{R}^{d_1 \times d_2} \equiv 0$. Moreover, by Assumption 3, the complementary submatrix $A_{S(\boldsymbol{\lambda})} \in \mathbb{R}^{d_2 \times d_2}$ is of rank $d_2$ thus invertible. Similarly for $\tilde{A}$, the submatrix $\tilde{A}_{S^c(\tilde{\boldsymbol{\lambda}})} \in \mathbb{R}^{d_1 \times \tilde{d}_2} \equiv 0$ by definition. Altogether, Eq. (14) can be further written as

$$\begin{aligned} A_{S(\boldsymbol{\lambda})} f^{-1}(\rho)_{S(\boldsymbol{\lambda})} &= \tilde{A}_{S(\tilde{\boldsymbol{\lambda}})} \tilde{f}^{-1}(\rho)_{S(\tilde{\boldsymbol{\lambda}})} + \boldsymbol{c} \\ \Leftrightarrow f^{-1}(\rho)_{S(\boldsymbol{\lambda})} &= A^{-1}_{S(\boldsymbol{\lambda})} \tilde{A}_{S(\tilde{\boldsymbol{\lambda}})} \tilde{f}^{-1}(\rho)_{S(\tilde{\boldsymbol{\lambda}})} + A^{-1}_{S(\boldsymbol{\lambda})} \boldsymbol{c}. \end{aligned} \tag{15}$$

Denote $L := A_{S(\boldsymbol{\lambda})}^{-1} \tilde{A}_{S(\tilde{\boldsymbol{\lambda}})}$ and $c := A_{S(\boldsymbol{\lambda})}^{-1} \boldsymbol{c}$, then we have

$$f^{-1}(\rho)_{S(\boldsymbol{\lambda})} = L\tilde{f}^{-1}(\rho)_{S(\tilde{\boldsymbol{\lambda}})} + c \tag{16}$$

thus completing the proof. □

We note that without additional conditions, the matrix $L$ may lack invertibility, suggesting that the inferred batch-dependent latent variables potentially encode more information than the ground truth. We now proceed to demonstrate that an equivalence relationship (as defined in Definition 2) can be established between the "sparsest solution" and the ground truth.

**Theorem 2.** *We define the sparsest parameter $\hat{\theta} := (\hat{\boldsymbol{\lambda}}, \hat{f})$ as the element in $\Theta$ that minimizes the cardinality of $S(\hat{\boldsymbol{\lambda}})$:*

$$\hat{\theta} \in \Theta; \quad \forall \tilde{\theta} \in \Theta,\ |S(\hat{\boldsymbol{\lambda}})| \leq |S(\tilde{\boldsymbol{\lambda}})|. \tag{17}$$

*Then the equivalence relationship in Definition 2 holds for $\hat{\theta}$ and ground truth $\theta$: $\hat{\theta} \sim \theta$.*

*Proof.* The $\sim$-identifiability requires verifying conditions (i), (ii), and (iii) in Definition 2 for $\hat{\theta}$ and $\theta$. The core of our proof focuses on establishing conditions (i) and (ii), while condition (iii) is a consequence of combining (i) and Theorem 1. Our proof comprises four key steps: First, as a preparatory step, we show that the sparsity condition leads to linear independence in $\hat{\boldsymbol{\lambda}}_{S(\hat{\boldsymbol{\lambda}})}$. Second, leveraging the result from the first step, we establish condition (i), which consequently leads to condition (iii). Third, we prove a weaker version of condition (ii), i.e. the existence of a function $F$ such that $F(f^{-1}(\rho)_{S^c(\boldsymbol{\lambda})}) = \hat{f}^{-1}(\rho)_{S^c(\hat{\boldsymbol{\lambda}})}$. Finally, we prove that condition (ii) holds by combining the results from the second and third steps.

In the subsequent proof, with a slight abuse of notation, we use the form $f(z, s + \boldsymbol{\lambda}_{S(\boldsymbol{\lambda})})$ to denote $f(u + \boldsymbol{\lambda})$ in cases where this substitution enhances readability.

**Step I.** In this step, we prove that the rank of the matrix $\hat{\boldsymbol{B}} := [\hat{\boldsymbol{\lambda}}_1, ..., \hat{\boldsymbol{\lambda}}_B] \in \mathbb{R}^{d \times B}$ equals its non-all-zero-row number $|S(\hat{\boldsymbol{\lambda}})|$. If not, then $|S(\hat{\boldsymbol{\lambda}})|$ is strictly larger than the rank of $\hat{\boldsymbol{B}}$. We next show that in this case, we can construct another parameter $\hat{\theta}' = (\hat{\boldsymbol{\lambda}}', \hat{f}') \in \Theta$ that satisfies $|S(\hat{\boldsymbol{\lambda}}')| < |S(\hat{\boldsymbol{\lambda}})|$, leading to a contradiction.

We next describe the construction of $\hat{\boldsymbol{\lambda}}'$ and $\hat{f}'$. Denote the singular value decomposition of $\hat{\boldsymbol{B}}$ as $\hat{\boldsymbol{B}} = U \Lambda V^T$, we define $\hat{\boldsymbol{\lambda}}' := U^T \hat{\boldsymbol{\lambda}}$. To construct $\hat{f}'$, as $\hat{f} \in$ MLP, we consider the following expression of $\hat{f}$:

$$\hat{f} := \hat{f}_n \circ \hat{f}_{n-1} ... \circ \hat{f}_1, \quad \hat{f}_i(x) := \sigma(W_i x + b_i) \tag{18}$$

Accordingly, we define $\hat{f}'$ as follows:

$$\begin{gathered} \hat{f}' := \hat{f}_n \circ \hat{f}_{n-1} ... \circ \hat{f}'_1, \quad \hat{f}_i(x) := \sigma(W_i x + b_i)(i \geq 2), \\ f_1(x) = \sigma(W_i U x + b_i), U \in \mathbb{R}^{d \times d} \text{ is the left singular value matrix of } \hat{\boldsymbol{B}}. \end{gathered} \tag{19}$$

We next show that the constructed parameter $(\hat{\boldsymbol{\lambda}}', \hat{f}')$ is legitimate. First, as $\hat{\boldsymbol{u}}', \hat{\boldsymbol{u}} \sim \mathcal{N}(0, I_d)$, and $\hat{\boldsymbol{\lambda}}' = U^T \hat{\boldsymbol{\lambda}}$, by the orthogonality of $U$,

$$U(\hat{\boldsymbol{u}}' + \hat{\boldsymbol{\lambda}}') = U\hat{\boldsymbol{u}}' + \hat{\boldsymbol{\lambda}} \stackrel{d}{=} \hat{\boldsymbol{u}} + \hat{\boldsymbol{\lambda}},$$

thus $\hat{f}'(\hat{\boldsymbol{u}}' + \hat{\boldsymbol{\lambda}}') = \hat{f}(U(\hat{\boldsymbol{u}}' + \hat{\boldsymbol{\lambda}}')) \stackrel{d}{=} \hat{f}(\hat{\boldsymbol{u}} + \hat{\boldsymbol{\lambda}})$. Therefore, $p_{\hat{\theta}', b}(x) = p_{\hat{\theta}, b}(x)$, implying that Definition 1 holds for $\hat{\theta}'$ (all other conditions automatically hold). Second, the resulting function $\hat{f}'$ is still within the MLP family, thus Assumption 1 holds for $(\hat{\boldsymbol{\lambda}}', \hat{f}')$. Finally, by the construction of $(\hat{\boldsymbol{\lambda}}', \hat{f}')$, for any $\rho_b \in \mathcal{P}$, $\hat{f}'^{-1}(\rho_b) = U^T \hat{f}^{-1}(\rho_b)$. Therefore we have

$$\hat{f}'^{-1}(\rho_b) - \hat{\boldsymbol{\lambda}}'(b) = U^T(\hat{f}^{-1}(\rho_b) - \hat{\boldsymbol{\lambda}}(b)).$$

Since $(\hat{\boldsymbol{\lambda}}, \hat{f})$ satisfies Assumption 4, Assumption 4 is also satisfied for $(\hat{\boldsymbol{\lambda}}', \hat{f}')$. Together, $(\hat{\boldsymbol{\lambda}}', \hat{f}') \in \Theta$.

By the definition of $\hat{\boldsymbol{\lambda}}'$, $|S(\hat{\boldsymbol{\lambda}}')|$ now equals the rank of $\hat{\boldsymbol{B}}$, which is strictly smaller than $|S(\hat{\boldsymbol{\lambda}})|$. This contradicts with the definition of $(\hat{\boldsymbol{\lambda}}, \hat{f})$. Thus, the rank of $\hat{\boldsymbol{B}}$ equals $\hat{d}_2$, and the submatrix $\hat{\boldsymbol{B}}_{S(\hat{\boldsymbol{\lambda}})} \in \mathbb{R}^{\hat{d}_2 \times B}$ is of full row-rank $\hat{d}_2$.

**Step II**. We now consider Eq. (12) in Theorem 1 for $(\hat{\boldsymbol{\lambda}}, \hat{f})$: $\forall \rho \in \mathcal{P},\ \forall b \in \{1, 2, ..., B\}$,

$$\log \mathrm{vol} J_{f^{-1}}(\rho) - ||f^{-1}(\rho) - \boldsymbol{\lambda}(b)||^2 = \log \mathrm{vol} J_{\hat{f}^{-1}}(\rho) - ||\hat{f}^{-1}(\rho) - \hat{\boldsymbol{\lambda}}(b)||^2. \tag{20}$$

Because $\hat{\boldsymbol{B}}$ is of rank $\hat{d}_2$, we can select $\{b_i\}_{i=1}^{\hat{d}_2}$ such that $\{\hat{\boldsymbol{\lambda}}(b_i)\}_{i=1}^{\hat{d}_2}$ are linearly independent. We further select another $b_0 \in \{1, 2, ..., B\}$ not in $\{b_i\}_{i=1}^{\hat{d}_2}$. Now we fix $\rho$ and plug each value $\{b_i\}_{i=0}^{\hat{d}_2}$ in Eq. (20) leading to $\hat{d}_2 + 1$ equations. We subtract the first equation for $b_0$ from the remaining $\hat{d}_2$ equations, yielding $\hat{d}_2$ new equations. Analogous to Eq. (13), these $\hat{d}_2$ equations can be written together in the following matrix form:

$$\begin{aligned} Af^{-1}(\rho) = \hat{A}\hat{f}^{-1}(\rho) + \hat{\boldsymbol{c}};\ \text{where} \quad & A \in \mathbb{R}^{\hat{d}_2 \times d} := [\boldsymbol{\lambda}(b_1) - \boldsymbol{\lambda}(b_0), ..., \boldsymbol{\lambda}(b_{\hat{d}_2}) - \boldsymbol{\lambda}(b_0)]^T; \\ & \hat{A} \in \mathbb{R}^{\hat{d}_2 \times d} := [\hat{\boldsymbol{\lambda}}(b_1) - \hat{\boldsymbol{\lambda}}(b_0), ..., \hat{\boldsymbol{\lambda}}(b_{\hat{d}_2}) - \hat{\boldsymbol{\lambda}}(b_0)]^T, \quad \hat{\boldsymbol{c}} \in \mathbb{R}^{\hat{d}_2}. \end{aligned} \tag{21}$$

We next show that, under Assumption 2 and 4, the submatrix $\hat{A}_{S(\hat{\boldsymbol{\lambda}})} \in \mathbb{R}^{\hat{d}_2 \times \hat{d}_2}$ is of rank $\hat{d}_2$ thus invertible. As $\{\hat{\boldsymbol{\lambda}}(b_i)\}_{i=1}^{\hat{d}_2}$ are linearly independent, it suffices to verify that $\forall b \in \{b_i\}_{i=1}^{\hat{d}_2}, \hat{\boldsymbol{\lambda}}(b) \neq \hat{\boldsymbol{\lambda}}(b_0)$. Denote $\rho_b = f(z, s + \boldsymbol{\lambda}(b)_{S(\boldsymbol{\lambda})})$, then when $z, s$ are fixed, by assumption 2 and the injectivity of $f$, $\rho_b$ has $B$ different values for $b \in \{1, ..., B\}$. Thus by the injectivity of $\hat{f}$, $\hat{f}^{-1}(\rho_b)$ has $B$ different values. Meanwhile, by Assumption 4, $\hat{f}^{-1}(\rho_b) - \hat{\boldsymbol{\lambda}}(b)$ is a constant when $z, s$ are fixed. Therefore, $\hat{\boldsymbol{\lambda}}(b)$ must have $B$ different values for $b \in \{1, 2, ..., B\}$. Thus $\forall b \in \{1, 2, ..., B\} \neq b_0, \hat{\boldsymbol{\lambda}}(b) \neq \hat{\boldsymbol{\lambda}}(b_0)$, which establishes the desired result.

As $\hat{A}_{S(\hat{\boldsymbol{\lambda}})}$ is invertible, analogous to the derivation in Eq. (15),

$$\exists \hat{L} \in \mathbb{R}^{\hat{d}_2 \times d_2}, \hat{c} \in \mathbb{R}^{\hat{d}_2},\ s.t.\ \hat{f}^{-1}(\rho)_{S(\hat{\boldsymbol{\lambda}})} = \hat{L} f^{-1}(\rho)_{S(\boldsymbol{\lambda})} + \hat{c}. \tag{22}$$

Combining Eqs. (22, 16), the following two equations hold:

$$(I - \hat{L}L)\hat{f}^{-1}(\rho)_{S(\hat{\boldsymbol{\lambda}})} = \hat{L}c + \hat{c}; \quad (I - L\hat{L})f^{-1}(\rho)_{S(\boldsymbol{\lambda})} = L\hat{c} + c. \tag{23}$$

Because the ranges of $\hat{f}^{-1}(\rho)_{S(\hat{\boldsymbol{\lambda}})}$ and $f^{-1}(\rho)_{S(\boldsymbol{\lambda})}$ are $\mathbb{R}^{\hat{d}_2}$ and $\mathbb{R}^{d_2}$ respectively, we must have $\hat{L}L = I_{d_2}$ and $\hat{L}L = I_{\hat{d}_2}$. Therefore $\hat{L}$ is invertible, which indicates that $\hat{d}_2 = d_2$, thus $\hat{d}_1 = d_1$. Together, (i) and (iii) in Definition 2 are verified.

**Step III.** We consider arbitrary vectors $(z, s) \in (\mathbb{R}^{d_1}, \mathbb{R}^{d_2})$ and $\boldsymbol{\lambda}(1), ..., \boldsymbol{\lambda}(B)$. Denote $\rho_b = f(z, s + \boldsymbol{\lambda}_{S(\boldsymbol{\lambda})}(b))$. By the injectivity of $\hat{f}$, we can define $F_1 : \mathbb{R}^d \to \mathbb{R}^{\hat{d}_1}$ as the $S^c(\hat{\boldsymbol{\lambda}})$ components of $\hat{f}^{-1} \circ f$: $F_1 := (\hat{f}^{-1} \circ f)_{S^c(\hat{\boldsymbol{\lambda}})}$, and analogously define $F_2 : \mathbb{R}^d \to \mathbb{R}^{\hat{d}_2}$ as $F_2 := (\hat{f}^{-1} \circ f)_{S(\hat{\boldsymbol{\lambda}})}$, then we have

$$\hat{f}^{-1}(\rho_b)_{S^c(\hat{\boldsymbol{\lambda}})} = F_1(z, s + \boldsymbol{\lambda}_{S(\boldsymbol{\lambda})}(b)), \quad \hat{f}^{-1}(\rho_b)_{S(\hat{\boldsymbol{\lambda}})} = F_2(z, s + \boldsymbol{\lambda}_{S(\boldsymbol{\lambda})}(b)). \tag{24}$$

By Assumption 4, $\hat{f}^{-1}(\rho_b)_{S^c(\hat{\boldsymbol{\lambda}})}$ is constant with respect to $b$. combining with the first part of Eq. (24) we have

$$\forall z \in \mathbb{R}^{d_1},\ s \in \mathbb{R}^{d_2},\ b_i, b_j \in \{1, ..., B\}, \quad F_1(z, s + \boldsymbol{\lambda}_{S(\boldsymbol{\lambda})}(b_i)) = F_1(z, s + \boldsymbol{\lambda}_{S(\boldsymbol{\lambda})}(b_j)). \tag{25}$$

Next, we consider $d_2$ functions of the following form:

$$G_i^{z,c}(t) = F_1(z, c + t(\boldsymbol{\lambda}(b_i) - \boldsymbol{\lambda}(b_0))). \tag{26}$$

Here $\{b_i\}_{i=0}^{d_2}$ are defined in Assumption 3. By Assumption 3, $(\boldsymbol{\lambda}(b_i) - \boldsymbol{\lambda}(b_0))_{i=1}^{d_2}$ are linear independent. Then by Eq. (25), $\forall z, c$, $G_i^{z,c}(t) = G_i^{z,c}(0)$ when $t$ is integer.

Next we prove that $G_i^{z,c}(t)$ must be a constant with respect to $t \in \mathbb{R}$. By Theorem 2 in [5], we have that

$$\forall f \in \text{MLP}, \lim_{t\to\infty} f(z, c + t(\boldsymbol{\lambda}(b_i) - \boldsymbol{\lambda}(b_0))) \text{ exists.} \tag{27}$$

Because $\hat{f}^{-1}$ is smooth, following Eq. (27), the limit $\lim_{t\to\infty} \hat{f}^{-1} \circ f(z, c + t(\boldsymbol{\lambda}(b_i) - \boldsymbol{\lambda}(b_0)))_{S^c(\tilde{\boldsymbol{\lambda}})}$ exists. Since $G_i^{z,c}(t)$ is defined as the $S^c(\tilde{\boldsymbol{\lambda}})$ components of this function, its limit also exists as $t \to \infty$. Now we show that $G_i^{z,c}(t)$ is a constant function with respect to $t$ by contradiction. If there exists $t_1, t_2 > 0$ satisfying $|t_1 - t_2| < 1$, such that $G_i^{z,c}(t_2) \neq G_i^{z,c}(t_1)$, then we have

$$\begin{aligned}
&|G_i^{z,c}(t_1) - G_i^{z,c}(t_2)| \\
=&\lim_{m\in\mathbb{Z}^+\to\infty} |G_i^{z,c}(t_1 + m) - G_i^{z,c}(t_2 + m)| \\
\leq&\lim_{m\in\mathbb{Z}^+\to\infty} |G_i^{z,c}(t_1 + m) - \lim_{t\to\infty} G_i^{z,c}(t)| + |G_i^{z,c}(t_1 + m) - \lim_{t\to\infty} G_i^{z,c}(t)| \\
=&0.
\end{aligned} \tag{28}$$

This is a contradiction. Therefore, $G_i^{z,c}(t)$ is a constant function with respect to $t$.

Finally, by Assumption 3, any vector $s \in \mathbb{R}^{d_2}$ can be expressed as a weighted sum of $(\boldsymbol{\lambda}(b_i) - \boldsymbol{\lambda}(b_0))_1^{d_2}$. Noting $G_i^{z,c}(t)$ are all constants, we have that

$$\forall b \in \{1, 2, ..., B\}, \forall z \in \mathbb{R}^{d_1}, \forall s \in \mathbb{R}^{d_2}, \quad F_1(z, s + \boldsymbol{\lambda}_{S(\boldsymbol{\lambda})}(b)) = F_1(z, 0). \tag{29}$$

Now we define a function $F(z) := F_1(z, 0)$. Then by the definition of $F_1$ (Eq. (24)) and Eq. (29),

$$\hat{f}^{-1}(\rho_b)_{S^c(\hat{\boldsymbol{\lambda}})} = F_1(z, 0) = F(z). \tag{30}$$

**Step IV.** Finally we prove the function $F(z)$ (defined in step III) is bijective, thus verifying (ii) in Definition 2. Analogous to step III, define functions $\hat{F}_1 := (f^{-1} \circ \hat{f})_{S^c(\boldsymbol{\lambda})}, \hat{F}_2 := (f^{-1} \circ \hat{f})_{S(\boldsymbol{\lambda})}$, then the following equations hold:

$$\forall z \in \mathbb{R}^{d_1},\ s \in \mathbb{R}^{d_2},\ b \in \{1, ..., B\},\ \rho_b := f(z, s + \boldsymbol{\lambda}(b)_{S(\boldsymbol{\lambda})}); \tag{31}$$

$$z = \hat{F}_1(\hat{f}^{-1}(\rho_b)_{S^c(\hat{\boldsymbol{\lambda}})}, \hat{f}^{-1}(\rho_b)_{S(\hat{\boldsymbol{\lambda}})}); \tag{32}$$

$$s + \boldsymbol{\lambda}(b) = \hat{F}_2(\hat{f}^{-1}(\rho_b)_{S^c(\hat{\boldsymbol{\lambda}})}, \hat{f}^{-1}(\rho_b)_{S(\hat{\boldsymbol{\lambda}})}). \tag{33}$$

By Eq. (22) and Eq. (30), Eq. (32) can be rewritten as

$$z = \hat{F}_1(F(z), \hat{L}(s + \boldsymbol{\lambda}(b)_{S(\boldsymbol{\lambda})}) + \hat{c}). \tag{34}$$

Because Eq. (34) holds for all $s \in \mathbb{R}^{d_2},\ b \in \{1, ..., B\}$, by the invertibility of $\hat{L}$ (step II), the following equation holds for any vector $\hat{s} \in \mathbb{R}^{d_2}$:

$$z = \hat{F}_1(F(z), \hat{s}).$$

Thus define $\hat{F} : \mathbb{R}^{d_1} \to \mathbb{R}^{d_1};\ \forall q \in \mathbb{R}^{d_1},\ \hat{F}(q) = \hat{F}_1(q, 0)$, then we have

$$\forall z \in \mathbb{R}^{d_1}, \quad z = \hat{F} \circ F(z). \tag{35}$$

Therefore $F$ must be injective. Moreover, by Definition 1, the range of $\hat{z}$ is $\mathbb{R}^{d_1}$, therefore the image of $F(z)$ is $\mathbb{R}^{d_1}$. This means that $F$ is surjective. Together, $F$ is bijective, thus (ii) in Definition 2 is satisfied. Combining this with step II, we have verified all conditions (i)-(iii) in Definition 2, thus $\hat{\theta} \sim \theta$ holds.

□

#### 1.4 Implications of the theoretical results

Our theoretical results demonstrate that batch-dependent and independent variations can be identified from observational distributions when sparsity conditions and other appropriate assumptions are met. Here, we explain why these disentangled variations, particularly the batch-dependent variation, are valuable for characterizing biological states.

Different single-cell RNA sequencing datasets, even collected within same tissues, typically capture distinct biological phenomena, such as disease states, perturbation states, donor-level heterogeneity, etc. Differences of biological states across datasets leads to distinct shifts of gene expression distribution. Consequently, the ground truth batch-dependent latent variable $\boldsymbol{s} = (\boldsymbol{u} + \boldsymbol{\lambda})_{S(\boldsymbol{\lambda})}$ encompasses both pure batch effects and various biological heterogeneities in the observed data. When the pretraining dataset compendium is sufficiently large and contains all possible forms of biological heterogeneities, the ground truth batch-dependent representation $\boldsymbol{s}$ should encode all desired biological heterogeneities alongside batch effects. Thus within one batch, the variation in $\boldsymbol{s}$ or $\hat{\boldsymbol{s}}$ should only encode the biological heterogeneities of interest.

When considering multiple datasets, the batch effect may not be canceled like the single-batch case. In general, the difference of batch-dependent variation $\boldsymbol{s}$ across datasets still comprises both batch effect and biological differences. Nevertheless, with additional mild assumptions, differences within one dataset (shifts) are still comparable across datasets. Specifically, we assume that batch effects and biological differences can be linearly decomposed in the ground truth $\boldsymbol{s}$ (i.e., no interactions between batch effect and biological state):

$$\boldsymbol{s} = \boldsymbol{s}_{\text{bio}} + \boldsymbol{\lambda}_{\text{batch}}, \tag{36}$$

where $\boldsymbol{\lambda}_{\text{batch}}$ is a batch-specific constant. As an example, this assumption holds when the batch-specific term $\boldsymbol{\lambda}$ can be decomposed as

$$\boldsymbol{\lambda} = \boldsymbol{\lambda}_{\text{batch}} + \boldsymbol{\lambda}_{\text{bio}}, \tag{37}$$

leading to the following decomposition of biological and batch information in $\boldsymbol{s}$ (for cells in batch $b$):

$$\boldsymbol{s} = (\boldsymbol{u}_{S(\boldsymbol{\lambda})} + \boldsymbol{\lambda}_{\text{bio}}(b)) + \boldsymbol{\lambda}_{\text{batch}}(b). \tag{38}$$

Under this assumption, by the theoretical identifiability result, the inferred batch-dependent variation $\hat{\boldsymbol{s}}$ becomes equivalent to the biological state representation up to a linear projection and a batch-specific constant. Therefore, when computing the differences of batch-dependent variations within the same dataset, this dataset-specific constant would be canceled. Thus, these difference terms can then be compared both within and across datasets.

Notably, such comparisons are not possible with methods lacking linear identifiability. In general cases where models learn nonlinear transformations of ground truth variations of the form

$$h(\boldsymbol{z}, \boldsymbol{s}) = h(\boldsymbol{z}, \boldsymbol{s}_{\text{bio}} + \boldsymbol{\lambda}_{\text{batch}}), \tag{39}$$

even with perfectly controlled batch-independent state $z$, for any biological states $s_A, s_B$ across two batches $b_1$ and $b_2$, we have the following result in general:

$$h(z, s_A + \boldsymbol{\lambda}_{\text{batch}}(b_2)) - h(z, s_B + \boldsymbol{\lambda}_{\text{batch}}(b_2)) \neq h(z, s_A + \boldsymbol{\lambda}_{\text{batch}}(b_1)) - h(z, s_B + \boldsymbol{\lambda}_{\text{batch}}(b_1)). \tag{40}$$

The equal sign above only unconditionally holds when $h$ is a linear mapping regarding $s + \boldsymbol{\lambda}_{\text{batch}}(b)$, as achieved by our framework. Notably, the argument may also be extended to biological differences across datasets that are not of interest, such as culturing conditions, baseline condition, etc. It underscores the crucial importance of linear identifiability in enabling cross-dataset comparisons, particularly in out-of-distribution settings where biological states are imperfectly aligned.

While models lacking linear identifiability could in principle align biological states across batches through additional fine-tuning, the fine-tuning would require ground truth state annotations—making it impractical for most real-world applications—and demand extra computational resources. In contrast, scShift requires no additional fine-tuning and can readily scale biological state annotation to the atlas level through universal linear probing.

### 2 Supplementary Methods

##### Coarse-grained cell type label

We standardized cell type annotations across multiple datasets to facilitate visualization and comparative analysis. For the blood compendium, we established broad cell type categories based on the original 'cell_type' annotations, including: CD4 T cells, CD8 T cells, other T cells, B cells, NK cells, monocytes, dendritic cells, megakaryocytes, and other cell populations.

The lung compendium presents an additional need for annotation standardization due to significant imbalances in cell type proportions. We implemented a two-step annotation procedure based on the original 'cell_type' annotations: First, we consolidated less frequent cell populations by grouping all cell types beyond the top 20 most abundant categories into an "Others" category. The major cell types were then categorized based on the original 'cell_type' annotations, including T cells, B cells, monocytes, epithelial cells, pericytes, fibroblasts, and endothelial cells.

##### Benchmarking on training sets

For benchmarking the integrated representations on the training sets, we compared scShift embeddings against batch-integrated embeddings from five established methods: Harmony [6], scVI [7], scANVI [8], SCALEX [9], and scPoli [10]. Our evaluation framework was based on scIB [11], which quantifies both batch removal and biological preservation metrics using pre-defined batch labels, biological labels, and baseline pre-integrated embeddings. For the blood dataset, we implemented Harmony and SCALEX using their default settings. For both blood and lung datasets, we configured scVI to match scShift's architecture, using 100 latent components and 1000 hidden components. The scANVI model built upon the trained scVI model, using coarse-grained cell type labels for supervision and additional training of 8 epochs for blood and 5 epochs for lung data. For scPoli, we followed the tutorial-recommended settings with 50-dimensional donor embeddings (comparable to scShift's biological embedding) and trained for 18 pretraining epochs, followed by 2 epochs with the prototype loss ($\eta = 5$) for both datasets. We evaluated scIB batch removal scores (AvgBatch) and biological preservation scores (AvgBio) on a subsampled training set of 100,000 cells [11]. AvgBatch averages values of k-means NMI (Normalized Mutual Information), k-means ARI (Adjusted Rand Index), and silhouette label scores. AvgBio averages values of Silhouette batch, kBET (k-nearest-neighbor batch-effect test), graph connectivity, and PCR (Principal Component Regression) scores. We used coarse-grained cell type label as biological label and sequencing assay as the batch label for both AvgBio and AvgBatch. Additionally, for the blood dataset, we assessed cell type removal by treating coarse-grained cell type as the batch label and disease status (presence/absence) as the biological label.

##### Benchmarking on hold-out interferon and pathogen stimulation datasets

In the benchmarking, we included batch correction methods which require additional batch label inputs (Harmony, scVI [6, 7]), perturbation modeling approaches (Mixscape, contrastiveVI [12, 13]) that require ground truth perturbation label as input, and label-unaware approaches in either fine-tuned (scVI-scArches, scPoli [14, 10]) or zero-shot setting (CPA, sVAE [15, 16]). For scVI, the batch label (for the interferon dataset) and the perturbation label (for the pathogen dataset) were input into the models, covering two possible use settings. We implemented Harmony in the interferon stimulation dataset using default settings in Harmonypy. The scVI model was trained from scratch for both datasets with 2 layers and 20 latent components. Mixscape and ContrastiveVI received perturbation labels as inputs. We implemented the perturbation signature calculation in Mixscape approach on the PCA components with default neighbor numbers (20) to define its corresponding cell type representation and perturbation representation. The contrastiveVI model was implemented with suggested parameters in its Alzheimers tutorial and trained for 100 epochs with batch size 500. For scArches, we fine-tuned the scVI model trained from the assembled blood atlas for 100 epochs. For scPoli, we fine-tuned the scPoli trained from the assembled blood atlas for 50 epochs, with 40 pretraining epochs and $\eta = 10$. To enforce the scPoli model to account for stimulation conditions in the single-cell level embedding, we set the donor identity as a single index ("interferon" or "pathogen") for each dataset. We employed the SCALEX model trained on the atlas-level dataset in a zero-shot manner. We additionally trained a CPA model [15] on the assembled blood compendium with batch size 1000 for 200 epochs, and a sVAE model [16] with compatible settings with scShift (100 latent components, 2 layers) for 50 epochs. The CPA and sVAE models were applied on interferon and pathogen stimulation data in a zero-shot manner.

##### Ablation study

We additionally performed ablation study of the scShift model and tested the performance difference on the hold-out interferon stimulation dataset. The ablation study was conducted by a combination of three components: 1. sparsity regularization; 2. the denoising autoencoding training scheme; 3. the MMD regularization. We trained models with the following combinations: None (base model); 1; 1+2; 1+3; 1+2+3 (full model). All models were optimized using the same setting.

##### Unsupervised characterization of donor-level CD4 T cell biological states

For Fig. 2f, we first calculated the median CD4 T cell biological embedding of the donors with adjustment 2, and of the drugs centralized with respect to negative control (Dimethyl Sulfoxide). We applied (cosine) kernel principal component analysis on the calculated drug conditions. Then we applied the same transformation on the donor cohort. For Fig. 2g, we calculated the median CD4 T biological embedding for each perturbation from both interferon and pathogen stimulation datasets. Each condition was then centralized using the control condition. Then we calculated the principal component projection of these conditions, and applied the transformation on the donor (adjustment 1) and drug conditions (centralized by negative control). We defined the interferon stimulation score as the z-score of the sum of log normalized expressions of interferon stimulated genes (ISGs) from our selected list (*IFITM1, ISG15, LY6E, MX1, IFI44L, IFI6, IFIT1, IFIT3*), averaged among CD4 T cells for each donor/condition. For comparative analysis, we applied the same procedure used in Fig. 2g to evaluate scVI embeddings and scShift biological embeddings. To analyze the scPoli donor-level embedding, we conducted additional fine-tuning of the interferon and pathogen stimulation datasets, using their respective stimulation conditions as donor identity. To ensure comparability with the scPoli donor-level embedding (which encompasses all cell types), we computed donor-level average normalized ISG expression across all cell types as the ISG score. The subsequent analysis followed the same methodology as adopted for Fig. 2g.

##### Benchmarking on humanized mouse dataset

We further benchmarked zero-shot performances of different lung models on the chronic COVID-19 humanized mouse dataset [17]. We included a subset of methods that perform reasonably well in the blood benchmarking, covering all categories of methods: batch correction methods: scVI [7]; perturbation modeling approaches: Mixscape, contrastiveVI [12, 13]; fine-tuned label-unaware approaches: scArches-scVI, scArches-scANVI, scPoli [14, 10]. We also included the scShift full embedding in the comparison. We fine-tuned scVI and scANVI lung models on the dataset for 50 epochs. The hyperparameters of other methods were the same as the implementation on blood datasets. In benchmarking, we used the myeloid cell subset, which comprised four biological conditions (0 day post infection (dpi), 4dpi, 14dpi, 28dpi) and was annotated into four subtypes (CD14+ monocytes, CD14+CD16+ monocytes, CD16+ monocytes, macrophages). We further filtered out a small number of outliers and doublet cells (preserving Leiden clusters 0, 1, 13 in Supplementary Fig. 9c). We used the cell subtype annotation label as the label (batch) key, and the biological condition label as the batch (label) key, to evaluate unperturbed (biological) embeddings. Each individual metric was averaged and rescaled to have a minimum of 0 and maximum of 1.

##### Definition of gene sets in fibrosis analysis

To investigate gene expression patterns, we analyzed genes consistently upregulated in both lethal COVID-19 patients and at 28 days post-infection (dpi) in our humanized mouse model. The genes expressed in fewer than 100 cells were filtered out in the lethal COVID-19 dataset. The genes expressed in fewer than 50 myeloid cells were filtered out in the humanized mouse dataset. We calculated differential expression using pseudobulk log fold change (LFC) values from the lethal COVID-19 dataset and mean log-normalized differences in the humanized mouse model. We filtered out genes with mean expression smaller than 0.01 in the humanized mouse model dataset and those downregulated in lethal COVID-19 dataset (LFC$\leq$0). From the remaining genes (4174 in total), we created two lists based on the geometric average of log fold changes (LFC) between lethal COVID-19 patients and the humanized mouse model (28 dpi vs 0 dpi). Genes with a geometric average exceeding $\sqrt{0.1}$ were assigned to the "strict" list (47 genes), while those exceeding $\sqrt{0.02}$ formed the "loose" list (229 genes). We then calculated ROC curves and AUC values using scShift correlation scores (derived from lethal COVID-19 patients) and pseudobulk log p-values (derived from lethal COVID-19 versus control) to assess how well these metrics predicted genes in both lists from the full set of 4174 genes. The optimal threshold for scShift correlation scores was determined using Youden's index. After applying this threshold, we identified 39 true positive genes from the 47-gene list and 151 true positive genes from the 229-gene list (Supplementary Tables 3-4).

We further defined genes aligning with patterns of COVID-19 and fibrosis scores in the humanized mouse model dataset. Using log-normalized expression values, we calculated mean differences between timepoints: 4 dpi versus 0 dpi (4vs0), 14 dpi versus 0 dpi (14vs0), and 14 dpi versus 4 dpi (14vs4). COVID-19 pattern genes were defined by two criteria: the product of 4vs0 and 14vs0 exceeding 0.025, and the absolute value of 14vs4 less than 0.05. Similarly, fibrosis pattern genes were identified where the product of 4vs0 and 14vs4 exceeded 0.015 and the absolute value of 14vs4 was greater than 0.15. We computed the KEGG enrichment for COVID-pattern genes, fibrosis-pattern genes, and identified gene sets using EnrichR wrapped by GSEAPY [18, 19]. For the identified gene sets, we used a background gene list comprising the overlapping genes from the lethal COVID-19 and the humanized mouse dataset (7,018 genes in total).

##### Probing of fibrosis state on drug perturbations

To generate the results shown in Fig. 3k, we applied our trained scShift fibrosis classifier ("biological" setting) to myeloid cells from the drug perturbation dataset [20]. We also examined how drug-induced changes in log-normalized expression corresponded to our previously identified 47-gene (strict) and 229-gene (loose) lists for the analysis in Extended Data Fig. 6a. The resulting gene sets were smaller due to the additional filtering of genes present in the drug perturbation dataset.

### 3 Supplementary figures and tables

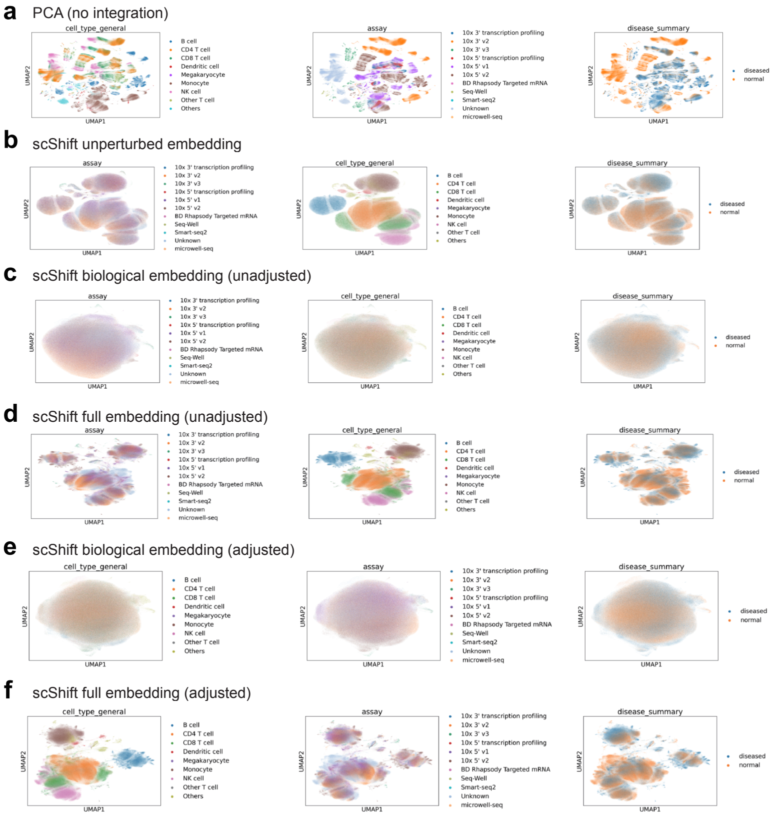

Supplementary Figure 1: UMAP visualizations of the blood training compendium using scShift embeddings and the baseline PCA representation. **a-f.** UMAP visualization using the PCA embedding (**a**), scShift unperturbed embedding (**b**), scShift unadjusted biological embedding (**c**), scShift unadjusted full embedding (**d**), scShift adjusted biological embedding (**e**), and scShift adjusted full embedding (**f**), colored by cell types, sequencing assays, and cell disease state.

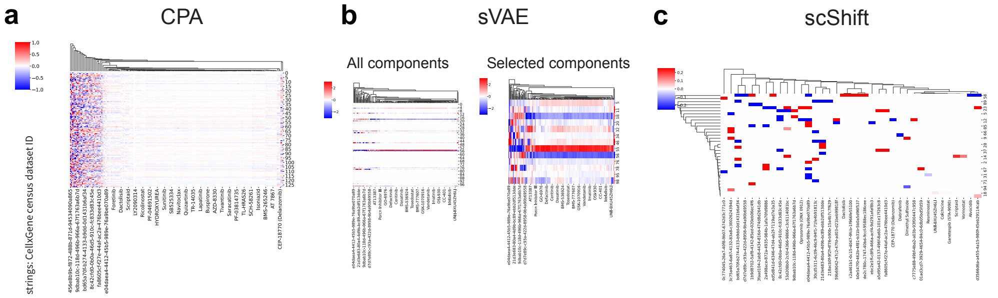

Supplementary Figure 2: Visualization of dataset-specific encodings for CPA, sVAE and scShift. **a.** CPA dataset-specific encoding. **b.** sVAE dataset-specific encoding for all components (left) and components with at least one value with absolute value larger than 0.5 (right). **c.** scShift dataset-specific encoding on biological components. CPA fails to achieve sparsity in dataset-label encoding, while sVAE's sparse components exhibit linear dependence. scShift's dataset label encoding achieves both sparsity and linear independence across components, two essential properties for model theoretical identifiability.

**a** Unperturbed embedding evaluation (batch effect removal and cell type preservation)

| Method | Bio conservation: Isolated labels | KMeans NMI | KMeans ARI | Silhouette label | cLISI | Batch correction: Silhouette batch | iLISI | KBET | Graph connectivity | Aggregate score: Batch correction | Bio conservation | Total |
|---|---|---|---|---|---|---|---|---|---|---|---|---|
| scPoli (fine-tuned) | 0.77 | 1.00 | 0.96 | 1.00 | 1.00 | 0.13 | 0.91 | 0.83 | 0.88 | 0.69 | 0.95 | 0.84 |
| Harmony (batch-aware) | 0.41 | 0.64 | 0.58 | 0.66 | 1.00 | 0.86 | 1.00 | 1.00 | 0.78 | 0.91 | 0.66 | 0.76 |
| Mixscape (perturbation-aware) | 0.46 | 0.93 | 1.00 | 0.79 | 1.00 | 0.54 | 0.49 | 0.36 | 0.62 | 0.50 | 0.84 | 0.70 |
| scShift (zero-shot) | 0.49 | 0.63 | 0.45 | 0.77 | 1.00 | 0.68 | 0.80 | 0.46 | 1.00 | 0.74 | 0.67 | 0.70 |
| scVI (batch-aware) | 0.56 | 0.34 | 0.30 | 0.58 | 1.00 | 0.84 | 0.86 | 0.46 | 1.00 | 0.79 | 0.56 | 0.65 |
| ContrastiveVI (perturbation-aware) | 0.58 | 0.89 | 0.69 | 0.73 | 1.00 | 0.64 | 0.00 | 0.15 | 0.90 | 0.42 | 0.78 | 0.64 |
| SCALEX (zero-shot) | 0.00 | 0.53 | 0.47 | 0.57 | 0.97 | 0.95 | 0.87 | 0.44 | 0.89 | 0.79 | 0.51 | 0.62 |
| sVAE (zero-shot) | 0.57 | 0.00 | 0.00 | 0.47 | 0.78 | 0.83 | 0.72 | 0.44 | 0.95 | 0.73 | 0.36 | 0.51 |
| scArches (fine-tuned) | 0.36 | 0.50 | 0.54 | 0.48 | 0.60 | 1.00 | 0.25 | 0.00 | 0.64 | 0.47 | 0.49 | 0.49 |
| CPA (zero-shot) | 1.00 | 0.02 | 0.08 | 0.00 | 0.94 | 0.00 | 0.83 | 0.66 | 0.88 | 0.59 | 0.41 | 0.48 |
| scShift (no-pretraining) | 0.48 | 0.30 | 0.15 | 0.40 | 0.00 | 0.94 | 1.00 | 0.77 | 0.00 | 0.68 | 0.26 | 0.43 |

**b** Unperturbed embedding evaluation (perturbation removal and cell type preservation)

| Method | Bio conservation: Isolated labels | KMeans NMI | KMeans ARI | Silhouette label | cLISI | Batch correction: Silhouette batch | iLISI | KBET | Graph connectivity | Aggregate score: Batch correction | Bio conservation | Total |
|---|---|---|---|---|---|---|---|---|---|---|---|---|
| scPoli (fine-tuned) | 1.00 | 1.00 | 0.96 | 1.00 | 1.00 | 0.32 | 0.25 | 0.26 | 0.88 | 0.43 | 0.99 | 0.77 |
| Mixscape (perturbation-aware) | 0.63 | 0.93 | 1.00 | 0.79 | 1.00 | 0.40 | 0.34 | 0.57 | 0.62 | 0.48 | 0.87 | 0.72 |
| ContrastiveVI (perturbation-aware) | 0.52 | 0.89 | 0.69 | 0.73 | 1.00 | 0.68 | 0.19 | 0.17 | 0.90 | 0.49 | 0.77 | 0.65 |
| scShift (zero-shot) | 0.47 | 0.63 | 0.45 | 0.77 | 1.00 | 0.64 | 0.49 | 0.30 | 1.00 | 0.61 | 0.66 | 0.64 |
| Harmony (batch-aware) | 0.49 | 0.64 | 0.58 | 0.66 | 1.00 | 0.77 | 0.07 | 0.07 | 0.78 | 0.42 | 0.68 | 0.57 |
| SCALEX (zero-shot) | 0.06 | 0.53 | 0.47 | 0.57 | 0.97 | 0.76 | 0.17 | 0.00 | 0.89 | 0.46 | 0.52 | 0.49 |
| scVI (batch-aware) | 0.34 | 0.34 | 0.30 | 0.58 | 1.00 | 0.82 | 0.00 | 0.03 | 1.00 | 0.46 | 0.51 | 0.49 |
| scArches (fine-tuned) | 0.00 | 0.50 | 0.54 | 0.48 | 0.60 | 1.00 | 0.32 | 0.01 | 0.64 | 0.49 | 0.42 | 0.45 |
| CPA (zero-shot) | 0.57 | 0.02 | 0.08 | 0.00 | 0.94 | 0.00 | 0.69 | 1.00 | 0.88 | 0.64 | 0.32 | 0.45 |
| scShift (no-pretraining) | 0.18 | 0.30 | 0.15 | 0.40 | 0.00 | 0.90 | 1.00 | 0.85 | 0.00 | 0.69 | 0.20 | 0.40 |
| sVAE (zero-shot) | 0.31 | 0.00 | 0.00 | 0.47 | 0.78 | 0.78 | 0.14 | 0.10 | 0.95 | 0.49 | 0.31 | 0.38 |

**c** Unperturbed embedding evaluation (Average of metrics in **a** and **b**)

| Method | Bio conservation: Isolated labels | KMeans NMI | KMeans ARI | Silhouette label | cLISI | Batch correction: Silhouette batch | iLISI | KBET | Graph connectivity | Aggregate score: Batch correction | Bio conservation | Total |
|---|---|---|---|---|---|---|---|---|---|---|---|---|
| scPoli (fine-tuned) | 0.95 | 1.00 | 0.96 | 1.00 | 1.00 | 0.21 | 0.67 | 0.56 | 0.88 | 0.58 | 0.98 | 0.82 |
| Mixscape (perturbation-aware) | 0.57 | 0.93 | 1.00 | 0.79 | 1.00 | 0.48 | 0.40 | 0.56 | 0.62 | 0.52 | 0.86 | 0.72 |
| scShift (zero-shot) | 0.54 | 0.63 | 0.45 | 0.77 | 1.00 | 0.66 | 0.68 | 0.42 | 1.00 | 0.69 | 0.68 | 0.68 |
| Harmony (batch-aware) | 0.48 | 0.64 | 0.58 | 0.66 | 1.00 | 0.82 | 0.67 | 0.51 | 0.78 | 0.69 | 0.67 | 0.68 |
| ContrastiveVI (perturbation-aware) | 0.63 | 0.89 | 0.69 | 0.73 | 1.00 | 0.66 | 0.00 | 0.18 | 0.90 | 0.44 | 0.79 | 0.65 |
| scVI (batch-aware) | 0.56 | 0.34 | 0.30 | 0.58 | 1.00 | 0.83 | 0.54 | 0.23 | 1.00 | 0.65 | 0.56 | 0.59 |
| SCALEX (zero-shot) | 0.00 | 0.53 | 0.47 | 0.57 | 0.97 | 0.87 | 0.60 | 0.20 | 0.89 | 0.64 | 0.51 | 0.56 |
| CPA (zero-shot) | 1.00 | 0.02 | 0.08 | 0.00 | 0.94 | 0.00 | 0.77 | 1.00 | 0.88 | 0.66 | 0.41 | 0.51 |
| scArches (fine-tuned) | 0.28 | 0.50 | 0.54 | 0.48 | 0.60 | 1.00 | 0.22 | 0.00 | 0.64 | 0.47 | 0.48 | 0.47 |
| sVAE (zero-shot) | 0.56 | 0.00 | 0.00 | 0.47 | 0.78 | 0.81 | 0.49 | 0.26 | 0.95 | 0.63 | 0.36 | 0.47 |
| scShift (no-pretraining) | 0.44 | 0.30 | 0.15 | 0.40 | 0.00 | 0.92 | 1.00 | 0.95 | 0.00 | 0.72 | 0.26 | 0.44 |

**d** Biological embedding evaluation (cell type removal and perturbation preservation)

| Method | Bio conservation: Isolated labels | KMeans NMI | KMeans ARI | Silhouette label | cLISI | Batch correction: Silhouette batch | iLISI | KBET | Graph connectivity | Aggregate score: Batch correction | Bio conservation | Total |
|---|---|---|---|---|---|---|---|---|---|---|---|---|
| scShift (zero-shot) | 1.00 | 0.82 | 0.70 | 0.84 | 0.78 | 0.96 | 0.87 | 0.64 | 1.00 | 0.87 | 0.83 | 0.84 |
| ContrastiveVI (perturbation-aware) | 0.73 | 0.53 | 0.47 | 0.88 | 0.74 | 1.00 | 1.00 | 0.96 | 1.00 | 0.99 | 0.67 | 0.80 |
| Mixscape (perturbation-aware) | 0.99 | 1.00 | 1.00 | 0.21 | 1.00 | 0.66 | 0.13 | 1.00 | 0.23 | 0.51 | 0.84 | 0.71 |
| sVAE (zero-shot) | 0.85 | 0.65 | 0.66 | 0.89 | 0.78 | 0.90 | 0.06 | 0.07 | 0.95 | 0.50 | 0.77 | 0.66 |
| SCALEX (zero-shot) | 0.86 | 0.74 | 0.68 | 0.89 | 0.73 | 0.58 | 0.02 | 0.01 | 0.93 | 0.38 | 0.78 | 0.62 |
| scVI (batch-aware) | 0.59 | 0.24 | 0.22 | 0.98 | 0.84 | 0.80 | 0.00 | 0.01 | 1.00 | 0.45 | 0.57 | 0.53 |
| scArches (fine-tuned) | 0.47 | 0.34 | 0.31 | 1.00 | 0.57 | 1.00 | 0.17 | 0.05 | 0.43 | 0.41 | 0.54 | 0.49 |
| Harmony (batch-aware) | 0.55 | 0.48 | 0.55 | 0.88 | 0.79 | 0.61 | 0.00 | 0.01 | 0.32 | 0.23 | 0.65 | 0.48 |
| scShift (no-pretraining) | 0.36 | 0.07 | 0.07 | 0.91 | 0.00 | 1.00 | 0.40 | 0.17 | 0.00 | 0.39 | 0.28 | 0.33 |
| scPoli (fine-tuned) | 0.46 | 0.18 | 0.14 | 0.69 | 0.65 | 0.00 | 0.00 | 0.00 | 0.70 | 0.17 | 0.42 | 0.32 |
| CPA (zero-shot) | 0.00 | 0.00 | 0.00 | 0.00 | 0.28 | 0.33 | 0.04 | 0.02 | 0.55 | 0.24 | 0.06 | 0.13 |

Supplementary Figure 3: Evaluating biological representation and cell type preservation performances across different methods in interferon stimulation data. **a.** scIB scores of different methods' unperturbed/integrated/background embeddings in batch removal and cell type preservation. **b.** scIB scores of different methods' unperturbed/integrated/background embeddings in perturbation removal and cell type preservation. **c.** Averaged scIB scores of different methods' unperturbed/integrated/background embeddings. **d.** scIB scores of different methods' biological/integrated/salient embeddings in cell type removal and perturbation preservation. Each metric shown is rescaled to have min 0 and max 1.

**a** Unperturbed embedding evaluation (batch effect removal and low resolution cell type preservation)

| | Bio conservation | | | | | Batch correction | | | | | Aggregate score | | |
|---|---|---|---|---|---|---|---|---|---|---|---|---|---|
| Method | Isolated labels | KMeans NMI | KMeans ARI | Silhouette label | cLISI | Silhouette batch | iLISI | KBET | Graph connectivity | PCR comparison | Batch correction | Bio conservation | Total |
| scPoli (fine-tuned) | 0.25 | 1.00 | 1.00 | 1.00 | 1.00 | 0.49 | 0.48 | 0.74 | 0.94 | 0.92 | 0.71 | 0.85 | 0.80 |
| scShift (zero-shot) | 0.48 | 0.65 | 0.53 | 0.57 | 0.97 | 0.74 | 1.00 | 0.55 | 0.95 | 0.77 | 0.80 | 0.64 | 0.70 |
| ContrastiveVI (perturbation-aware) | 0.75 | 0.64 | 0.49 | 0.43 | 0.97 | 0.78 | 0.69 | 0.63 | 1.00 | 0.45 | 0.71 | 0.66 | 0.68 |
| scVI (perturbation-aware) | 0.59 | 0.65 | 0.63 | 0.40 | 0.98 | 0.80 | 0.26 | 0.44 | 1.00 | 0.62 | 0.62 | 0.65 | 0.64 |
| Mixscape (perturbation-aware) | 0.00 | 0.90 | 0.98 | 0.59 | 0.95 | 0.50 | 0.15 | 0.33 | 0.61 | 0.46 | 0.41 | 0.69 | 0.57 |
| CPA (zero-shot) | 1.00 | 0.00 | 0.00 | 0.00 | 0.61 | 0.00 | 0.82 | 1.00 | 0.81 | 1.00 | 0.73 | 0.32 | 0.48 |
| SCALEX (zero-shot) | 0.14 | 0.58 | 0.48 | 0.44 | 0.90 | 0.67 | 0.34 | 0.08 | 0.82 | 0.00 | 0.38 | 0.51 | 0.46 |
| scShift (no-pretraining) | 0.40 | 0.30 | 0.29 | 0.19 | 0.00 | 0.95 | 0.84 | 0.64 | 0.00 | 0.82 | 0.65 | 0.24 | 0.40 |
| sVAE (zero-shot) | 0.50 | 0.04 | 0.03 | 0.27 | 0.84 | 0.70 | 0.14 | 0.24 | 0.97 | 0.00 | 0.41 | 0.34 | 0.37 |
| scArches (fine-tuned) | 0.33 | 0.24 | 0.20 | 0.24 | 0.77 | 1.00 | 0.00 | 0.00 | 0.61 | 0.00 | 0.32 | 0.36 | 0.34 |

**b** Unperturbed embedding evaluation (batch effect removal and high resolution cell type preservation)

| | Bio conservation | | | | Batch correction | | | | | Aggregate score | | |
|---|---|---|---|---|---|---|---|---|---|---|---|---|
| Method | KMeans NMI | KMeans ARI | Silhouette label | cLISI | Silhouette batch | iLISI | KBET | Graph connectivity | PCR comparison | Batch correction | Bio conservation | Total |
| scShift (zero-shot) | 0.88 | 0.59 | 0.91 | 0.88 | 0.65 | 1.00 | 0.61 | 0.88 | 0.77 | 0.78 | 0.82 | 0.80 |
| scPoli (fine-tuned) | 1.00 | 0.80 | 1.00 | 0.92 | 0.43 | 0.48 | 0.19 | 0.83 | 0.92 | 0.57 | 0.93 | 0.79 |
| scVI (perturbation-aware) | 0.94 | 0.79 | 0.86 | 1.00 | 0.78 | 0.26 | 0.37 | 0.99 | 0.62 | 0.61 | 0.90 | 0.78 |
| ContrastiveVI (perturbation-aware) | 0.85 | 0.54 | 0.86 | 0.93 | 0.71 | 0.69 | 0.49 | 1.00 | 0.45 | 0.67 | 0.79 | 0.74 |
| Mixscape (perturbation-aware) | 0.98 | 1.00 | 0.78 | 0.80 | 0.44 | 0.15 | 0.00 | 0.55 | 0.46 | 0.32 | 0.89 | 0.66 |
| SCALEX (zero-shot) | 0.82 | 0.57 | 0.80 | 0.62 | 0.64 | 0.34 | 0.05 | 0.76 | 0.00 | 0.36 | 0.70 | 0.57 |
| scShift (no-pretraining) | 0.37 | 0.32 | 0.55 | 0.00 | 0.97 | 0.84 | 0.84 | 0.00 | 0.82 | 0.69 | 0.31 | 0.46 |
| scArches (fine-tuned) | 0.40 | 0.25 | 0.78 | 0.66 | 1.00 | 0.00 | 0.00 | 0.59 | 0.00 | 0.32 | 0.52 | 0.44 |
| sVAE (zero-shot) | 0.00 | 0.00 | 0.67 | 0.71 | 0.68 | 0.14 | 0.14 | 0.88 | 0.00 | 0.37 | 0.35 | 0.35 |
| CPA (zero-shot) | 0.11 | 0.11 | 0.00 | 0.21 | 0.00 | 0.82 | 1.00 | 0.59 | 1.00 | 0.68 | 0.11 | 0.34 |

**c** Biological embedding evaluation (cell type removal and perturbation preservation)

| | Bio conservation | | | | | Batch correction | | | | | Aggregate score | | |
|---|---|---|---|---|---|---|---|---|---|---|---|---|---|
| Method | Isolated labels | KMeans NMI | KMeans ARI | Silhouette label | cLISI | Silhouette batch | iLISI | KBET | Graph connectivity | PCR comparison | Batch correction | Bio conservation | Total |
| scShift (zero-shot) | 1.00 | 0.82 | 0.91 | 0.87 | 0.81 | 0.95 | 1.00 | 1.00 | 0.92 | 1.00 | 0.97 | 0.88 | 0.92 |
| Mixscape (perturbation-aware) | 0.80 | 1.00 | 1.00 | 0.93 | 1.00 | 0.92 | 0.22 | 0.60 | 0.57 | 0.89 | 0.64 | 0.95 | 0.82 |
| ContrastiveVI (perturbation-aware) | 0.83 | 0.86 | 0.95 | 1.00 | 1.00 | 0.83 | 0.25 | 0.34 | 1.00 | 0.68 | 0.62 | 0.93 | 0.81 |
| scArches (fine-tuned) | 0.23 | 0.78 | 0.81 | 0.81 | 0.92 | 0.98 | 0.08 | 0.07 | 0.62 | 0.89 | 0.53 | 0.71 | 0.64 |
| sVAE (zero-shot) | 0.75 | 0.58 | 0.56 | 0.76 | 0.89 | 0.85 | 0.03 | 0.08 | 0.93 | 0.69 | 0.52 | 0.71 | 0.63 |
| SCALEX (zero-shot) | 0.44 | 0.25 | 0.21 | 0.60 | 0.66 | 0.54 | 0.01 | 0.02 | 0.87 | 0.00 | 0.29 | 0.43 | 0.38 |
| scVI (perturbation-aware) | 0.08 | 0.03 | 0.04 | 0.56 | 0.71 | 0.69 | 0.00 | 0.00 | 0.92 | 0.10 | 0.34 | 0.28 | 0.31 |
| scShift (no-pretraining) | 0.00 | 0.17 | 0.19 | 0.56 | 0.00 | 1.00 | 0.37 | 0.17 | 0.00 | 0.91 | 0.49 | 0.18 | 0.31 |
| CPA (zero-shot) | 0.02 | 0.00 | 0.00 | 0.00 | 0.16 | 0.85 | 0.13 | 0.28 | 0.74 | 0.87 | 0.57 | 0.04 | 0.25 |
| scPoli (fine-tuned) | 0.01 | 0.02 | 0.03 | 0.43 | 0.47 | 0.00 | 0.00 | 0.00 | 0.85 | 0.00 | 0.17 | 0.19 | 0.18 |

Supplementary Figure 4: Evaluating biological representation and cell type preservation performances across different methods in pathogen stimulation data. **a.** scIB scores of different methods' unperturbed/integrated/background embeddings in perturbation removal and low-resolution cell type preservation. **b.** scIB scores of different methods' unperturbed/integrated/background embeddings in perturbation removal and high-resolution cell type preservation. **c.** scIB scores of different methods' biological/integrated/salient embeddings in cell type removal and perturbation preservation. Each metric shown is rescaled to have min 0 and max 1.

**a** Unperturbed embedding evaluation (batch effect removal and cell type preservation)

| Method | Bio conservation | | | | | Batch correction | | | | Aggregate score | | |
|---|---|---|---|---|---|---|---|---|---|---|---|---|
| | Isolated labels | KMeans NMI | KMeans ARI | Silhouette label | cLISI | Silhouette batch | iLISI | KBET | Graph connectivity | Batch correction | Bio conservation | Total |
| **Full model** | 0.70 | 0.50 | 0.37 | 0.62 | 1.00 | 0.88 | 0.28 | 0.32 | 0.95 | 0.61 | 0.64 | 0.63 |
| **+ l0 + permutation** | 0.72 | 0.37 | 0.23 | 0.56 | 1.00 | 0.88 | 0.29 | 0.30 | 0.93 | 0.60 | 0.58 | 0.59 |
| **+ l0 + MMD** | 0.68 | 0.31 | 0.19 | 0.52 | 1.00 | 0.89 | 0.21 | 0.23 | 0.90 | 0.56 | 0.54 | 0.55 |
| **Base model** | 0.69 | 0.29 | 0.18 | 0.51 | 0.98 | 0.90 | 0.20 | 0.24 | 0.92 | 0.56 | 0.53 | 0.54 |
| **+ l0** | 0.66 | 0.25 | 0.14 | 0.51 | 0.99 | 0.88 | 0.21 | 0.22 | 0.92 | 0.56 | 0.51 | 0.53 |

**b** Unperturbed embedding evaluation (perturbation removal and cell type preservation)

| Method | Bio conservation | | | | | Batch correction | | | | Aggregate score | | |
|---|---|---|---|---|---|---|---|---|---|---|---|---|
| | Isolated labels | KMeans NMI | KMeans ARI | Silhouette label | cLISI | Silhouette batch | iLISI | KBET | Graph connectivity | Batch correction | Bio conservation | Total |
| **Full model** | 0.61 | 0.50 | 0.37 | 0.62 | 1.00 | 0.91 | 0.45 | 0.47 | 0.95 | 0.70 | 0.62 | 0.65 |
| **+ l0 + permutation** | 0.59 | 0.37 | 0.23 | 0.56 | 1.00 | 0.90 | 0.35 | 0.30 | 0.93 | 0.62 | 0.55 | 0.58 |
| **+ l0 + MMD** | 0.58 | 0.31 | 0.19 | 0.52 | 1.00 | 0.91 | 0.32 | 0.23 | 0.90 | 0.59 | 0.52 | 0.55 |
| **Base model** | 0.58 | 0.29 | 0.18 | 0.51 | 0.98 | 0.93 | 0.31 | 0.23 | 0.92 | 0.60 | 0.51 | 0.54 |
| **+ l0** | 0.57 | 0.25 | 0.14 | 0.51 | 0.99 | 0.91 | 0.35 | 0.28 | 0.92 | 0.61 | 0.49 | 0.54 |

**c** Biological embedding evaluation (cell type removal and perturbation preservation)

| Method | Bio conservation | | | | | Batch correction | | | | | Aggregate score | | |
|---|---|---|---|---|---|---|---|---|---|---|---|---|---|
| | Isolated labels | KMeans NMI | KMeans ARI | Silhouette label | cLISI | Silhouette batch | iLISI | KBET | Graph connectivity | PCR comparison | Batch correction | Bio conservation | Total |
| **Full model** | 0.51 | 0.17 | 0.09 | 0.49 | 0.68 | 0.93 | 0.24 | 0.12 | 0.61 | 0.84 | 0.55 | 0.39 | 0.45 |
| **+ l0** | 0.51 | 0.10 | 0.05 | 0.48 | 0.66 | 0.85 | 0.27 | 0.18 | 0.55 | 0.37 | 0.44 | 0.36 | 0.39 |
| **Base model** | 0.50 | 0.08 | 0.04 | 0.47 | 0.68 | 0.84 | 0.08 | 0.09 | 0.58 | 0.17 | 0.35 | 0.36 | 0.35 |
| **+ l0 + permutation** | 0.51 | 0.14 | 0.07 | 0.46 | 0.68 | 0.77 | 0.09 | 0.08 | 0.57 | 0.04 | 0.31 | 0.37 | 0.35 |
| **+ l0 + MMD** | 0.50 | 0.03 | 0.01 | 0.47 | 0.59 | 0.81 | 0.19 | 0.08 | 0.41 | 0.00 | 0.30 | 0.32 | 0.31 |

Supplementary Figure 5: Assessment of scShift ablation models in interferon stimulation data. **a.** scIB scores of different models' unperturbed embeddings in batch removal and cell type preservation. **b.** scIB scores of different models' unperturbed embeddings in perturbation removal and cell type preservation. **c.** scIB scores of different models' biological embeddings in selected cell type removal and perturbation preservation.

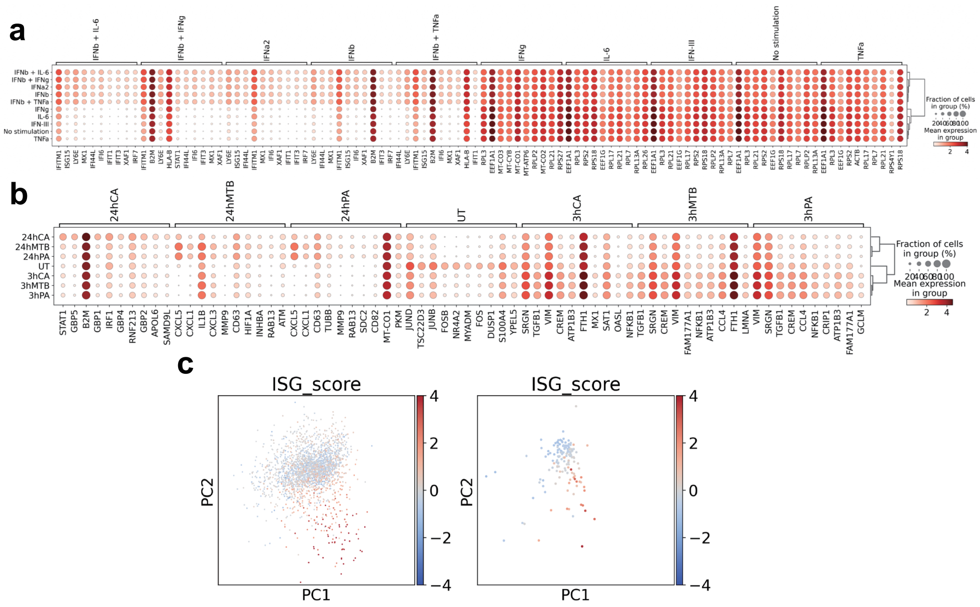

Supplementary Figure 6: Additional analysis interpreting scShift biological states. **a, b.** Dotplots showing representative differentially expressed genes in interferon (a) and pathogen (b) stimulation conditions. **c.** Visualization of donor CD4 T cells (left) and drug-perturbed CD4 T cells (right), colored by z-score of log-normalized ISG expressions. These coordinates are the same as those shown in Fig. 2g.

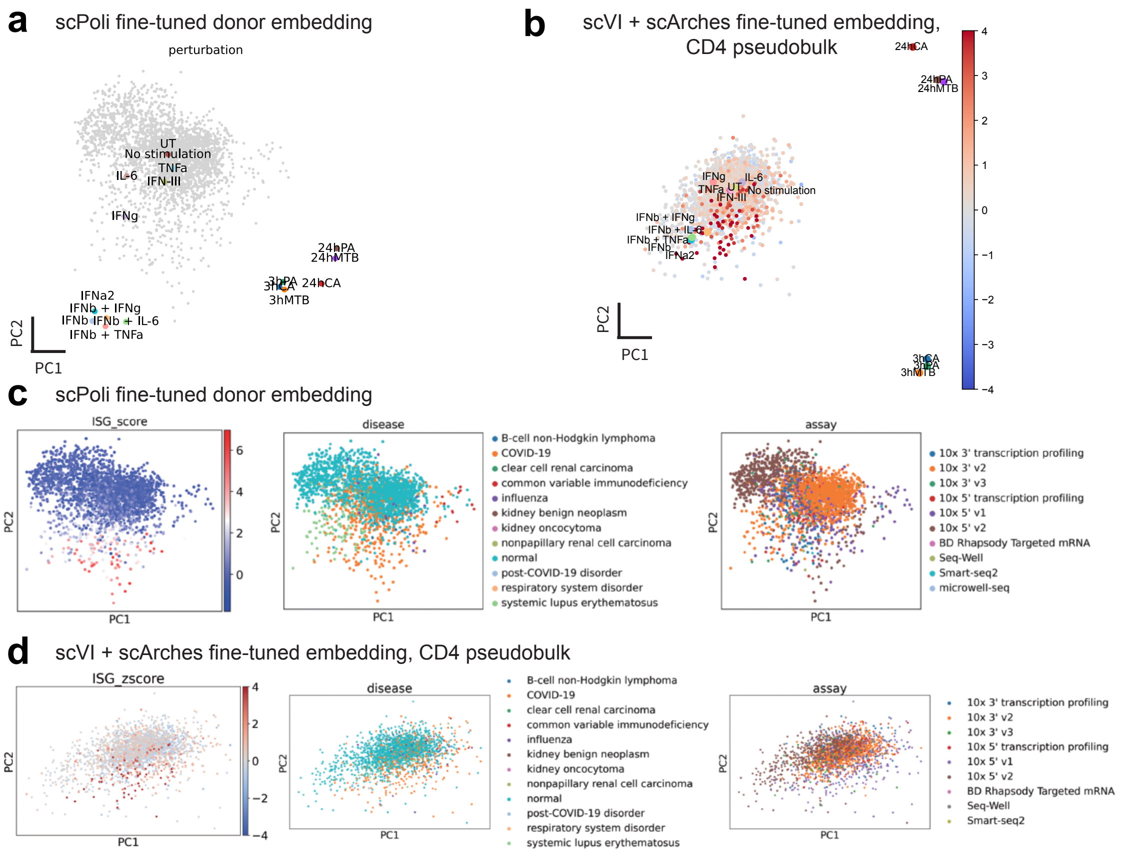

Supplementary Figure 7: Comparisons of scPoli donor-level embeddings and scVI median CD4 T cell embeddings. See Supplementary Methods for the procedure of generating these coordinates. **a.** scPoli donor-level embedding. Perturbation state were labeled in color. Grey points represent donors from the CellXGene blood atlas. **b.** scVI median CD4 T cell embedding. Perturbation state were labeled in discrete color. Other points represent donors from the CellXGene blood atlas, colored by z-score of ISG normalized expression. **c.** scPoli donor-level embedding. Points were colored by ISG score (As the scPoli donor-level embedding does not distinguish cell types, the median log-normalized ISG expression from all cell types for each donor was used), disease and assay. **d.** scVI median CD4 T cell embedding. Points were colored by ISG score (z-score of median CD4 T cell log-normalized ISG expressions), disease and assay.

**a**

| Method | Bio conservation | | | | | Batch correction | | | | | Aggregate score | | |
|---|---|---|---|---|---|---|---|---|---|---|---|---|---|
| | Isolated labels | KMeans NMI | KMeans ARI | Silhouette label | cLISI | Silhouette batch | iLISI | KBET | Graph connectivity | PCR comparison | Batch correction | Bio conservation | Total |
| **scVI (perturbation-aware)** | 0.54 | 0.15 | 0.08 | 0.52 | 0.81 | 0.96 | 0.39 | 0.58 | 0.88 | 0.65 | 0.69 | 0.42 | 0.53 |
| **scShift (zero-shot)** | 0.60 | 0.23 | 0.18 | 0.54 | 0.86 | 0.91 | 0.32 | 0.42 | 0.90 | 0.13 | 0.54 | 0.48 | 0.50 |
| **Mixscape (perturbation-aware)** | 0.56 | 0.17 | 0.06 | 0.52 | 0.79 | 0.90 | 0.29 | 0.55 | 0.74 | 0.40 | 0.58 | 0.42 | 0.48 |
| **scPoli (fine-tuned)** | 0.51 | 0.02 | 0.01 | 0.49 | 0.66 | 0.92 | 0.43 | 0.78 | 0.50 | 0.41 | 0.61 | 0.34 | 0.45 |
| **scShift-full (zero-shot)** | 0.57 | 0.08 | 0.03 | 0.52 | 0.84 | 0.88 | 0.25 | 0.25 | 0.90 | 0.00 | 0.46 | 0.41 | 0.43 |
| **scArches-scANVI (fine-tuned)** | 0.56 | 0.14 | 0.09 | 0.52 | 0.82 | 0.91 | 0.18 | 0.19 | 0.87 | 0.00 | 0.43 | 0.42 | 0.43 |
| **ContrastiveVI (perturbation-aware)** | 0.54 | 0.04 | 0.03 | 0.52 | 0.78 | 0.91 | 0.32 | 0.39 | 0.84 | 0.00 | 0.49 | 0.38 | 0.43 |
| **scPoli (zero-shot)** | 0.55 | 0.08 | 0.01 | 0.45 | 0.67 | 0.84 | 0.39 | 0.51 | 0.79 | 0.09 | 0.52 | 0.35 | 0.42 |
| **scArches-scVI (fine-tuned)** | 0.54 | 0.11 | 0.01 | 0.51 | 0.80 | 0.94 | 0.16 | 0.16 | 0.83 | 0.00 | 0.42 | 0.39 | 0.40 |

**b**

| Method | Bio conservation | | | | | Batch correction | | | | | Aggregate score | | |
|---|---|---|---|---|---|---|---|---|---|---|---|---|---|
| | Isolated labels | KMeans NMI | KMeans ARI | Silhouette label | cLISI | Silhouette batch | iLISI | KBET | Graph connectivity | PCR comparison | Batch correction | Bio conservation | Total |
| **ContrastiveVI (perturbation-aware)** | 0.55 | 0.31 | 0.30 | 0.54 | 0.90 | 0.92 | 0.26 | 0.38 | 0.94 | 0.07 | 0.51 | 0.52 | 0.52 |
| **Mixscape (perturbation-aware)** | 0.52 | 0.14 | 0.05 | 0.47 | 0.78 | 0.97 | 0.27 | 0.59 | 0.74 | 0.52 | 0.62 | 0.39 | 0.48 |
| **scPoli (fine-tuned)** | 0.52 | 0.02 | 0.01 | 0.50 | 0.57 | 0.92 | 0.34 | 0.76 | 0.64 | 0.74 | 0.68 | 0.32 | 0.47 |
| **scShift (zero-shot)** | 0.54 | 0.14 | 0.14 | 0.51 | 0.69 | 0.84 | 0.34 | 0.51 | 0.73 | 0.33 | 0.55 | 0.40 | 0.46 |
| **scArches-scVI (fine-tuned)** | 0.52 | 0.10 | 0.08 | 0.52 | 0.84 | 0.94 | 0.20 | 0.18 | 0.85 | 0.15 | 0.47 | 0.41 | 0.43 |
| **scArches-scANVI (fine-tuned)** | 0.53 | 0.11 | 0.12 | 0.52 | 0.82 | 0.91 | 0.18 | 0.22 | 0.87 | 0.00 | 0.44 | 0.42 | 0.43 |
| **scShift-full (zero-shot)** | 0.53 | 0.09 | 0.07 | 0.51 | 0.75 | 0.87 | 0.16 | 0.22 | 0.84 | 0.00 | 0.42 | 0.39 | 0.40 |
| **scPoli (zero-shot)** | 0.52 | 0.05 | 0.01 | 0.48 | 0.61 | 0.83 | 0.33 | 0.61 | 0.73 | 0.00 | 0.50 | 0.33 | 0.40 |
| **scVI (perturbation-aware)** | 0.51 | 0.02 | 0.01 | 0.50 | 0.61 | 0.92 | 0.19 | 0.23 | 0.73 | 0.04 | 0.42 | 0.33 | 0.37 |

Supplementary Figure 8: Additional results of scShift-based fibrosis analysis on the humanized mouse data. **a.** scIB scores of different methods' unperturbed/integrated/background embeddings in condition removal and myeloid subtype preservation. **b.** scIB scores of different methods' biological/integrated/salient embeddings in myeloid subtype removal and condition preservation.

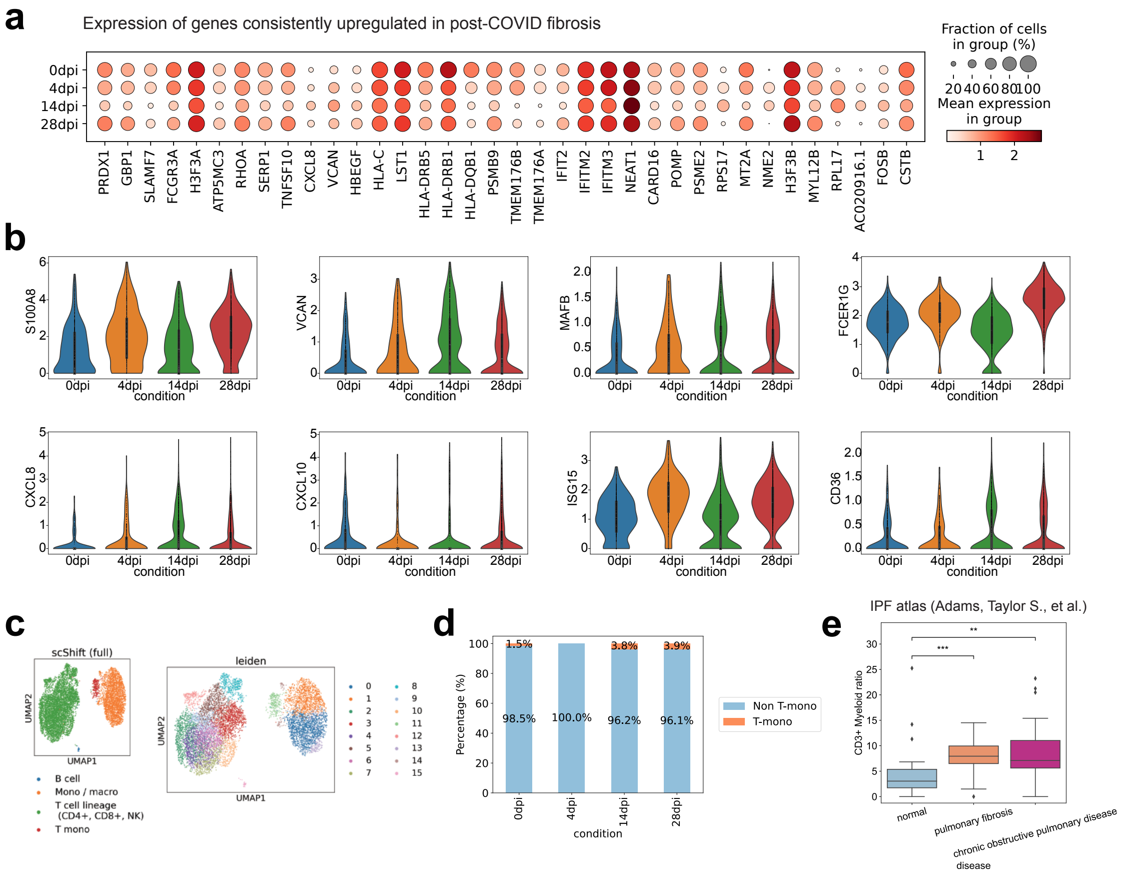

Supplementary Figure 9: Additional analysis of the humanized mouse dataset. **a.** Dotplot showing log-normalized expressions of representative genes across different conditions (days post infection, dpi) in the humanized mouse dataset. **b.** Violin plots depicting log-normalized expression of signature genes in myeloid cells from the humanized mouse dataset across different conditions (days post infection, dpi). **c.** UMAP visualization of the dataset using scShift full embedding, colored by coarse-grained cell type and Leiden clustering. **d.** Stacked bar plot showing T-mono proportions across different days post infection. **e.** Boxplot of CD3+ macrophage ratios by disease condition for each donor in IPF atlas [21] (n=28,32,18). Statistical significance was determined by two-sided Mann-Whitney-Wilcoxon test. ns: 5.00e-02<p≤1.00e+00; *: 1.00e-02<p≤5.00e-02; **: 1.00e-03<p≤1.00e-02; ***: 1.00e-04<p≤1.00e-03; ****: p≤1.00e-04.

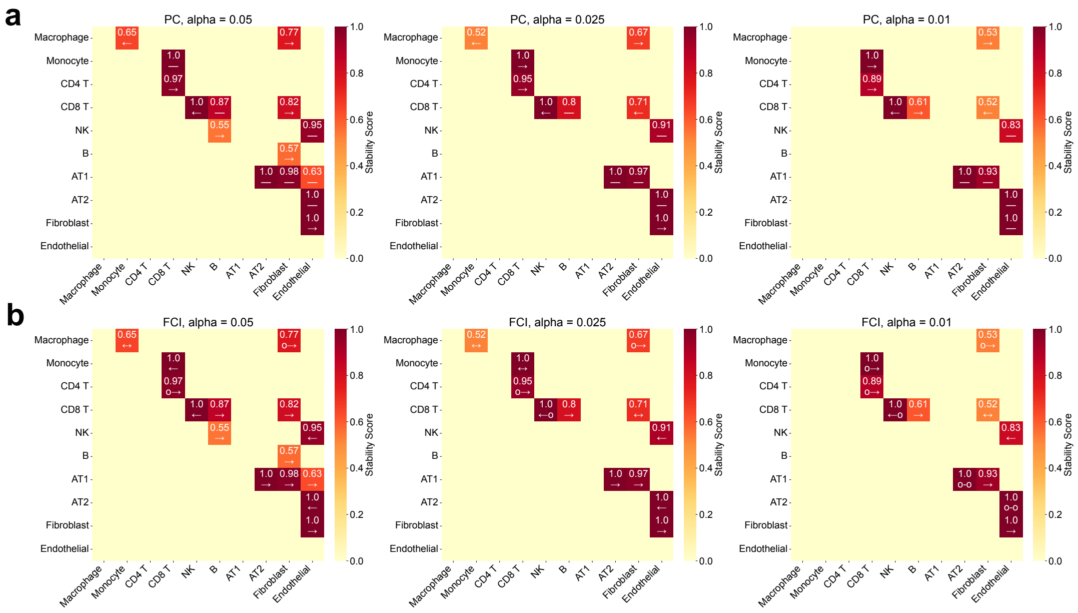

Supplementary Figure 10: Additional results from scShift-based causal modeling of fibrosis across cell types. **a.** Heatmap of stability-selected PC edges, where colors indicate edge existence ratios and labels show edge types ($\to$: row variable causes column variable; $\leftarrow$: column variable causes row variable; $-$: edge direction cannot be determined). **b.** Heatmap of stability-selected FCI edges, where colors indicate edge existence ratios and labels show edge types ($\to$: row variable causes column variable; $\leftarrow$: column variable causes row variable; o$\to$: column variable is not an ancestor of row variable; o–o: no set d-separates row and column variables; $\leftrightarrow$: latent cause exists between row and column variables). Only the upper triangular entries are shown in the heatmaps.

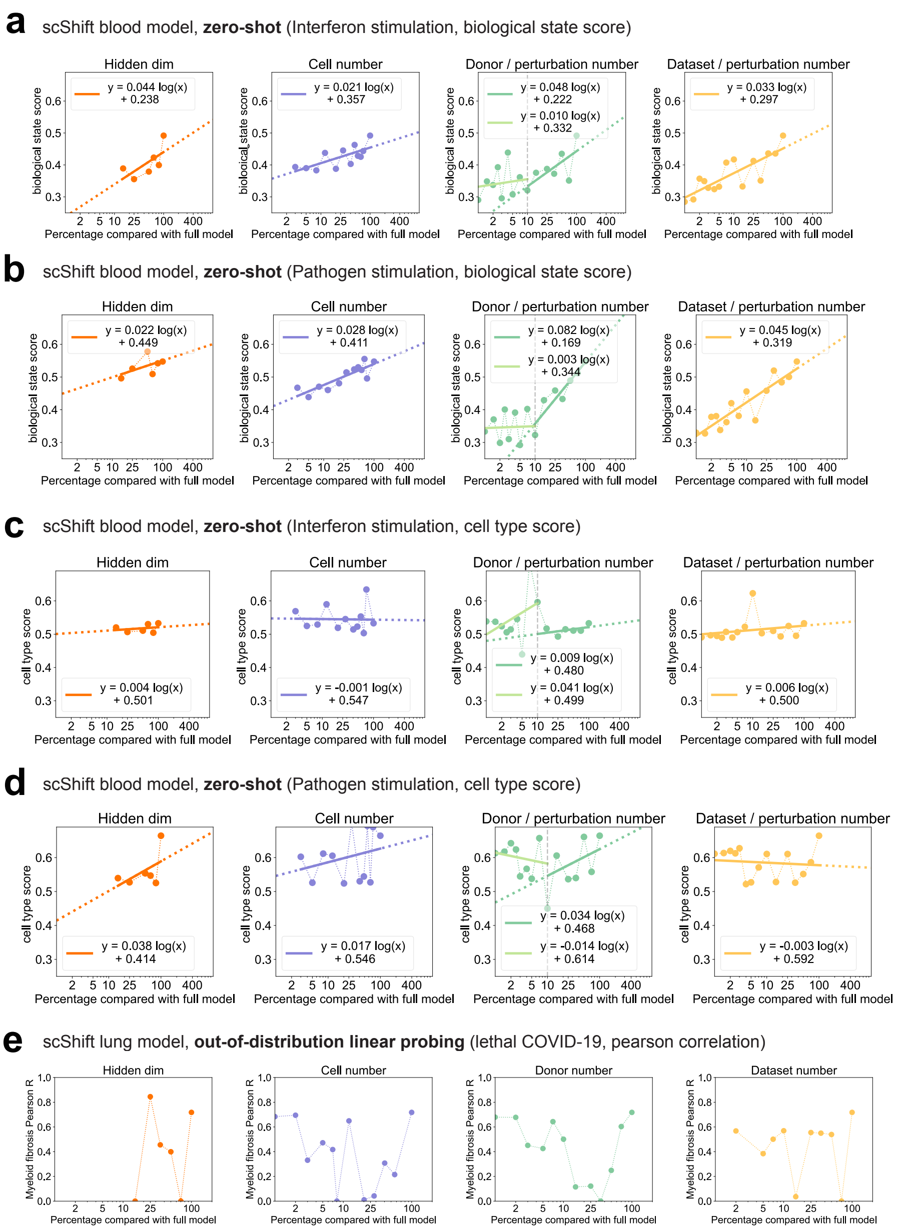

Supplementary Figure 11: Scaling analysis of zero-shot and linear probing performance across scShift blood and lung models. **a-d.** Performance metrics for scShift blood models trained under different settings: **a.** biological state scores for interferon stimulation; **b.** biological state scores for pathogen stimulation; **c.** cell type scores for interferon stimulation; **d.** cell type scores for pathogen stimulation. **e.** Pearson correlation between lethal COVID-19 patient fibrosis score and pathological fibroblast ratio for scShift lung models trained under different settings.

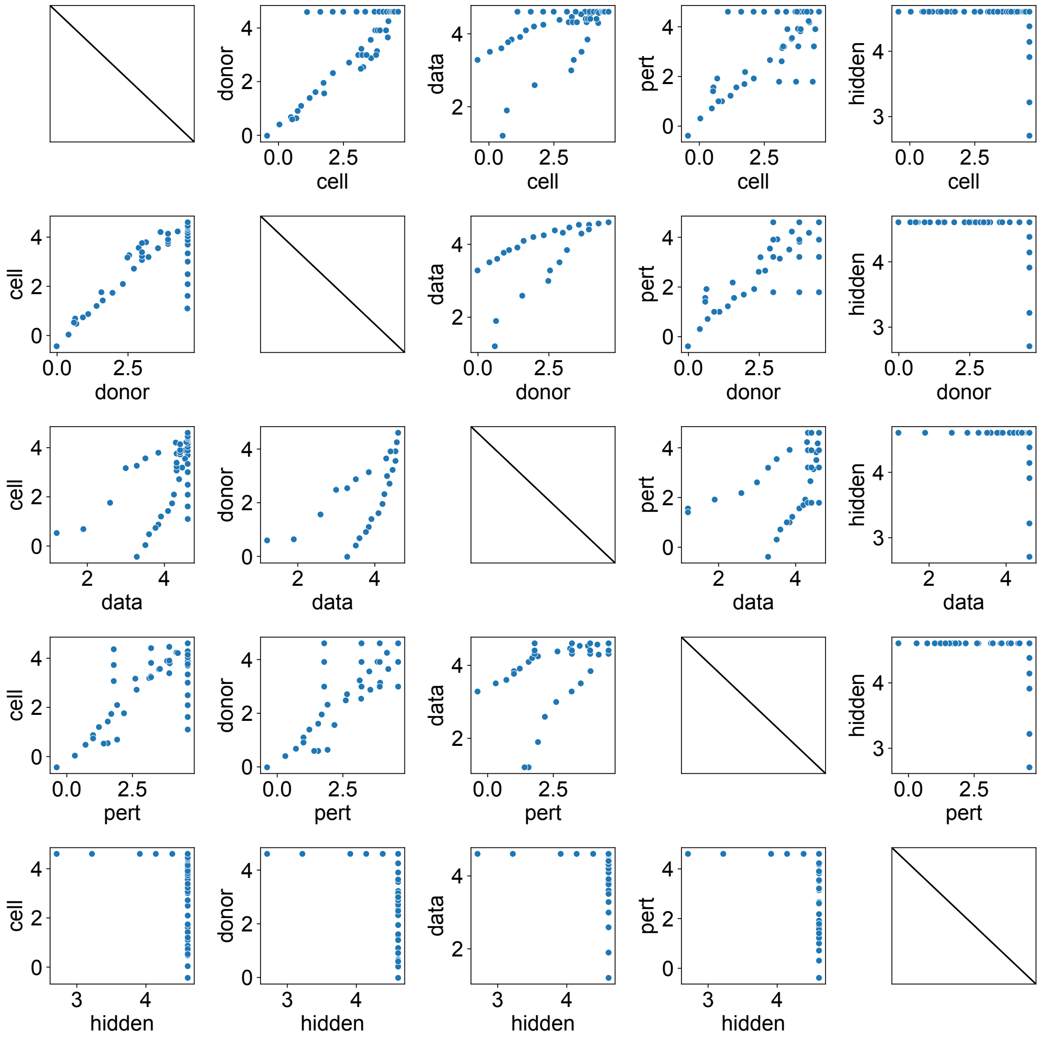

Supplementary Figure 12: Pairwise relationships between variables in scShift blood model configurations. Variables are log-normalized according to $x_{norm} = \log(100x/x_{max})$.

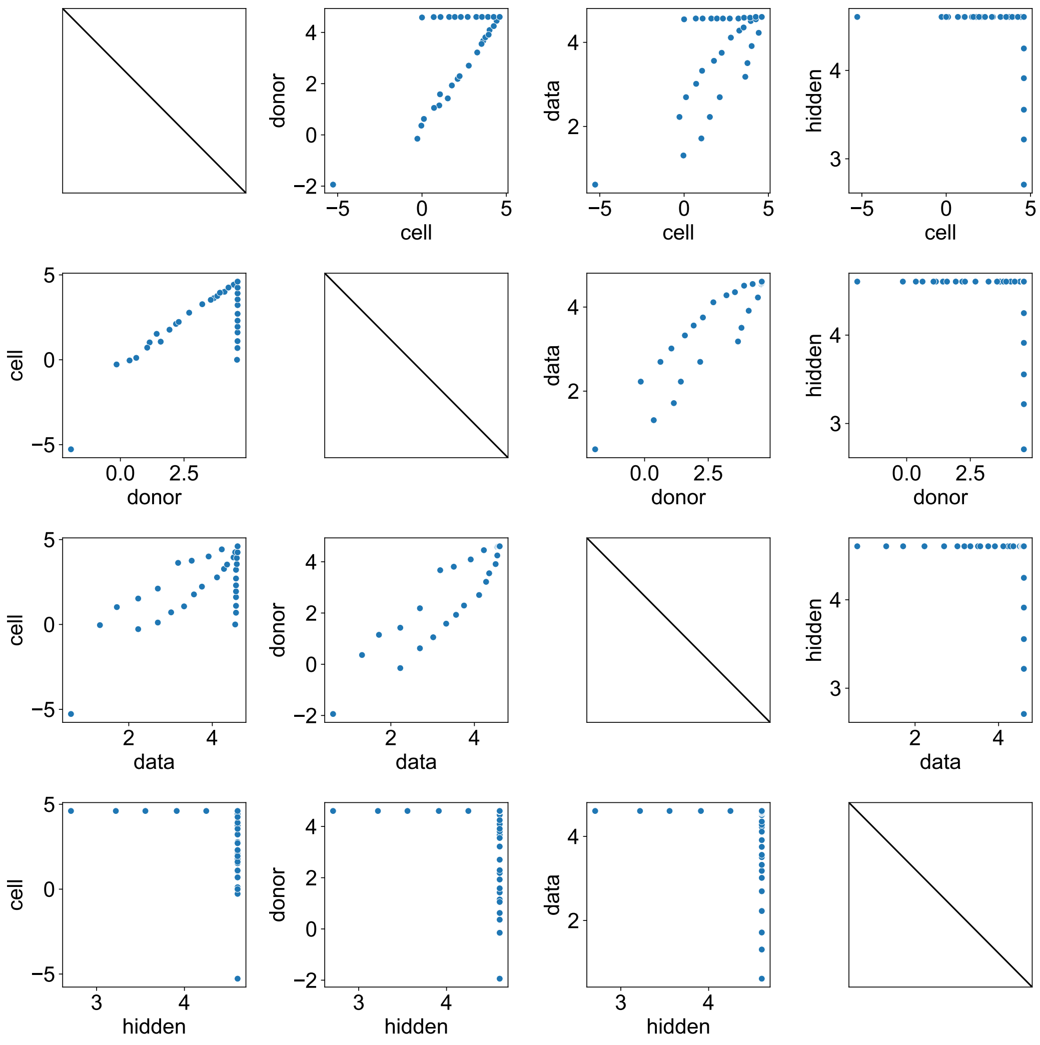

Supplementary Figure 13: Pairwise relationships between variables in scShift lung model configurations. Variables are log-normalized according to $x_{norm} = \log(100x/x_{max})$.

| Variables | Model 1 | Model 2 |
|---|---|---|
| Constant | 0.158* | 0.194*** |
| | (0.086) | (0.067) |
| | [0.074] | [0.006] |
| $\log(\text{Cell} \times 100 / \text{Cell}_{max})$ | 0.020*** | 0.020*** |
| | (0.005) | (0.005) |
| | [0.001] | [0.000] |
| $\log(\text{Donor} \times 100 / \text{Donor}_{max})$ | 0.019* | 0.023*** |
| | (0.010) | (0.006) |
| | [0.077] | [0.000] |
| $\log(\text{Data} \times 100 / \text{Data}_{max})$ | 0.017 | – |
| | (0.018) | – |
| | [0.360] | – |
| $\log(\text{Pert} \times 100 / \text{Pert}_{max})$ | -0.004 | – |
| | (0.006) | – |
| | [0.509] | – |
| $\log(\text{Hidden} \times 100 / \text{Hidden}_{max})$ | 0.023* | 0.024** |
| | (0.012) | (0.012) |
| | [0.060] | [0.043] |
| *Model Statistics* | | |
| Observations | 41 | 41 |
| R-squared | 0.522 | 0.503 |
| Adjusted R-squared | 0.454 | 0.463 |
| F-statistic | 7.656*** | 12.50*** |
| AIC | -176.9 | -179.2 |
| BIC | -166.6 | -172.4 |

Notes: Standard errors in parentheses. P-values in square brackets.
Significance levels: *** $p<0.01$, ** $p<0.05$, * $p<0.1$

Supplementary Table 1: Scaling law model selection for scShift blood models (donor count > 258) based on average zero-shot biological state scores. Two models are evaluated: Model 1 incorporates all parameters (cells, donors, datasets, perturbations, hidden dimensions), while Model 2 considers only cells, donors, and hidden dimensions. $\text{Cell}_{max}$ = 1,240,090; $\text{Donor}_{max}$ = 2,538; $\text{Data}_{max}$ = 30; $\text{Pert}_{max}$ = 147; $\text{Hidden}_{max}$ = 100.
Model 1: $y = \text{Constant} + \log(\text{Cell} \times 100 / \text{Cell}_{max}) + \log(\text{Donor} \times 100 / \text{Donor}_{max}) + \log(\text{Data} \times 100 / \text{Data}_{max}) + \log(\text{Pert} \times 100 / \text{Pert}_{max}) + \log(\text{Hidden} \times 100 / \text{Hidden}_{max})$;
Model 2: $y = \text{Constant} + \log(\text{Cell} \times 100 / \text{Cell}_{max}) + \log(\text{Donor} \times 100 / \text{Donor}_{max}) + \log(\text{Hidden} \times 100 / \text{Hidden}_{max})$.

| | Interferon stimulation data | | Pathogen stimulation data | |
|---|---|---|---|---|
| Variables | Model 1 | Model 2 | Model 1 | Model 2 |
| Constant | 0.046 | 0.125 | 0.270** | 0.264** |
| | (0.106) | (0.083) | (0.130) | (0.100) |
| | [0.665] | [0.141] | [0.046] | [0.012] |
| log(Cell × 100 / $Cell_{max}$) | 0.015** | 0.015** | 0.026*** | 0.026*** |
| | (0.006) | (0.006) | (0.008) | (0.008) |
| | [0.029] | [0.031] | [0.003] | [0.002] |
| log(Donor × 100 / $Donor_{max}$) | 0.004 | 0.014* | 0.034** | 0.031*** |
| | (0.013) | (0.007) | (0.016) | (0.009) |
| | [0.767] | [0.060] | [0.037] | [0.001] |
| log(Data × 100 / $Data_{max}$) | 0.031 | – | 0.003 | – |
| | (0.022) | – | (0.027) | – |
| | [0.174] | – | [0.922] | – |
| log(Pert × 100 / $Pert_{max}$) | -0.003 | – | -0.005 | – |
| | (0.007) | – | (0.009) | – |
| | [0.717] | – | [0.564] | – |
| log(Hidden × 100 / $Hidden_{max}$) | 0.041*** | 0.042*** | 0.005 | 0.007 |
| | (0.014) | (0.014) | (0.018) | (0.017) |
| | [0.008] | [0.006] | [0.776] | [0.711] |
| *Model Statistics* | | | | |
| Observations | 41 | 41 | 41 | 41 |
| R-squared | 0.323 | 0.282 | 0.471 | 0.466 |
| Adjusted R-squared | 0.227 | 0.224 | 0.396 | 0.423 |
| F-statistic | 3.344** | 4.839*** | 6.244*** | 10.76*** |
| AIC | -159.9 | -161.5 | -142.7 | -146.3 |
| BIC | -149.6 | -154.6 | -132.5 | -139.5 |

Notes: Standard errors in parentheses. P-values in square brackets.

Significance levels: *** p<0.01, ** p<0.05, * p<0.1

Supplementary Table 2: Scaling law model selection for scShift blood models (donor count > 258) based on biological state scores in interferon and pathogen stimulation datasets. Models 1 and 2 follow the definitions provided in Supplementary Table 1.

| Gene List Name | Genes |
|---|---|
| COVID pattern genes | *PARK7, C1QA, C1QB, ATP6V0B, RPS27, KRTCAP2, VAMP5, IGKC, COX5B, DBI, UBXN4, ARPC2, SEPTIN2, BRK1, ARL6IP5, RPL24, RNF7, CXCL10, HINT1, ANKHD1, SERPINB1, CLIC1, HLA-DQA1, PSMB8, HLA-DMA, DNPH1, PSMB1, MRPS24, LINC-PINT, GIMAP4, RAB2A, ZNF706, VPS28, TXN, OPTN, GSTO1, LSP1, SLC15A3, PLAAT4, FKBP2, RESF1, PTGES3, NACA, NAP1L1, NDUFA12, LCP1, RPS29, RPL36AL, COX5A, RPS15A, PYCARD, CIAO2B, PSMB6, C17orf49, GPS2, UBB, SNHG29, CCL2, CCL3, NDUFV2, C18orf32, CYB5A, MICOS13, EIF3K, RPS19, PPM1N, SELENOW, MAFB, ATP5PF, BID, IGLC2, IGLC3, SAT1, RPL36A, SEPTIN6, SSR4, RPS4Y1, DDX3Y, MT-CO1, MINOS1, FAM129A, SEPT2, COL4A3BP, SEPT7, XIST, SEPT6, C8orf59, RARRES3, FAM45A, KIAA1551, FAM96B, SEPT9, C19orf24, C19orf70* |
| Fibrosis pattern genes | *PRDX1, GBP1, SLAMF7, FCGR3A, H3F3A, ATP5MC3, RHOA, SERP1, TNFSF10, CXCL8, VCAN, HBEGF, HLA-C, LST1, HLA-DRB5, HLA-DRB1, HLA-DQB1, PSMB9, TMEM176B, TMEM176A, IFIT2, IFITM2, IFITM3, NEAT1, CARD16, POMP, PSME2, RPS17, MT2A, NME2, H3F3B, MYL12B, RPL17, AC020916.1, FOSB, CSTB* |
| The 47-gene list | *PTP4A2, C1orf162, CD48, FCER1G, PPP1CB, PCBP1, CAPG, VAMP5, H1FX, TN-FSF10, SLC39A8, VCAN, HSP90AB1, MARCKS, ACTB, PSMA2, PPIA, CD36, SRGN, IFIT1, CTSD, MS4A6A, TMEM179B, CD63, ATP5F1B, UBC, KCTD12, NPC2, B2M, ANXA2, PSMA4, BCL2A1, MAF, ACTG1, MYL12A, MYL12B, CHMP1B, BST2, PLAUR, FTL, LILRA6, FKBP1A, MAFB, MRPS6, TYMP, SAT1, RBM3* |
| The 39-gene list (filtered by scShift) | *PTP4A2, CD48, FCER1G, PCBP1, CAPG, VAMP5, H1FX, TNFSF10, SLC39A8, VCAN, HSP90AB1, MARCKS, ACTB, PSMA2, PPIA, CD36, SRGN, IFIT1, CTSD, TMEM179B, CD63, ATP5F1B, UBC, KCTD12, NPC2, B2M, ANXA2, PSMA4, BCL2A1, ACTG1, MYL12A, MYL12B, BST2, FTL, LILRA6, FKBP1A, MAFB, TYMP, SAT1* |
| The 229-gene list | *VAMP3, PGD, SDHB, CDC42, EPHB2, RPL11, SH3BGRL3, IFI6, PEF1, PTP4A2, MACF1, PRDX1, UQCRH, TMEM59, PDE4B, GBP4, TMEM167B, C1orf162, CTSS, S100A4, ADAR, MNDA, CD48, FCER1G, CREG1, IER5, GLUL, SRP9, IRF2BP2, ID2, LAPTM4A, SF3B6, FOSL2, PPP1CB, RPS27A, MXD1, PCBP1, MOB1A, MTHFD2, TMSB10, CAPG, VAMP8, VAMP5, MAP3K2, KYNU, CLK1, SUMO1, IDH1, ARL4C, FLNB, EIF4E3, CD47, PARP9, PARP14, H1FX, CDV3, ATP1B3, RNF13, SELENOT, RAP2B, MFSD1, TNFSF10, ATP13A3, TMEM33, EREG, AREG, HERC5, H2AFZ, SLC39A8, UBE2D3, C4orf33, ACSL1, LPCAT1, SNX18, BTF3, TMEM167A, VCAN, ARRDC3, ST8SIA4, TGFBI, SSR1, C6orf62, HLA-F, AIF1, SRSF3, CDKN1A, VEGFA, HSP90AB1, CD2AP, ELOVL5, PNRC1, MARCKS, SMPDL3A, ARMT1, DYNLT1, QKI, PSMB1, ACTB, RAC1, PSMA2, PPIA, PURB, SNHG15, FGL2, CD36, BRI3, IFRD1, CTSB, ASAH1, PLEKHA2, SDCBP, LY96, PDP1, TP53INP1, COX6C, KLF10, FAM49B, EEF1D, C9orf72, CHMP5, CARD19, ATP5F1C, CREM, HNRNPF, SRGN, PPIF, GHITM, IFIT1, PIK3AP1, PRDX3, CTSD, ADM, RNF141, LMO2, MS4A6A, TMEM179B, NEAT1, CFL1, UCP2, TMEM123, CASP1, MLF2, SLC2A3, SLC38A2, TMBIM6, METTL7A, DAZAP2, PFDN5, CD63, STAT2, ATP5F1B, NACA, GNS, BTG1, PLXNC1, OAS1, UBC, POMP, KCTD12, PCID2, ARHGEF40, DAD1, SLC7A7, PSME1, PSME2, ARF6, GCH1, HIF1A, ZFP36L1, NPC2, GPR65, HSP90AA1, PDIA3, B2M, SQOR, DMXL2, ANXA2, PSTPIP1, PSMA4, BCL2A1, MEFV, ARL6IP1, PRKCB, TENT4B, MAF, PFN1, EIF4A1, UBB, WSB1, PRKAR1A, SUMO2, ACTG1, METRNL, MYL12A, MYL12B, CHMP1B, C18orf32, RPL17, MCOLN1, BST2, PLAUR, NAPA, FTL, LILRA6, LILRA5, LILRB1, LILRB4, FKBP1A, PRNP, RIN2, MAFB, ZFAS1, PRELID3B, SAMSN1, ATP5PO, MRPS6, ETS2, TCN2, HMOX1, APOL6, TSPO, TYMP, TMSB4X, SAT1, CYBB, ATP6AP2, RBM3, PGK1, ATP6AP1* |

Supplementary Table 3: Gene lists identified in the analysis.

| Gene List Name | Genes |
|---|---|
| The 151-gene list (filtered by scShift) | *PGD, CDC42, RPL11, SH3BGRL3, IFI6, PEF1, PTP4A2, PRDX1, GBP4, CTSS, S100A4, MNDA, CD48, FCER1G, IER5, GLUL, SRP9, IRF2BP2, ID2, LAPTM4A, SF3B6, RPS27A, MXD1, PCBP1, MOB1A, MTHFD2, TMSB10, CAPG, VAMP8, VAMP5, SUMO1, FLNB, EIF4E3, PARP14, H1FX, CDV3, SELENOT, RAP2B, TNFSF10, ATP13A3, TMEM33, H2AFZ, SLC39A8, BTF3, TMEM167A, VCAN, SSR1, HLA-F, AIF1, SRSF3, CDKN1A, HSP90AB1, ELOVL5, PNRC1, MARCKS, ARMT1, PSMB1, ACTB, RAC1, PSMA2, PPIA, FGL2, CD36, BRI3, CTSB, SDCBP, COX6C, KLF10, EEF1D, CHMP5, HNRNPF, SRGN, PPIF, GHITM, IFIT1, PIK3AP1, PRDX3, CTSD, ADM, RNF141, LMO2, TMEM179B, CFL1, UCP2, TMEM123, CASP1, MLF2, SLC2A3, TMBIM6, METTL7A, DAZAP2, PFDN5, CD63, ATP5F1B, NACA, GNS, BTG1, PLXNC1, OAS1, UBC, POMP, KCTD12, DAD1, SLC7A7, PSME1, PSME2, ARF6, GCH1, ZFP36L1, NPC2, HSP90AA1, PDIA3, B2M, SQOR, ANXA2, PSMA4, BCL2A1, ARL6IP1, PRKCB, PFN1, EIF4A1, UBB, PRKAR1A, SUMO2, ACTG1, METRNL, MYL12A, MYL12B, C18orf32, RPL17, BST2, FTL, LILRA6, LILRA5, LILRB1, LILRB4, FKBP1A, PRNP, MAFB, PRELID3B, ATP5PO, HMOX1, APOL6, TSPO, TYMP, TMSB4X, SAT1, CYBB, ATP6AP2, PGK1, ATP6AP1* |

Supplementary Table 4: Gene lists identified in the analysis (continued).
